# The Amot/Integrin protein complex transmits mechanical forces required for vascular expansion

**DOI:** 10.1101/2021.01.14.426638

**Authors:** Yuanyuan Zhang, Yumeng Zhang, Sumako Kameishi, Giuseppina Barutello, Yujuan Zheng, Nicholas P. Tobin, John Nicosia, Katharina Hennig, David Kung-Chun Chiu, Martial Balland, Thomas H. Barker, Federica Cavallo, Lars Holmgren

**Affiliations:** Department of Oncology and Pathology, Bioclinicum, Karolinska Institutet, SE-171 64 Stockholm, Sweden; Department of Molecular Biotechnology and Health Sciences, Molecular Biotechnology Center, University of Turin, 10126 Turin, Italy; Wallace H. Coulter Department of Biomedical Engineering, Georgia Institute of Technology, Atlanta, Georgia 30332, United States; Department of Biomedical Engineering, University of Virginia, Charlottesville, Virginia 22904, United States; Laboratoire Interdisciplinaire de Physique, Université Joseph Fourier (Grenoble 1), 38402 Saint Martin d’Hères Cedex, France

## Abstract

Vascular development is a complex multistep process involving the coordination of cellular functions such as migration, proliferation and differentiation. Understanding the underlying mechanisms of these processes is of importance due to involvement of vessel expansion in various pathologies. How mechanical forces generated by cells and transmission of these physical forces control vascular development is poorly understood. Using an endothelial-specific genetic model in mice, we show that deletion of the scaffold protein, Angiomotin (Amot), inhibits migration and expansion of physical and pathological vascular network. We further show that Amot is required for tip cell migration and the extension of cellular filopodia. Exploiting *in vivo* and *in vitro* molecular approaches, we show that Amot binds talin and is essential for relaying forces between fibronectin and the cytoskeleton. Finally, we provide evidence that Amot is a novel component of the endothelial integrin adhesome and propose that Amot integrates spatial cues from the extra-cellular matrix in order to form a functional vascular network.

## Introduction

Most, if not all, organs are dependent on the formation of an organized blood circulatory network to function properly. The mechanisms that control the ingrowth of vessels have garnered much attention given its importance in the pathogenesis of various disease conditions such as ischemia, cancer and diabetes (Folkman, 1995, Potente, Gerhardt et al., 2011). The *de novo* blood vessel formation, angiogenesis, is triggered by morphogenic gradients that promote orderly collective migration of endothelial cells (ECs) that give rise to a blood circulatory network (Eilken & Adams, 2010, Gerhardt & Betsholtz, 2005). Endothelial tip cells guide the expanding vessel network and probe the environment by extending cellular protrusions, known as filopodia (Gerhardt, Golding et al., 2003). These filopodia not only detect and respond to chemotactic factors such as VEGF-A and axon guidance factors but also mechanically interact with the surrounding extra-cellular matrix (ECM). The ECM exerts mechanical control on the endothelium via binding to integrins that relay forces to the cellular cytoskeleton (Hynes, 2002, Schwartz, 2010). Here, integrins form a tension-dependent link between fibrillar ECM and the cytoskeleton that is critical for translating of mechanical forces into biochemical signals (Geiger, Bershadsky et al., 2001). The actual tension is generated by actomyosin contraction, which provides tractional force required for migration together with the actual stiffness of the ECM (Baneyx, Baugh et al., 2002, Zhang, Magnusson et al., 1997, Zhong, Chrzanowska-Wodnicka et al., 1998). A major component of the developing blood vessels is Fibronectin (Fn) (Pankov & Yamada, 2002). The Fn-binding endothelial integrins α5β1 and αvβ3/β5 integrin receptors have emerged as possible targets for anti-angiogenic therapies. However, the mechanisms of these molecular pathways controlling angiogenesis are complex. Inactivation of integrin α5 and αv during mouse development causes death *in utero* due to vascular defects (Murphy, Begum et al., 2015). Although highly expressed in tumor endothelium, conditional deletion of Fn, α5β1 and αvβ3/β5 does not inhibit tumor expansion, which suggests the up-regulation of alternative compensatory pathways. Indeed, analysis of the integrin adhesome has identified over 180 proteins, with some of those regulated by actin tension (Schiller, Friedel et al., 2011, Zaidel-Bar & Geiger, 2010). Further insight on how mechanical forces are transduced may provide perspective on how these pathways could be targeted in a more efficient way.

In this report we have studied the role of Angiomotin (Amot) in the transmission of force during EC migration. We have previously reported the identification of the Amot protein family, which consist of three members Amot, Amot-Like 1 (AmotL1) and AmotL2 (Bratt, Wilson et al., 2002). All three members are scaffold proteins characterized by a coiled-coil and WW domains as well as a PDZ-binding motif, and each of them has two identified isoforms, including p130/p80 Amot, p100/p90 AmotL1 and p100/p60 AmotL2. They are of interest as they bind and localize membrane receptors with polarity proteins complexes and regulate the Hippo pathway and the cytoskeleton. We have recently shown that Amot, AmotL1 and AmotL2 exert distinct functions in blood vessel development, using zebrafish and conditional knockout mice (Hultin, Zheng et al., 2014, Zheng, Zhang et al., 2016). AmotL2 was demonstrated to bind to the VE-cadherin complex and is required for formation of radial actin filaments. These filaments transduce force at the cellular junctions and inactivation in mouse and zebrafish embryos abrogates aortic expansion. A similar function of AmotL2 is found in epithelial cells where AmotL2 binds to E-cadherin, organizes radial actin filaments, and is required for blastocyst hatching in pre-implantation embryos (Hildebrand, Hultin et al., 2017). Amot was the original protein of this family and is essential for EC migration. In this study we investigated the role of Amot in angiogenesis, setting up its involvement in the transmission of force in ECs. We used an endothelial specific mouse knockout model to study post-natal angiogenesis. We present evidence that Amot is a part of the endothelial integrin adhesome and is essential for force transmission between ECM, integrins and the actin cytoskeleton.

## Results

### Amot is expressed in blood vessels of human placenta as well as human tumors

We have previously reported a preferential expression of Amot in endothelial cells during physiological angiogenesis of zebrafish and mouse (Aase, Ernkvist et al., 2007). In addition, recent publications have implicated Amot as a regulator of the Hippo pathway in epithelial cells *in vitro*, suggesting a role in carcinogenesis (Zeng, Ortiz et al., 2017). To clarify this apparent discrepancy, we analyzed the Amot expression pattern in normal and cancerous human tissues. Amot expression pattern was mapped in both normal and pathological tissue sections with immunoaffinity-purified Amot antibodies, which do not cross-react with AmotL1 or AmotL2 (Levchenko, Aase et al., 2003). Amot positivity was restricted to blood vessels of third trimester placenta but not detectable in adult endometrium (**Figure 1A** and **1B**). Furthermore, protein expression was not detectable in areas outside blood vessels in adult tissues such as pancreas, brain, breast, prostate, and colon (**Figure 1C**). In addition to blood vessels of angiogenic tissues, Amot staining was also detected in blood vessels of the adult epididymis and in podocytes of the kidney glomeruli (**Figure S1A** and **S1B**). Analysis of human cancers of different origins showed an apparent upregulation of Amot in tumor blood vessels and stroma whereas the adjacent tumor cells were negative (**Figure 1D** and **Figure S1C-F**).

**Figure 1.**
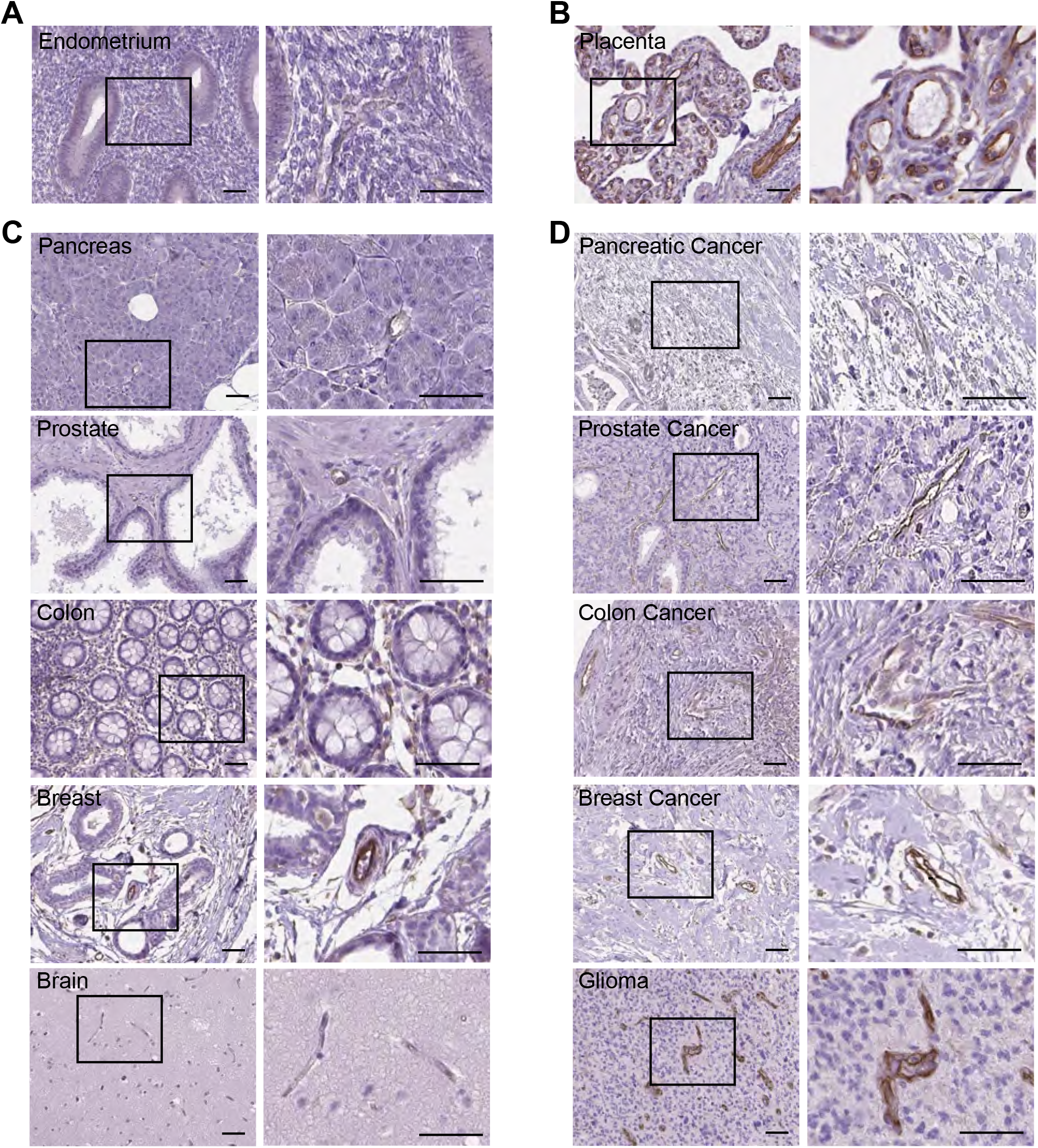
Localization of Amot in blood vessels of human tissues. (A-D) Immunohistochemical (IHC) localization of Amot protein. Rabbit Amot antibodies were generated against the c-terminal 20 a.a. and immunoaffinity purified. The IHC of patient samples was performed in collaboration with human protein atlas (Uhlen, Fagerberg et al., 2015). Detailed information of patient samples is listed in **Table S5**. Human paraffin sections of endometrium (A) and placenta (B) were stained with Amot antibodies positive signal shown in brown. (C) and (D) shows representative images of Amot protein expression in normal, including pancreas, prostate, colon, breast and brain, as well as corresponding cancerous tissues. Right panels show magnification of boxed areas. Scale bars, 50 μm.

### Amot is essential for endothelial tip cell migration

The mouse retina is an established model for the studies of physiological angiogenesis, as it becomes vascularized in a highly reproducible manner over the first 10 days after birth (Pitulescu, Schmidt et al., 2010, Uemura, Kusuhara et al., 2006). We performed whole-mount immunofluorescence (IF) staining to detect Amot expression at neonatal (day 6) and in adult retinas. Amot was ubiquitously expressed in retinal vessels at P6, whereas no signal above background was detected in retinas of adult mice (**Figure 2A**). We further analyzed Amot steady-state protein levels in whole retinas at postnatal day 6 (P6) as well as retinas from adult mice. The results showed that Amot expression decreased by 60 % from P6 when compared to the adult stage (**Figure 2B** and **C**).

**Figure 2.**
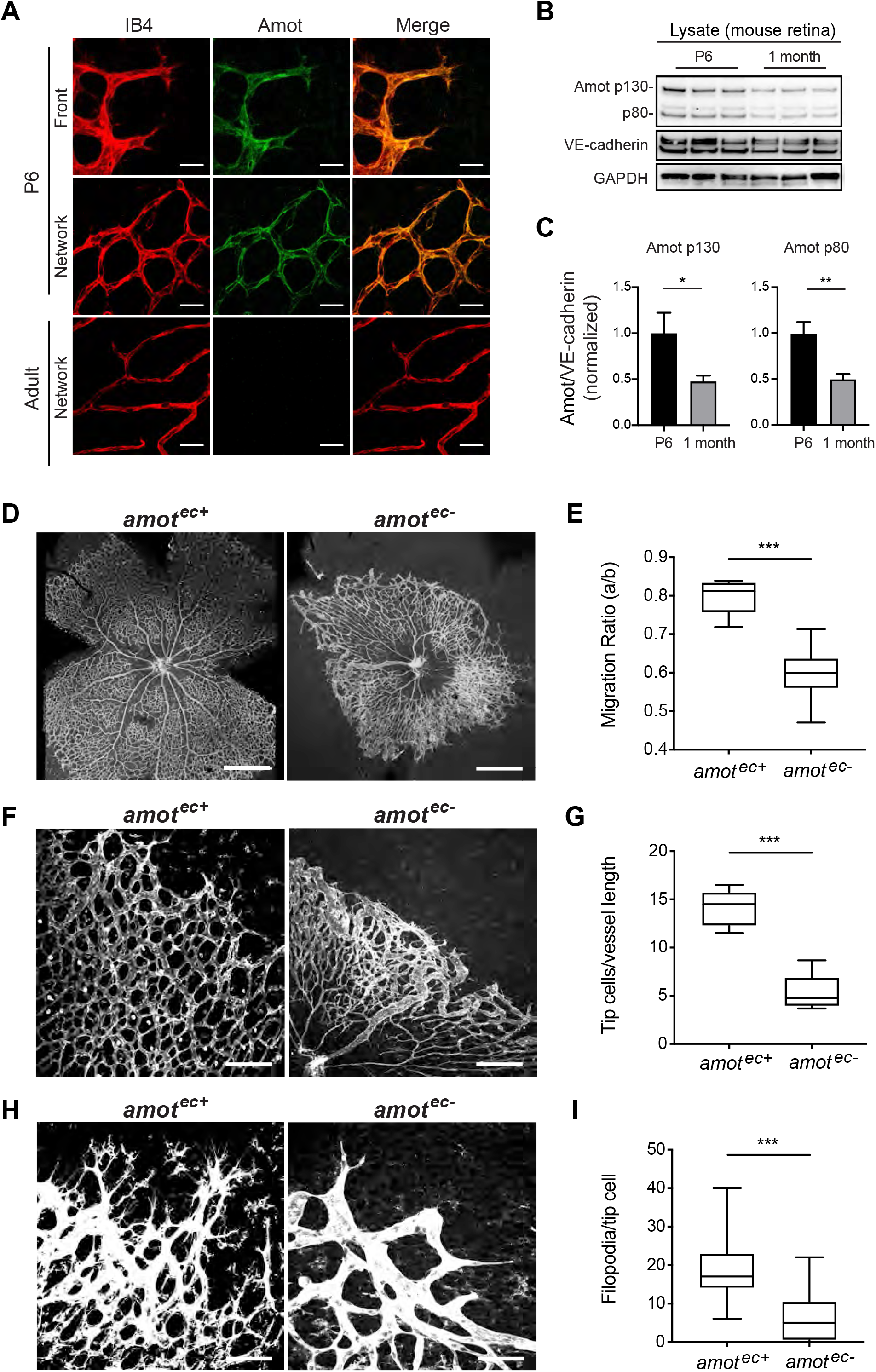
Amot is required for radial vessel expansion. (A) Whole-mount fluorescence staining of a wild-type mouse retina with isolectin B4 (IB4, in red) and Amot (in green). The leading front of the vasculature front and capillary networks at P6 are displayed in upper and middle panels. Capillaries in adult retina (24 weeks) are shown in the lower panel. Scale bars, 25 μm (upper/middle panel) and 50 μm (lower panel). (B) Western blot (WB) analysis of Amot (p130 and p80), together with VE-cadherin and GAPDH in mouse retinas at P6 and adult stage (four weeks). Retinas were mixed together for protein extraction and analyzed for each time point (n=3). (C) Quantification of Amot intensity on WB bands by ImageJ. Amot intensity was normalized to VE-cadherin levels in the same mouse. (D), (F), and (H) Overview images of *amot*^*ec*+^ and *amot*^*ec*−^ mouse retinas visualized by IB4 staining. Scale bars, (D) 1 mm, (F) 100 μm, (H) 50 μm. (E) Radial vessel expansion was quantified as the length of expanded vessel divided by the full length of retina (measured from the optic nerve to the edge). Five *amot*^*ec*+^ mice and six *amot*^*ec*−^ mice were included in the quantification. (G) Box diagram showing the quantification of the tip cell number per vessel length at the sprouting front in *amot*^*ec*+^ (n=5) and *amot*^*ec*−^ (n=6) retinas. At least four images per retina were analyzed. (I) Box plot illustrating the decrease in the number of filopodia per tip cell after *amot* ablation. 51 tip cells were quantified in *amot*^*ec*+^ mice and 61 in *amot*^*ec*−^ mice. \*\*\**P* < 0.001.

Our previous studies showed that 75% of Amot conventional knockout mice die before embryonic day 12 due to severe vascular defects (Aase et al., 2007). In order to study the role of Amot in normal and tumor angiogenesis in postnatal mice, we employed an inducible genetic deletion approach to silence Amot expression in ECs. To inactivate Amot (localized on the X-chromosome) in a cell-type specific fashion, we crossed mice containing loxP sites flanking exon 4 and 5 with Cdh5 (PAC)^CreERT2^ transgenic mice (hereafter abbreviated *amot*^e*c*+^ or *amot*^*ec*−^) (Pitulescu et al., 2010). This model allowed efficient tamoxifen-inducible recombinase expression in ECs. Tamoxifen-induced recombination was verified by genomic PCR analysis and monitored at a cellular level by the activation of the ROSA26-EYFP reporter (**Figure S2A**).

To investigate the role of Amot during postnatal retinal angiogenesis, we induced Cre recombinase activity by injecting tamoxifen at P1-3. The retinas were harvested at P6, dissected and analyzed by whole-mount staining. Inactivation of *amot* resulted in inhibition of radial vessel expansion, which appeared non-symmetrical and partially collapsed (**Figure 2D** and **E**). Endothelial tip cells guide vessel expansion and sense environmental cues via the extension of filopodia. Quantification of tip cells showed a significant decrease in the number of tip cells per certain length of vascular outline in *amot*^*ec*−^ retinas, compared with *amot*^*ec*+^ retinas (**Figure 2F** and **G**). In addition, *amot*^*ec*−^ tip cells exhibited a four-fold reduction in the number of filopodial extensions (**Figure 2H** and **I**). By contrast, deletion of *amot* in adult mice did not result in detectable aberrations of the retinal vasculature (**Figure S2B**).

### Amot is required for endothelial tip cell positioning

Our data show that Amot is expressed in both tip cells and capillaries of developing retinas. However, Amot depletion seemed to primarily affect EC tip cells. In order to further investigate this apparent discrepancy, we made use of the fact that injection of decreasing doses of tamoxifen results in lower recombination frequencies. In order to trace and follow positioning of individual EC, we crossed *amot*^*ec*+^ mice with ROSA26-EYFP Cre-reporter mice (Srinivas, Watanabe et al., 2001). Cre activity was monitored by estimating the ratio of YFP-positivity in EC cells in tip cell position vs the capillary bed (**Figure 3A** and **S3A**). In *amot*^*ec*+^/ROSA26-EYFP retinas, YFP^+^ tip and capillary EC showed a strong linear relationship and therefore no difference in recombination at the two distinct sites (**Figure 3B**). We then went on to compare the recombination frequency of *amot*^*ec*+^ and *amot*^*ec*−^ tip cells (**Figure 3C** and **S3B-D**). At 50%-85% recombination the ratio of YFP^+^ tip cells vs capillaries in *amot*^*ec*+^ EC was normalized to 1 and compared to the ratio of *amot*^*ec*−^. Analysis of the *amot*^*ec*−^ retinas showed that the comparable ratio was significantly lower (0.3) indicating a decreased ability to occupy tip cell position (**Figure 3C**). Higher recombination frequencies (85%-100%) forced *amot*^*ec*−^ tip cells to localize at the leading edge, however, these EC tip cells exhibited severe defects (**Figure 3D** and **E**).

**Figure 3.**
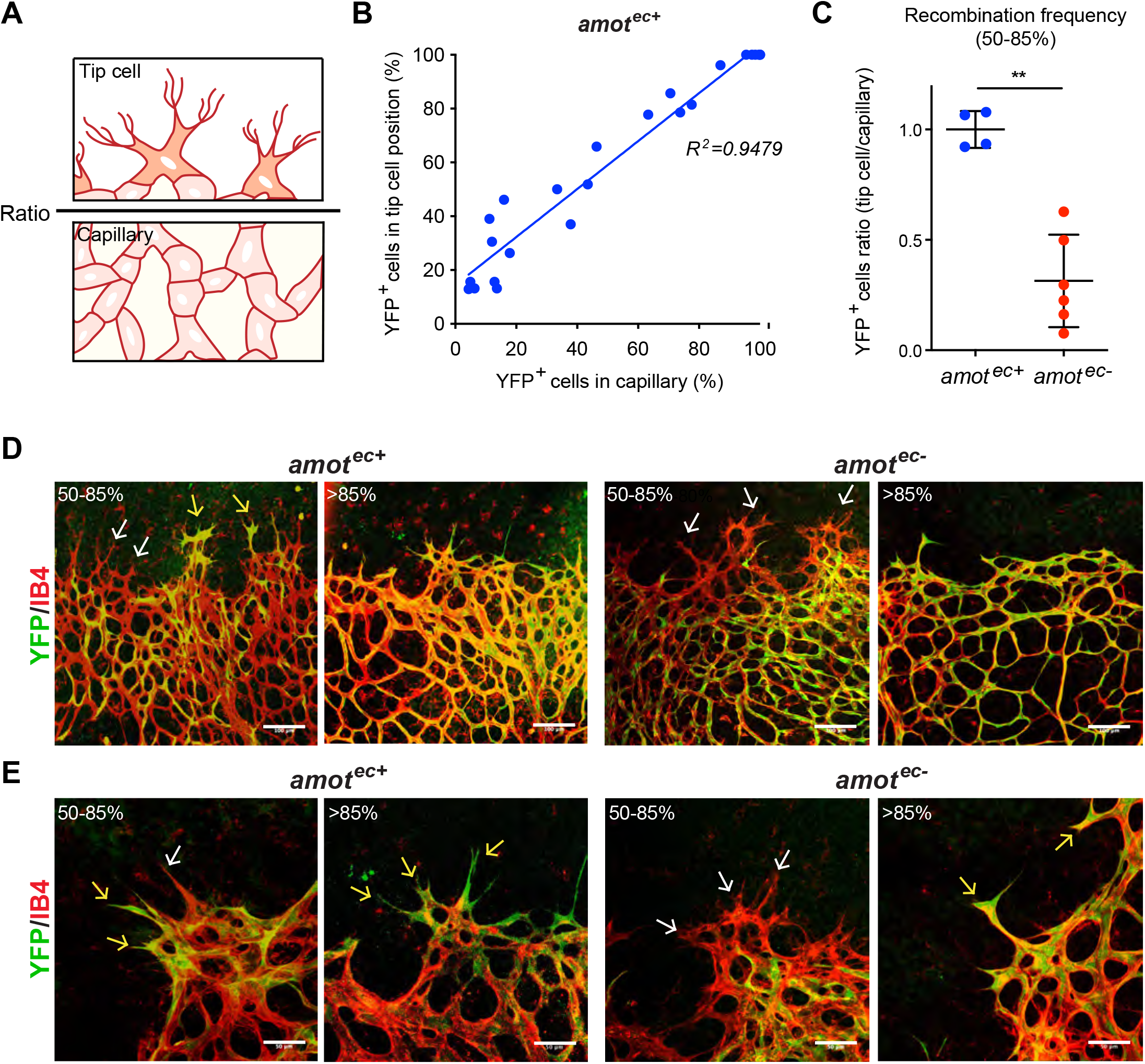
Amot is required for EC tip cell positioning. (A) Schematic indicating tip cell and capillary areas where quantification of recombination was performed. The “ratio” represents the Rosa26-EYFP positive (YFP^+^) cell percentage of the tip area divided by that of the capillaries. (B) Linear scatter plot showing correlation of recombination (YFP^+^) ratio between tip cell position and capillary area in *amot*^*ec*+^ retinas (n=24). The data was analyzed using Pearson’s correlation *R*^2^ (*R-squared*) *=0.9479*. (C) The ratio of YFP^+^ (*amot*^*ec*−^) cell percentage at tip cell positions in *amot*^*ec*+^ (n=4) retinas was normalized to 1 and compared that of *amot*^*ec*−^ (n=6) retinas. Retinas with recombination frequencies of 50-85% were chosen for statistical analysis. (D and E) Distribution of YFP^+^ cells in *amot*^*ec*+^ and *amot*^*ec*−^ retinas stained for EC (IB4, in red) and YFP (in green) at > 85%” or 50-85%” recombination frequencies as indicated in the figures. Yellow arrows indicate YFP^+^ tip cells, while white arrows indicate YFP^−^ cells. \*\**P* < 0.01. Scale bars, (D) 100 μm, (E) 50 μm.

### Amot relays mechanical force between matrix and EC

In order to generate functional vessels, ECs are dependent on the active interaction with other cell types, including pericytes, microglia and astrocytes (Fruttiger, 2007). The latter migrate a few days earlier than ECs and act as a guiding template for the patterning of the vascular network in the retina (Fruttiger, Calver et al., 1996, Jiang, Liou et al., 1994). We used the astrocyte marker Glial Fibrillary Acidic Protein (GFAP) to investigate whether *amot* depletion in ECs affected astrocyte patterning in a non-autonomous manner. In *amot*^*ec*+^ retinas, GFAP staining was oriented in a radial pattern around tip cells in the vascular/astrocyte border zone suggesting that tip cells remodeled the astrocyte network by exerting mechanical force (**Figure 4A**). Interestingly, the apparent radial deformations of the astrocytes around tip cells were not observed in *amot*^*ec*−^ retinas (**Figure 4A**). These observations prompted us to investigate whether Amot is required for the interaction between the migrating tip cells and the pre-existing astrocytes.

**Figure 4.**
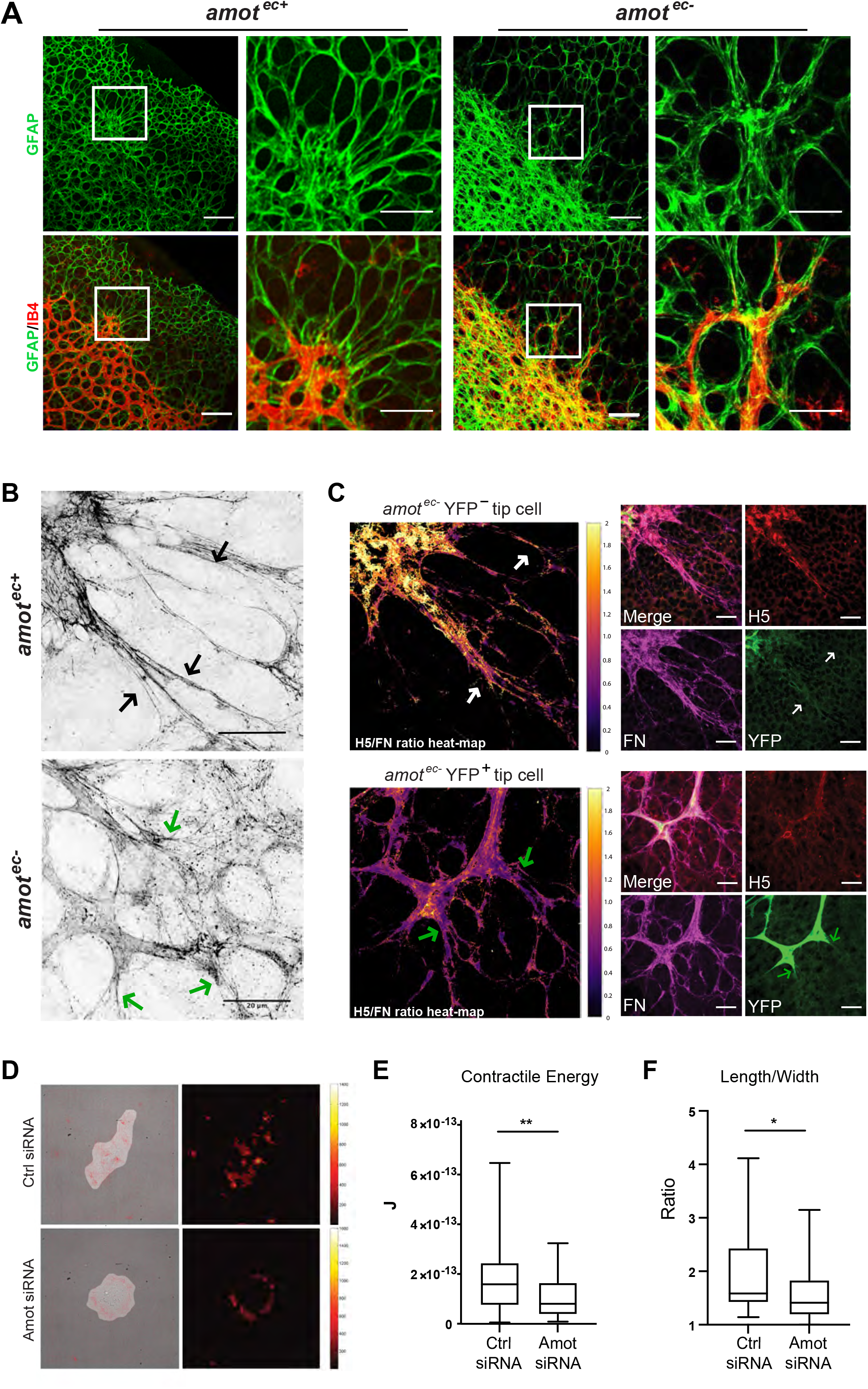
Amot relays force between EC and Fn. IF staining of IB4 (in red) and GFAP (in green) in retinas from *amot*^*ec*+^ and *amot*^*ec*−^ mice. Radial GFAP staining patterns in an area of interaction between astrocytes and EC is indicated by the white box in the *amot*^*ec*+^ retina. Similar structure was not found in *amot*^*ec*−^ retinas. Boxed areas are shown in higher magnification in the right panel. (B) IF staining of Fn was visualized by enhanced-resolution confocal microscopy. Black arrows in *amot*^*ec*+^ tip cell position highlight strained Fn fibrils, while green arrows indicate non-directional and relaxed fibrils in *amot*^*ec*−^ retinas. (C) Immunofluorescent stainings analyzing the ratio of strained vs total Fn in the retina of an *amot* floxed mouse injected with Tamoxifen. The Rosa26EYFP (YFP) marker was used to trace recombined EC (as shown in green). The H5 (H5-myc single-chain antibody) detects strained Fn (as shown in red) whereas Pan-Fn that binds relaxed and strained Fn is shown in magenta. Heatmap images illustrate the ratio of strained vs total Fn. (D) Representative images from traction force microscopy (TFM) experiments. Left panel shows bright field images of single MS1 EC cells plated on a 5 kPa polyacrylamide gel homogenously covered with Fn with the superimposed force vector map. The length and orientation of red arrow indicate the magnitude and directions of exerted traction forces. Stress magnitude was denoted in color scale in Pascal (Pa, right panel). Brighter signals indicate higher force. One pixel is equal to 0.144 μm. (E) Box plot showing contractile energy generated by individual cells from control and Amot siRNA-depleted cells (n≥40 in each group), and the length-to-width ratio quantification of cells plated on the Fn substratum \*\**P* < 0.005; \*\*\**P* < 0.001. Scale bars, (A-C) 50 μm.

As previously mentioned, astrocytes migrate into the retina, prior to the onset of angiogenesis, and deposit a network of ECM which acts as guidance cue to migrating ECs (Turner, Badu-Nkansah et al., 2017). A major component of this ECM is fibronectin (Fn), which forms fibrils that display elasticity when subjected to mechanical force (Klotzsch, Smith et al., 2009). Indeed, *in vitro* evidence indicates that Fn molecules are exposed to cell-derived forces and in response exhibit molecular strain (Lemmon, Chen et al., 2009). To further investigate the interaction of EC tip cells and astrocytes, we first mapped the Fn expression in P6 retinas. IF staining showed the intense expression of Fn in astrocytes ahead of the vascular front and around tip cells as previously reported (Stenzel, Lundkvist et al., 2011), whereas Fn was not detectable in the basal membrane of mature vessels of adult mice (**Figure S4A**). Analysis of Fn expression in post-natal *amot*^*ec*^ retinas did not reveal any significant differences in expression levels (**Figure S4B**).

Next, we employed enhanced-resolution microscopy to visualize Fn fibrils in areas of interaction between endothelial tip cells and astrocytes. Fn fibrils were stretched and aligned with tip cell filopodia in *amot*^*ec*+^, whereas Fn was localized to punctate and shorter fibrils in *amot*^*ec*−^ (**Figure 4B**). Several reports have shown that cell-mediated Fn fibril assembly is regulated by mechano-signals and that cell-generated tractional forces may stretch Fn fibers several fold (Dallas, Sivakumar et al., 2005, Leiss, Beckmann et al., 2008, Zhong et al., 1998). To determine force patterns *in vivo*, we made use of a single-chain antibody (clone H5), generated by Barker and co-workers, that specifically binds the force-extended conformation of the FnIII9-10 integrin-binding domain (Cao, Nicosia et al., 2017). This allows the detection of cryptic epitopes that are masked in a relaxed state but exposed upon mechanical tension. As we have previously published, this method can be used to measure strained Fn in post-natal retinas as well as in pathological conditions such as lung fibrosis (Cao et al., 2017).

A heatmap was generated using the ratio between the signal from the H5, stretch-specific antibody and a pan-Fn antibody detecting both stretched and relaxed forms. Increased ratio of H5 staining was observed in areas where tip cell filopodia made contact with Fn fibrils in *amot*^*ec*+^ retinas, suggesting EC-mediated force generation (**Figure S4C**). We then went on to assess whether inactivation of *amot* would affect the strained conformation of Fn fibrils in the tip cell area. As shown in **Figure 4C**, non-recombined EC (YFP^−^), exhibited a H5/pan-Fn signal ratio similar to that of *amot*^*ec*+^ retinas. By contrast, YFP^+^ EC showed a marked reduction in the ratio of H5/pan-Fn signal.

We went on to use traction force microscopy (TFM) to more precisely measure the force involved in the interaction between EC and the ECM *in vitro*. This method measures cell-generated traction forces by quantifying the deformation of the surrounding ECM (Butler, Tolic-Norrelykke et al., 2002). The murine endothelial cell line MS1 was transfected with siRNA targeting Amot or scrambled controls and subsequent Western Blot (WB) analysis showed that both isoforms of Amot, p130 and p80, were efficiently depleted (**Figure S4D**). Contractile energy was indirectly converted by calculation of force exerted on ECM (Mandal, Wang et al., 2014, Maruthamuthu, Sabass et al., 2011). Consistent with our *in vivo* observations, Amot siRNA depleted cells exhibited a significant decrease in force exerted on the Fn coated polyacrylamide gel (**Figure 4D** and **E**). Additionally, we observed that depletion of Amot resulted in changes in cell shape and did not elongate upon attachment to the ECM (**Figure 4F**).

### Amot is a novel component of the integrin adhesome

We have previously shown that the other members of the Amot protein family link membrane receptors to the cytoskeleton. Thus, we hypothesized that Amot would function in a similar manner by mediating force from the ECM to the cytoskeleton via ECM binding membrane proteins. In support of this notion, we found that GFP-tagged Amot co-localized with vinculin in spreading endothelial cells (**Figure 5A**). In order to identify potential interacting partners, we performed Mass Spectrometry (MS) of proteins that were co-immunoprecipitated with Amot in lysates of primary endothelial BAE (bovine aortic endothelial) cells. All proteins (299) from the list (**Table S1**) were selected for pathway analysis using the Panther 2016 software as described in the *Methods*. The identified pathways are shown in **Figure 5B** with the “Integrin signaling pathway” and “Cytoskeletal regulation by Rho GTPase” appearing as the top regulated pathways (**Table S2**). In support, several integrin subtypes were also present in the MS data, including β3, α5- and β1 (**Figure 5C**).

**Figure 5.**
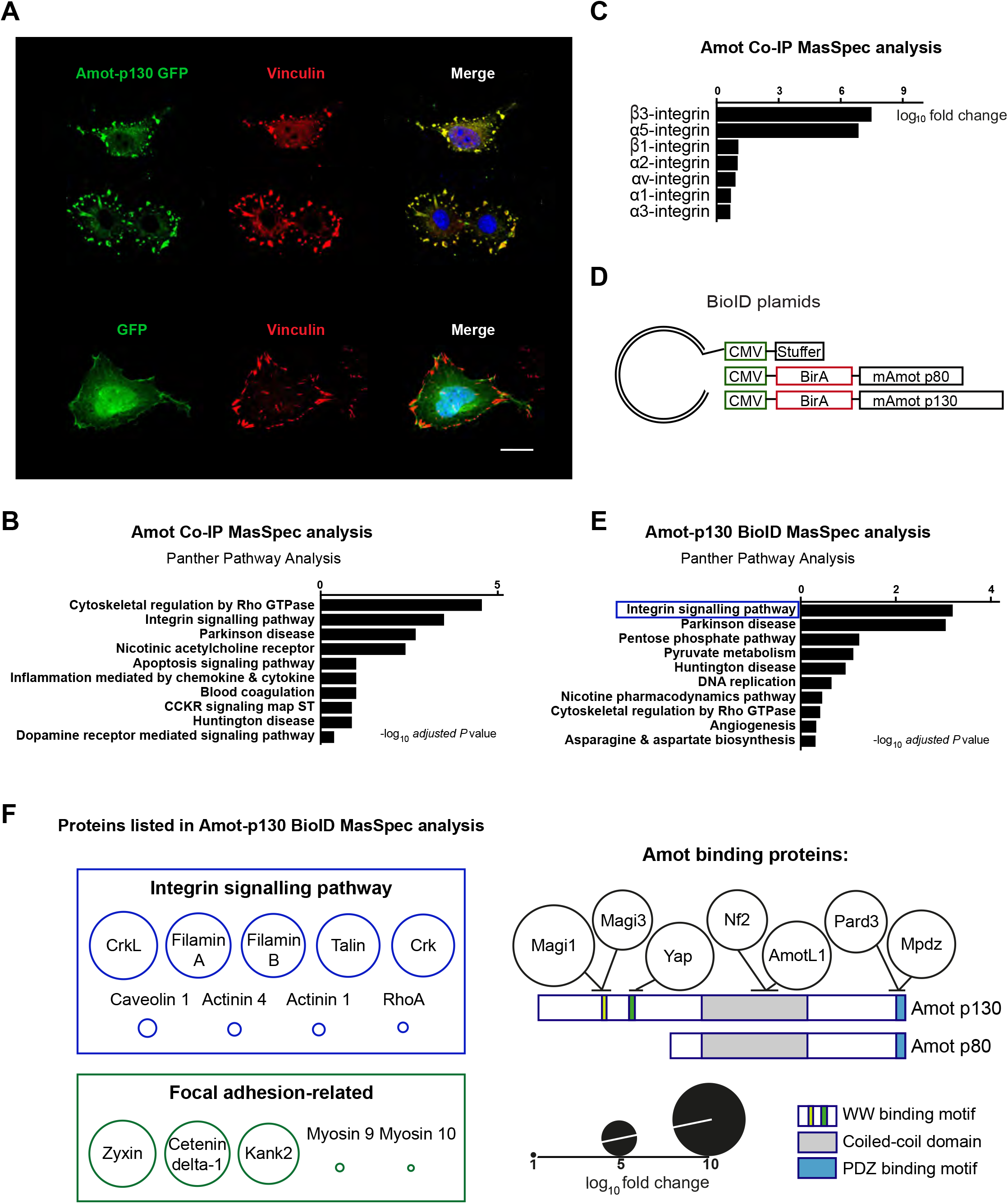
Amot is a novel component of the integrin adhesome. (A) Amot-p130 localizes to focal adhesions. Image shows the localization of GFP-tagged Amot-p130 (in green) which overlaps with vinculin as shown by immunofluorescent staining (in red) in MS1 cell transfected by Amot-p130 GFP plasmids. GFP-alone was used as negative control. (B) Panther pathway analysis of proteins that were co-immunoprecipitated with Amot antibodies. Protein lysates were harvested from the BAE cells culture in 40% confluent for Co-IP using rabbit IgG (n=3) and Amot antibodies (n=2). For each protein, fold change (fc) was calculated, and the ranking is based on positive log_10_ (fc) values, listed in **Table S1**. (C) Integrin subtypes identified as binding partners by MS analysis, which is ranking by log_10_ (adjusted *P* value). See also **Table S1**. (D) Schematic illustrating the BioID plasmids we used for lenti virus particles construction and subsequent transfection and BioID experiments. (E) The bar diagram shows the top 10 pathways enriched among proteins biotinylated by the BirA Amot-p130 construct (n=178) as identified by MS analysis in transfected MS1 cells (fc>0). MS analysis included *empty vector (+/− biotin)*, *Amot p80 BioID +/− biotin*, and *Amot p130 BioID +/− biotin*. Fc was calculated as the subtraction value of average quantity detected in control groups (*empty vector +/− biotin*, *Amot p80/p130 BioID - biotin)*, from the one in *Amot p130 BioID +biotin*. The ranking was based on positive log_10_ (fc) values listed in **Table S3**. (F) The figure shows the identity of proteins identified in the Panther Integrin signaling pathway (blue square). The green square shows identified focal adhesion proteins not part of the Panther integrin pathway grouping. Circle sizes indicate log_10_ (fc) values. The BioID strategy also identified seven known Amot binders which are presented at the right side.

To identify proteins in close proximity of Amot (the Amot adhesome), we used the BioID (proximity-dependent biotin identification) methodology. This method is based on fusing the promiscuous biotin ligase, BirA, with the protein of interest. When expressed in cells it can be induced to biotinylate interacting proteins (Roux, Kim et al., 2018). We fused the BirA to the N-terminus of the p130 and p80 Amot isoforms (schematic shown in **Figure 5D**). The constructs were transfected into MS1 endothelial cells and verified using antibodies against the biotinylase and Amot (**Figure S5A**). BirA tagged to Amot-p130 was localized to cell periphery in sub-confluent cells, which was distinct from those cells expressing the Amot-p80 BirA localizing to intra-cellular vesicles (**Figure S5B**). Both BirA tagged biotinylated proteins was successfully pulled down using streptavidin-beads as analyzed by WB (**Figure S5C**). To identify the interactors of Amot-p130 and -p80, MS was performed on streptavidin affinity purified proteins. 178 proteins with a fold change (fc)>0 were scored as positive hits (**Table S3**). Panther pathway analysis of the positive hits highlighted ‘Integrin signaling pathway’ as the top enriched pathway (**Figure 5E** and **Table S4**). Additionally, from the analysis, we also found zyxin, kank2, catenin delta-1 and myosin 9/10, which have been previously identified as focal adhesion-related proteins (**Figure 5F**) (Bondzie, Chen et al., 2016, Rahikainen, Ohman et al., 2019, Smith, Blankman et al., 2010, Sun, Tseng et al., 2016). The BioID approach also recognized the known direct interactors of Amot such as, Pard3, Nf2 and Yap as shown in **Figure 5F**.

### Amot promotes actin filament formation at focal adhesions

Next, we verified the BioID data by co-immunoprecipitation analysis. As Amot depletion negatively affects endothelial cell migration, we performed the IP analysis at 40% and 100 % confluency. The steady-state levels of the components of the focal adhesion adhesome remained unchanged as shown by WB (**Figure 6A**). Interestingly, the association of Amot to actin and focal adhesion proteins was readily detected in sub-confluent cells but largely lost when cells were confluent (**Figure 6A**). The association of Amot, as well as actin and vinculin, was also verified by kank2 and zyxin pull-down (**Figure S5D** and **S5E**).

**Figure 6.**
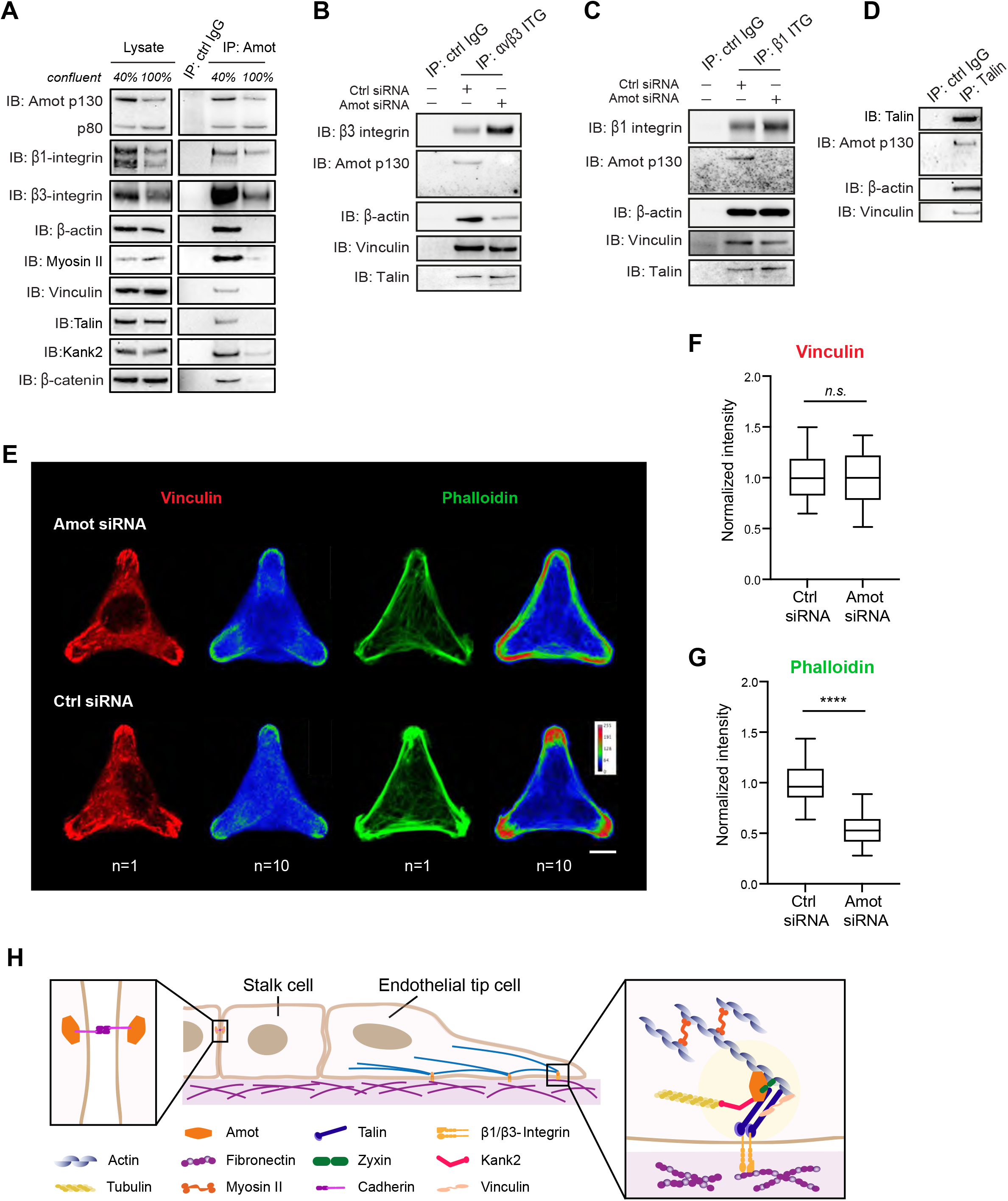
Amot promotes actin filament formation at focal adhesions. (A) Co-immunoprecipitation (Co-IP) analysis of Amot association to the integrin adhesome. Steady state levels of integrin and focal adhesion proteins was analyzed by WB in lysates of BAE cells grown at 40% and 100% confluency. Amot was immunoprecipitated in parallel and subjected to WB analysis with the antibodies indicated in the figure. (B-C) WB analysis of IP using αvβ3-integrin antibody (B) and β1-integrin antibody (C) in BAE cells transfected by control or Amot siRNA. (D) Co-IP and WB analysis of the interaction of Amot with talin. Three independent experiments were performed for all IP experiments. (E) Control or Amot siRNA depleted BAE cells were plated on Y-shaped CYTOO chips for 5 hours. Cells were stained with antibodies against vinculin (in red) and actin was visualized by phalloidin staining (green). Images of overlay of 10 cells are also shown. Scale bars: 10 μm. (F and G) The intensities of vinculin and phalloidin signals at three vertexes of Y-shape were measured and normalized to the signaling at the central area within the same cell. *n.s.,* not significant; \*\*\*\**P* <0.0001. (H) Hypothetical schematic of focal adhesions where Amot links the actin filaments to Fn in the ECM through integrin associated adhesive complex in migratory endothelial tip cells.

To verify the influence on integrins, we harvested protein lysates from control siRNA and Amot siRNA treated BAE cells. Interestingly, both αvβ3 and β1 integrins bound specifically to Amot p130 thus arguing that the interaction is mediated via the N-terminal domain (**Figure 6B** and **C)**. Immunoprecipitation of talin revealed a similar Amot isoform preference (**Figure 6D).**

Our data indicated that Amot did not bind actin under confluent conditions. Furthermore, depletion of Amot did not affect the association of talin or vinculin to αvβ3 and β1 integrins (**Figure 6B** and **C)**. However, the immunoprecipitation of αvβ3 in Amot depleted cells resulted in less binding of actin even though steady stat levels remained unchanged (75% reduction (**Figure 6B**, and **Figure S5F-G)**.

We hypothesized that Amot may affect the association or polymerization of actin filaments in response to integrin activation. To compare the focal adhesion organization in BAE cells transfected with control or Amot siRNA, we plated individual cells on Y-shaped Fn-coated CYTOO chips. Amot depletion did not affect the recruitment of vinculin to sites of focal adhesion whereas the intensity of actin staining was significantly decreased (**Figure 6E**, quantification in **Figure 6F** and **G**). Taken together, we present evidence that Amot is a novel component of the integrin adhesome in EC where under pro-migratory or sub confluent conditions promote actin filament formation (hypothetic schematic in **Figure 6H**).

### Amot is required for tumor angiogenesis

The ECM of tumors is heterogenous and the composition varies dependent on tumor origin and subtype. This may explain why endothelial-specific inactivation of Fn, α5 and αv integrin severely affects developmental angiogenesis and causes lethality *in utero* but has little effect on tumor angiogenesis in the adult stage (Murphy et al., 2015). As we observed expression of Amot in blood vessels of human tumors, we investigated whether Amot is also implicated in tumor angiogenesis. For this purpose, we studied pathological angiogenesis in the tumor model with Lewis Lung Carcinoma (LLC) transplantation, and in the transgenic mouse model of breast cancer MMTV-PyMT. The LLC model has been used extensively to assess efficacy of anti-angiogenic therapies. Mice were injected with tamoxifen two weeks before and after subcutaneous injection with LLC cells (diagram in **Figure 7A**), in order to provide either *amot*^*ec*+^ or *amot*^*ec*−^ environment for tumor growth. As shown in **Figure 7B**, LLC tumors from *amot*^*ec*−^ animals were significantly smaller as assessed by tumor weight (**Figure 7C**).

**Figure 7.**
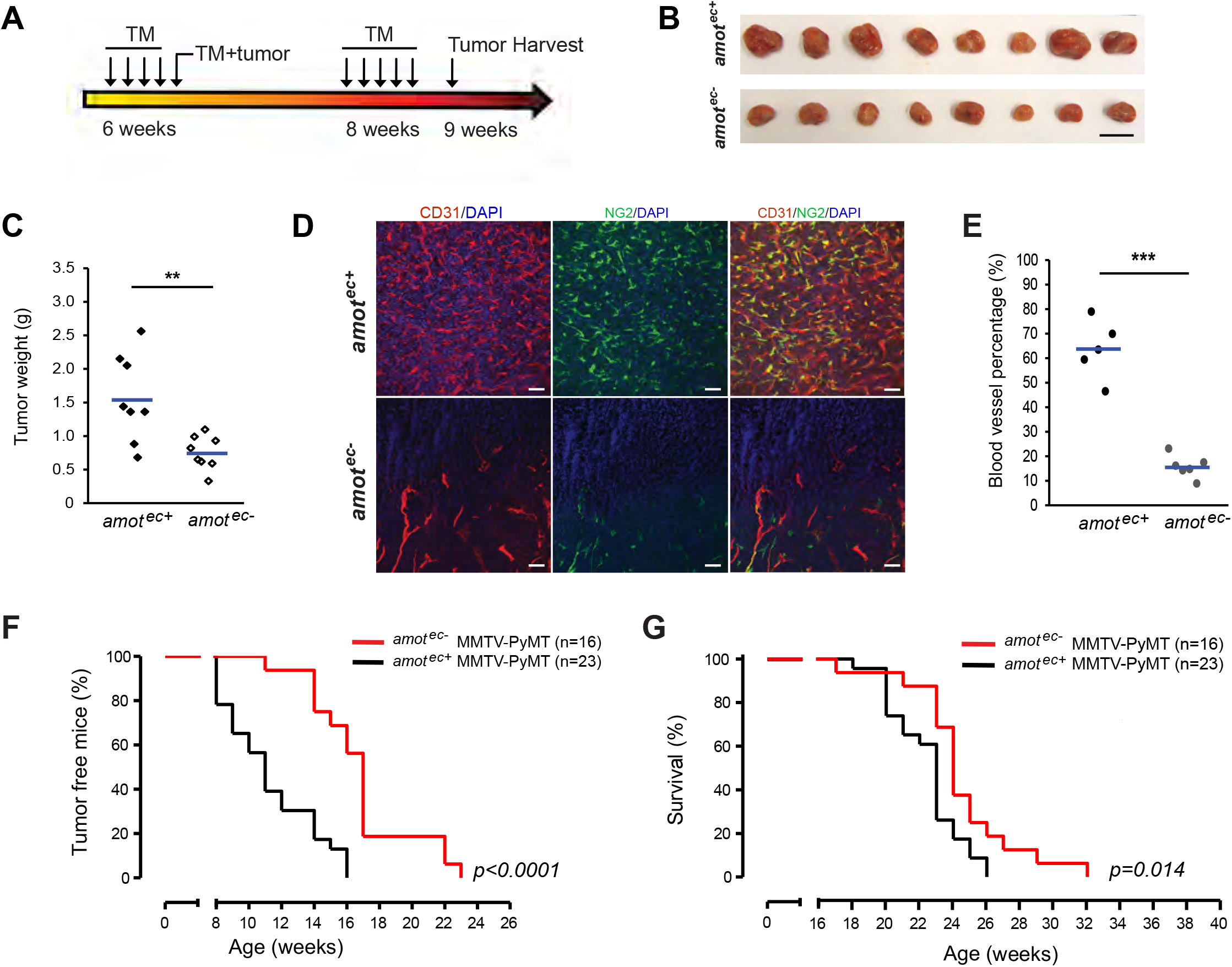
Abrogation of endothelial Amot inhibits tumor growth in transplantation and transgenic tumor mouse models. (A) Schematic of experimental procedure. Tamoxifen injections were performed in *amot*^*ec*−^ mice at six weeks. Lewis lung carcinoma (LLC) tumor cells were injected with tamoxifen together at eight weeks. LLC tumors were harvested when the largest tumor reached a maximal diameter of 10 mm. (B) Representative allografted tumors dissected from *amot*^*ec*+^ (n=8) and *amot*^*ec*−^ (n=8) mice. Scale bar, 10mm. (C) Dot plot analysis of tumor weight in *amot*^*ec*+^ (n=8) and *amot*^*ec*−^ (n=8) tumors. (D) IF staining of LLC tumor vibratome sections (100 μm thick) by anti-CD31 (in red) and anti-NG2 (in green) antibodies. Note the regressing blood vessels in *amot*^*ec*−^ tumor. Nuclei were visualized by DAPI (in blue). Scale bar, 50 μm. (E) Analysis of blood vessel coverage by quantification of CD31 staining on cryosection of *amot*^*ec*+^ (n=5) and *amot*^*ec*−^ (n=6) tumors. (F) Analysis of breast tumor incidence in MMTV-PyMT mice with *amot*^*ec*−^ (n=16) and *amot*^*ec*+^ (n=23). (G) Kaplan-Meier survival curves of *amot*^*ec*+^ MMTV-PyMT (black line, n=23) and *amot*^*ec*−^ mice (red line, n=16). Interval omitted is from 2nd-16th week. \*\**P* < 0.005; \*\*\**P* < 0.001.

Immunostaining of sections from resected tumors showed a marked decrease in vascular density and an increased percentage of necrosis in *amot*^*ec*−^ tumors, compared to *amot*^*ec*+^ (**Figure 7D** and **E**).

Next, we used the MMTV-PyMT mice, in which Polyoma Middle-T oncogene driven by the MMTV promoter rendered multiple-site breast tumor occurrence at around 8-9 weeks of age (Lin, Jones et al., 2003). We crossed *amot*^*ec*−^ mice into the MMTV-PyMT background and induced *amot* endothelial specific depletion at week four by tamoxifen injection. Analysis of early onset tumor occurrence showed an average of six weeks longer latency in *amot*^*ec*−^ mice, compared to *amot*^*ec*+^ mice, confirming the protective effect of *amot* deletion on tumor incidence by inhibiting angiogenesis (**Figure 7F**). Furthermore, overall survival was significantly improved in *amot*-ablated mice (**Figure 7G**).

## Discussion

In this report, we show that Amot is a novel component of the integrin adhesome and is essential in relaying force between the ECM and the cellular cytoskeleton. Furthermore, disruption of this relationship by EC-specific gene deletion of Amot resulted in inhibition of both normal and pathological angiogenesis. This highlights the importance of Amot in maintaining normal and pathological transmission of forces in vascular expansion.

Early work by Ingber and Folkman showed that mechanical interactions between ECs and ECM regulated capillary development *in vitro* (Ingber & Folkman, 1989). This led to findings that ECM properties, such as matrix stiffness, control cell shape and function by modulating the cytoskeletal network (Ingber, Prusty et al., 1995). The forces transmitted through the integrins are in the picomolar range and involves integrin clustering and the formation of a mechanosensory adhesome. This adhesome involves the integrin binding protein talin, which relays force by unfolding subdomains and exposing cryptic protein interaction sites. The integrin adhesome is highly complex involving over 200 proteins that have been implicated in the cell-ECM interaction (Horton & Humphries, 2016). Our data provide evidence that Amot is a novel component of this integrin adhesome. Despite the intense research focusing on the mechanotransduction of cell-matrix adhesions, the role of Amot in force transmission has not been described. This may be explained that the fibroblasts typically involved in cellular adhesion studies lack Amot expression. Indeed, our studies show that in mice and humans, Amot is primarily expressed in ECs during developmental angiogenesis and in tumor angiogenesis. Recent reports have also shown that Amot is expressed in cells of the inner cell mass of the blastocyst, there controlling cellular differentiation as well as in the developing neural system. The developing of the neural system is quite reminiscent of the fractal vascular patterning of the retina. Amot inactivation in neural cells of the developing mouse brain also affects cellular branching in a similar manner. In the neural system, Amot triggers the actin filament formation and thereby stabilizes dendritic spines (Wigerius & Quinn, 2018). In our previously published data, we show that filopodia are destabilized upon Amot depletion. Indeed, inactivation of Amot in zebrafish resulted in a “treadmilling phenotype” where the tip cells are extending filopodia that are unstable and extend and retract at a high rate (Aase et al., 2007). This further argues that Amot is required for the stability of filopodial extensions by establishing contacts between the ECM and the cytoskeleton.

The data presented in this report show that Amot is essential for the transmission of force between the Fn component of ECM and the cytoskeleton of migrating ECs. Several independent experiments support this notion. EC-specific knockout of Amot led to changes in Fn fibrillogenesis as analyzed by enhanced-resolution microscopy *in vivo*. We exploited the fact that Fn changes conformation when exposed to mechanical strain, thus unmasking a cryptic epitope detected by the H5 single chain antibody. Amot deficient cells had a lower ratio of strained vs total Fn in the adjacent matrix. This was supported by direct measurements of force using TFM on Fn coated matrix *in vitro*. On the intra-cellular side, Amot regulates the formation of contractile actin filaments. We have previously shown that Amot not only binds actin but also triggers the formation of actin filaments. Indeed, Amot has been reported to control Rho-GAP activity, which may explain the extensive formation of actin fibers upon p130Amot overexpression (Wells, Fawcett et al., 2006, Yi, Shen et al., 2013). How Amot or AmotL2 triggers Actin filament formation is yet unclear, but both proteins bind the Rho-GEF protein PLEKHG1/Syx, which promotes EC migration by controlling RhoA GTPase activity (Wells et al., 2006, Yi et al., 2013). Furthermore, in migrating ECs Amot is required for the spatially restricting Rho-GTPAse to cellular protrusions or lamellipodia as shown by Rho-FRET analysis (Ernkvist, Luna Persson et al., 2009).

Taken together, our data point to a role of Amot in integrating mechanical cues sensed by the migrating endothelium that is translated to actomyosin contractility that is required for EC migration.

## Methods

### Mice

The *amot*^*flox*^ mice, carrying a loxP-flanked *amot* gene, were crossed with Cdh5(PAC)^CreERT2^ and ROSA26-EYFP double transgenic mice. *Amot*^*ec*+^ and *amot*^*ec*−^ are the abbreviations used for depicting amot^wt^-Cdh5(PAC)^CreERT2^-ROSA26-EYFP and amot^flox^-Cdh5(PAC)^CreERT2^-ROSA26-EYFP mice, respectively, as the *amot* gene localizes to the X chromosome. Mice were also crossed with mice carrying Polyomavirus Middle T under control of the MMTV promoter (MMTV-PyMT) in the animal facility of the Molecular Biotechnology Center (Turin) and treated in conformity with European Guidelines and policies, as approved by the ethical Committee of the University of Torino. All the mice in this report were in C57BL/6 background. Biopsies from mouse tail-tip and ear were used for genotyping of each mouse included in these studies. Both female and male animals were included. Ethical permission was granted by the North Stockholm Animal Ethical Committee. All animal housing and experiments were carried out in accordance with the guidelines of the Swedish Board of Agriculture.

### Tamoxifen administration

For retinal angiogenesis study, tamoxifen (400 μg/mouse/day) was administered by intraperitoneal (IP) injection from postnatal day 1 (P1) to P3. For mice over 6 weeks old, tamoxifen (2 mg/mouse/day) was given for 5 continuous days.

### Retinal angiogenesis assay

Mouse retinal angiogenesis assay was performed as previously published (Pitulescu et al., 2010, Stahl, Connor et al., 2010). In brief, eyeballs were dissected at different time points including: P1-7, P10 and adult. A portion of them was fixed in 4% paraformaldehyde (PFA) for 1.5 hours (1.5 h) before whole-mount immunostaining. For retinal protein analysis, retinas were dissected out from eyeballs freshly and homogenized on ice to extract protein.

### Tumor allograft model

The *amot*^*ec*+^and *amot*^*ec*−^ male mice (≥8 mice per group) at 6 weeks of age were injected by tamoxifen to induce *amot* depletion. Lewis lung carcinoma cells (LLC, 0.5 × 10^6^/mouse) were injected subcutaneously on the last day of tamoxifen injection. Tumor growth was manually inspected twice per week and tumor volumes were calculated according to the formula: 0.52 × length × width × width (cm^3^). Starting from 2 weeks after the first injection, the same dosage of tamoxifen was injected into the tumor-bearing mice for 5 days. When tumors reached the diameter of 10 mm, mice were sacrificed according to the ethical permission.

### Cell culture

MS1 cells (Mile Sven 1, ATCC CRL.2279), SV40-transformed mouse endothelial cells, were cultured in RPMI-1640 medium supplemented with 10% FBS and 1% penicillin/streptomycin (P/S) at 37 °C, 10% CO_2_ condition. Primary BAE (Bovine Aortic Endothelial) cells purchased from Sigma were cultured in Bovine Endothelial Cell Growth Medium at 37 °C, 10% CO_2_ condition. LLC cells were cultured in DMEM medium supplemented with 10% FBS and 1% P/S. All products mentioned above are listed in **Table S5**.

### Transient transfection for knockdown and overexpression

For siRNA transfections, MS1 and BAE cells were seeded the day before transfection in 10 mm culture Petri dishes pre-coated with 0.5% Fn, at 30% confluent. Both control and Amot siRNAs with final concentration of 25 nM were transfected into the cells with Lipofectamine^®^ RNAiMAX Reagent according to the manufacturer’s protocol. The cells were allowed to grow for 56 h, and sequentially re-plated to 15 mm dishes or 8-well chamber slides for another 16 h culture, which is considered sub-confluent status (40%). Immunofluorescence staining or WB were processed afterwards.

For Amot overexpression studies, MS1 cells were transfected with mouse Amot-p130-GFP or GFP plasmid using Lipofectamine 3000 according to the protocol of the manufacturer. Immunostaining was performed 24 h post-transfection.

### Proximity biotinylation (BioID)

BioID expression constructs were created by cloning cDNA fragments encoding mouse Amot p130 or mouse Amot p80 into the destination vector pcDNA3.1 -BirA(R118G) by VectorBuilder to obtain mouse p130 Amot and mouse p80 Amot fused to the N-terminus of BirA*. Empty vector pcDNA3.1 was used as negative control. All constructs were verified by Restriction Enzyme Digestion. BioID constructs were packaged into lenti virus using Lipofectamine 3000 Transfection Reagent according to the protocol of the manufacturer. MS1 cells were used as target cells for Lenti virus transfection with 0.5 mg/mL geneticin. Stable transfected cells were cultured in RPMI-1640 medium supplemented with 10% FBS and 0.5 mg/mL geneticin. For BioID analysis, MS1 cells were treated 16 h with 50 μM biotin, followed harvesting in lysis buffer consisting of 50 mM Tris·Cl pH 7.4, 8 M urea, 1 mM DTT supplemented with protease inhibitors. Lysates was supplemented with 1% Triton X-100 before being sonicated. Biotinylated proteins were purified by incubating the cell lysates with Streptavidin beads overnight at 4 °C. After five washes with 8 M urea in 50 mM Tris·Cl pH 7.4 and one wash with 50 mM Tris·Cl pH 7.4, beads were resuspended in PBS and ready for WB and MS analysis. Detailed sequencing information is provided in **Table S5-6**.

### Immunofluorescence (IF) staining

Cells were fixed by 4% PFA for 10 min at room temperature (RT) and then were permeabilized with 0.1% Triton X-100 for 1 min. After 1 h blocking with 5% horse serum in PBS, cells were incubated with primary antibodies for 2 h followed by Alexa Fluor^®^ conjugated secondary antibodies for 1 h at RT.

Mouse retinal whole-mount staining protocol has been previously described (Pitulescu et al., 2010). In brief, fixed retinas were permeabilized and blocked in PBS containing 0.3% Triton X-100 and 2% BSA at 4 °C overnight. Primary antibodies in Pblec buffer (1.0% Triton X-100 plus 0.1 M MgCl_2_, 0.1 M CaCl_2_, 0.01 M MnCl_2_ in PBS) were added to the retina and incubated at 4 °C overnight and subsequently incubated with the fluorescent-conjugated secondary antibodies.

Digital images were acquired using a Zeiss LSM 700 confocal microscope/Airyscan-resolution microscopy and analyzed by Image J (NIH). All the primary and secondary antibodies used were listed in **Table S6**.

### H5-myc single-chain variable fragment (scFv) antibody

Conformation-sensitive single-chain antibody clone H5 with myc tag (H5-myc) was developed using phage display against engineered Fn fragments, and has previously been shown to detect a force-induced extension in the FnIII9-10 integrin-binding domain(Cao et al., 2017). H5-myc was used as ordinary primary antibody for retinal staining as described above, except with a 2 h RT incubation with Anti-myc antibody for amplifying the signal before secondary antibody addition. Samples were additionally stained with a pan-Fn antibody to control for total Fn in the matrix. To visualize the degree of force-induced Fn unfolding in the matrix, ratiometric images of H5-myc: Pan-Fn was generated using a custom Matlab program. Images were first gated using Otsu's method, and masked to analyze only regions where both H5-myc and pan-Fn passed the threshold. Fluorescence intensity of the H5-myc signal was then divided by fluorescence intensity of the pan-Fn signal on a pixel-by-pixel basis, and plotted using a heatmap to display the ratiometric values for each pixel, where a higher ratio indicates greater force-induced unfolding of Fn.

### Traction force microscopy (TFM)

5 kPa polyacrylamide gels embedded with fluorescent beads (Thermo Fisher Scientific, F8807) were prepared as previously described (Tseng, Wang et al., 2011). For quantitative force measurements, MS1 cells, 72 h after siRNA transfection, were trypsinized and seeded as single cells on top of the hydrogel. Images were acquired using an Eclipse Ti inverted microscope (Nikon) with a 60 x oil immersion objective with 1.5 x zoom (Nikon 1.4 NA) and a Zyla sCMOS camera (Andor), both driven by Andor iQ3, as well as a temperature control system set at 37 °C. Images of fluorescent beads within the relaxed and stressed hydrogel were taken before and after cell detachment, respectively, and one bright field image of the cell doublet was recorded.

The method used for force calculations on continuous substrates was described previously (Butler et al., 2002, Mandal et al., 2014). All image processing and calculations have been performed in matlab. The displacement field was calculated using a combination of particle image velocimetry and single particle tracking. After drift correction, the mean displacement was calculated by cross-correlating corresponding sub-windows (size 9.22 μm) of the stressed and relaxed bead image. Following, single particle tracking within each sub-window was performed, which led to a high spatial resolution and accuracy of the displacement measurements. The final displacement field was obtained on a regular grid of 1.15 μm by linear interpolation. Forces were reconstructed using Fourier transform traction cytometry with zeroth-order regularization. From this, the contractile energy U, which represents the total energy transferred from the cell to the elastic distortion of the substrate, was calculated as follows:

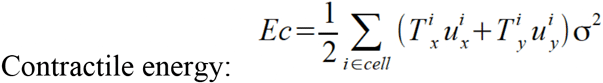

### Western blot (WB) analysis and Co-immunoprecipitation (Co-IP)

Cells were scraped directly from cell culture dishes in lysis buffer (50 mM Hepes buffer, 150 mM NaCl, 1.5 mM MgCl_2_, 1 mM EGTA, 10% Glycerol, 1% Triton X-100), with 1 x protease inhibitors at 4°C and thereafter. Mouse retina were dissected, cut into small pieces and then immersed in lysis buffer. Samples were homogenized using a tissue homogenizer (VWR) in order to improve protein extraction. SDS sample buffer containing 10% sample reducing agent was added to cell lysates and proteins were separated in a polyacrylamide Bis-Tris 4-12% gradient gel, and transferred to a nitrocellulose membrane. The membrane was blocked in PBS containing 5% non-fat milk and 0.1% Tween 20 and sequentially incubated with the primary antibody and horseradish peroxidase (HRP) conjugated secondary antibody. Labelled proteins were detected using Western Lightning Plus-ECL.

Cell lysates were pre-treated with protein G sepharose beads for 1.5 h. 2 μg Amot/β1-/αvβ3-Integrin antibody or rabbit/rat/mouse control IgG was incubated overnight with agitation. Bound proteins were harvested using protein G beads for 2 h, followed by five consecutive washes with lysis buffer. Protein complexes were further analyzed by WB. Primary/secondary antibodies and reagents used are listed in **Table S6**.

### Mass spectrometry (MS) sample preparation

IP samples were dissolved in 200 μL lysis buffer (4% SDS, 50 mM HEPES pH 7.6, 1 mM DTT). After heating at 95 °C for 5 min and centrifugation at 14,000 g for 15 min, the supernatant was transferred to new tubes. Protein digestion was performed using a modified SP3-protocol (Hughes, Foehr et al., 2014). In brief, each sample was alkylated with 40 mM chloroacetamide and Sera-Mag SP3 bead mix (20 μL) was transferred into the protein sample together with 100% acetonitrile to a final concentration of 70%. The mix was incubated under rotation at room temperature for 20 min. The mixture was placed on the magnetic rack and the supernatant was discarded, followed by two washes with 70% ethanol and one with 100% acetonitrile. The beads-protein was reconstituted in 100 μL trypsin buffer (50 mM HEPES pH 7.6 and 0.8 μg trypsin) and incubated overnight at 37 °C. The eluted samples were dried in a SpeedVac. Approximately 10 μg was suspended in LC mobile phase A and 1 μg was injected on the LC-MS/MS system.

### LC-MS/MS analysis and peptide/protein identification

Online LC-MS was performed using a Dionex UltiMate™ 3000 RSLCnano System coupled to a Q-Exactive-HF mass spectrometer (Thermo Scientific). IP samples were trapped on a C18 guard-desalting column (Acclaim PepMap 100, 75 μm x 2 cm, nanoViper, C18, 5 μm, 100 Å), and separated on a 50 cm long C18 column (Easy spray PepMap RSLC, C18, 2 μm, 100 Å, 75 μm x 50 cm). The nano capillary solvent A was 95% water, 5% DMSO, 0.1% formic acid; and solvent B was 5% water, 5% DMSO, 95% acetonitrile, 0.1% formic acid. At a constant flow of 0.25 μL min^−1^, the curved gradient went from 2% B up to 40% B in 240 min, followed by a steep increase to 100% B in 5 min.

FTMS master scans with 70,000 resolution (and mass range 300-1700 m/z) were followed by data-dependent MS/MS (35,000 resolution) on the top 5 ions using higher energy collision dissociation (HCD) at 30-40% normalized collision energy. Precursors were isolated with a 2 m/z window. Automatic gain control (AGC) targets were 1e^6^ for MS1 and 1e^5^ for MS2. Maximum injection times were 100 ms for MS1 and 150-200 ms for MS2. The entire duty cycle lasted ~2.5 s. Dynamic exclusion was used with 60 s duration. Precursors with unassigned charge state or charge state 1 was excluded and an underfill ratio of 1% was used. The proteins detected in those samples were input to online Enrichr (Ma’ayan Laboratory, Computational systems biology) for Panther pathway analyses (Chen, Tan et al., 2013, Kuleshov, Jones et al., 2016). Calculation of “Combination score” is explained in Enrichr website https://amp.pharm.mssm.edu/Enrichr/help#background&q=4.

### Statistical analysis

Graphs and statistical analysis were performed using GraphPad Prism software. Statistical significance in Figure 2C, 2D, 2E, 2G, 2I, 3C, 4E, 4F, 6G, 6H, 7C, 7E, Figure S3B-D, S5B, S5E and S5F were determined by unpaired two-tailed Student’s t-test. Data in Figure 3B was analyzed by linear regression. Tumor incidence/survival curve (Figure 7F and 7G) were compared by Log-rank (Mantel-Cox) test. The numbers of biological replicate samples and *P* values (n.s., not significant, \**P* < 0.05, \*\**P* < 0.01, *** *P* < 0.001) are provided in individual figures. *P* < 0.05 were considered statistically significant.

## Acknowledgments

We thank Dr. Akihiko Shimono, National University of Singapore for sharing Amot^floxed^ conditional knockout mouse and Dr. Ralf H. Adams from University of Münster for the Cdh5(PAC)^CreERT2^ and ROSA26-EYFP transgenic mice. Mass spectrometry analysis was performed by the Clinical Proteomics Mass Spectrometry facility, Karolinska Institutet/Karolinska University Hospital/Science for Life Laboratory.

This study was supported by following grants: Swedish Cancer Society, the Swedish Childhood Cancer Foundation, Cancer Society of Stockholm and the Swedish Research Council, The Swedish Heart and Lung Society, Cancer and Allergy Foundation, Knut and Alicia Wallenberg Foundation (Y.Zhang, S.K., Y.Zheng, Yumeng Zhang, D.C. and L.H.); PhD scholarship (No.2011622044/201808130149) from Chinese Scholarship Council (Y.Zhang/Yumeng Zhang); grants from Uehera Memorial Foundation in Japan (S.K.); grants of Fondazione Ricerca Molinette Onlus 8893/5 and Italian Association for Cancer Research - IG 11675 (G.B. and F.C.); Iris, Stig och Gerry Castenbäcks Stiftelse for Cancer Research (N.T.); grant ANR-17-CE30-0032-01(K.H. and M.B.).

## Author contribution

Y.Zhang wrote the manuscript, maintained Amot conditional KO mouse model, performed experiments and data analysis, drew schematics and finalized all figures. Yumeng Z., S.K. helped with mouse handling, performed experiment and data analysis. Y.Zheng established Amot conditional KO mouse model, performed experiments and data analysis. G.B. and F.C. backcrossed PyMT tumor-prone mice with *amot*^*ec*+^ mice to generate *amot^ec+^/*PyMT and *amot^ec-^/*PyMT mice. G.B., J.N., K.H., D.C., M.B. and T.B. performed experiment and data analysis. L.H. contributed to design experiments and manuscript writing. All authors reviewed the manuscript prior to submission.

