## Supplemental Figures and Tables for "The Amot/Integrin protein complex transmits mechanical forces required for vascular expansion"

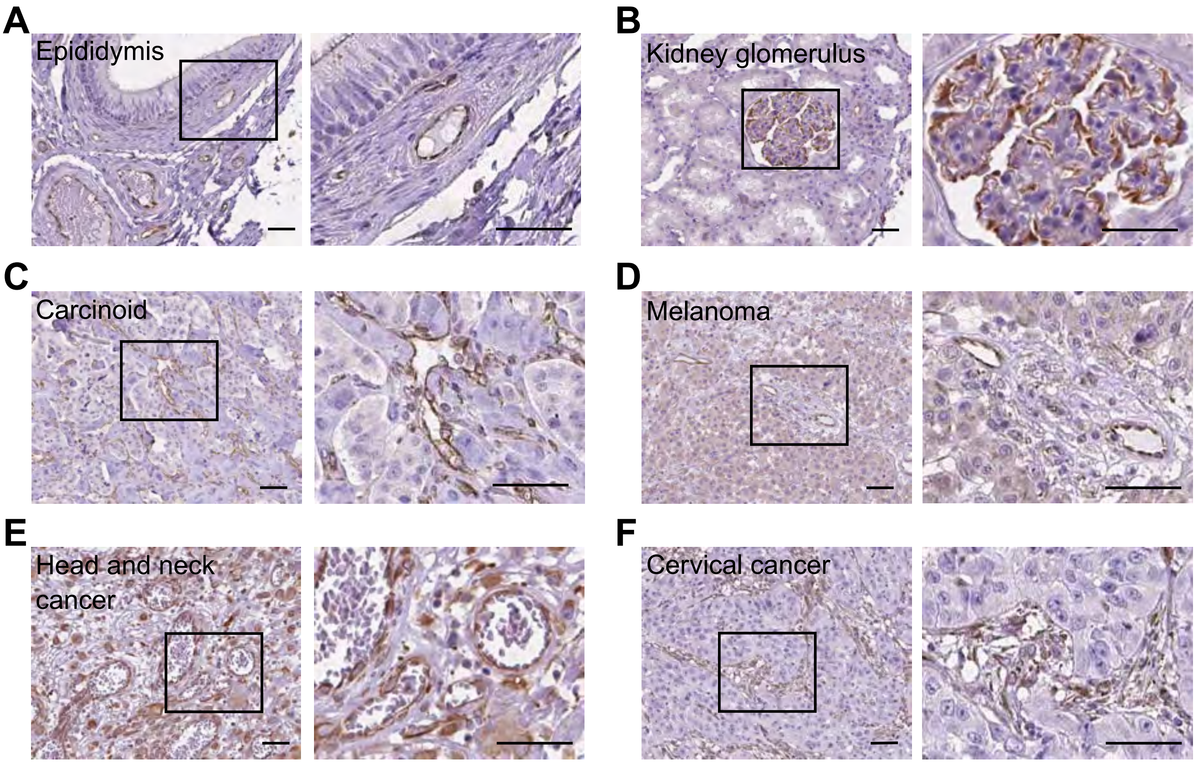

A
Epididymis
B
Kidney glomerulus
C
Carcinoid
D
Melanoma
E
Head and neck cancer
F
Cervical cancer

**Figure S1. Amot expression in vasculature of human normal and tumor tissues.**

(A-F) Representative IHC images of Amot expression (in brown) is shown in section of epididymis (A), kidney glomerulus (B) and pathological tissue sections, including carcinoid (C), melanoma (D), head and neck cancer (E) and cervical cancer (F). Magnification of the boxed area at left panels is shown on the right side. Scale bars, 50 μm.

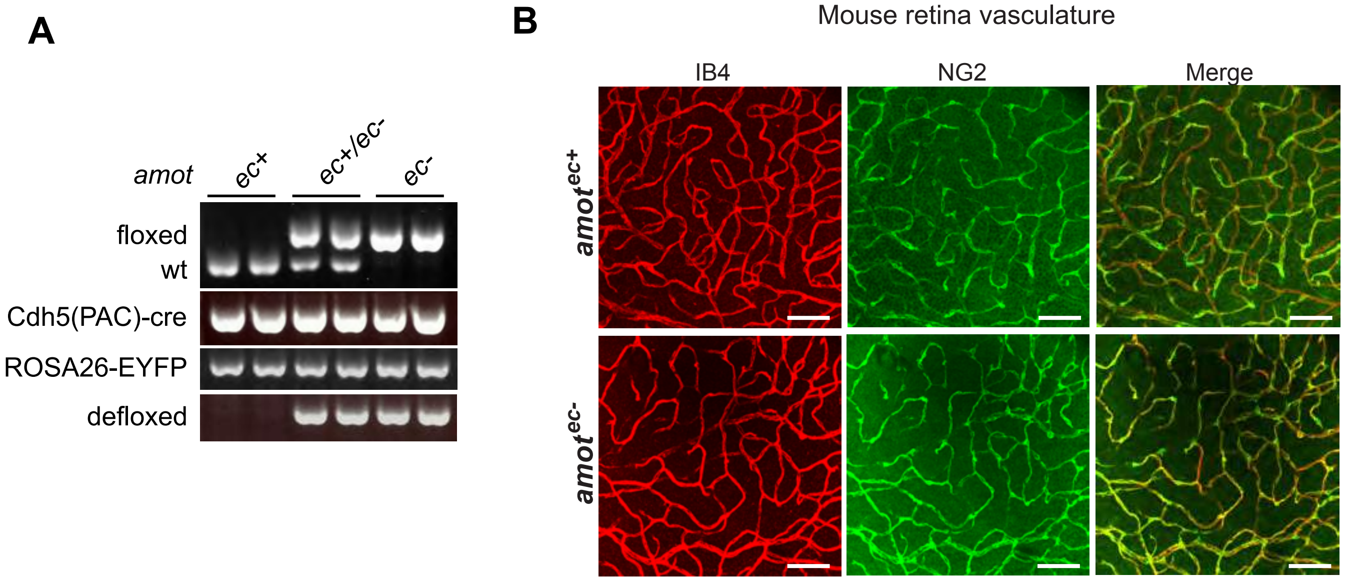
A
amot
ec+
ec+/ec-
ec-
floxed
wt
Cdh5(PAC)-cre
ROSA26-EYFP
defloxed
B
Mouse retina vasculature
IB4
NG2
Merge
amot^ec+
amot^ec-

**Figure S2. Vasculature in adult retina was not affected in *amot*$^{ec-}$ mice.**

(A) Genotyping of *amot*$^{ec+}$, *amot*$^{ec+/ec-}$ and *amot*$^{ec-}$ mice. DNA was isolated from ear/tail tissue. Two genomic DNA samples from each group were randomly chosen for PCR analysis for *amot*-floxed, Cdh5(PAC)-CreERT2, Rosa26-EYFP reporter and *amot*-defloxed. As the *amot* gene is localized on the X chromosome the male genotype can be *amot*$^{ec+}$ or *amot*$^{ec-}$, while the females can be *amot*$^{ec+/ec+}$, *amot*$^{ec+/ec-}$ or *amot*$^{ec-/ec-}$. However, in this study, *amot*$^{ec+/ec-}$ was not included, and therefore we used *amot*$^{ec+}$ and *amot*$^{ec-}$ to represent wild-type and *amot* knockout mice, respectively. (B) Retinal fluorescent images with blood vessel network (IB4, in red) and mural cell staining (NG2, in green) in *amot*$^{ec+}$ and *amot*$^{ec-}$ retinas of adult age (24 weeks, n=6 for each group. Scale bars, 100 µm.

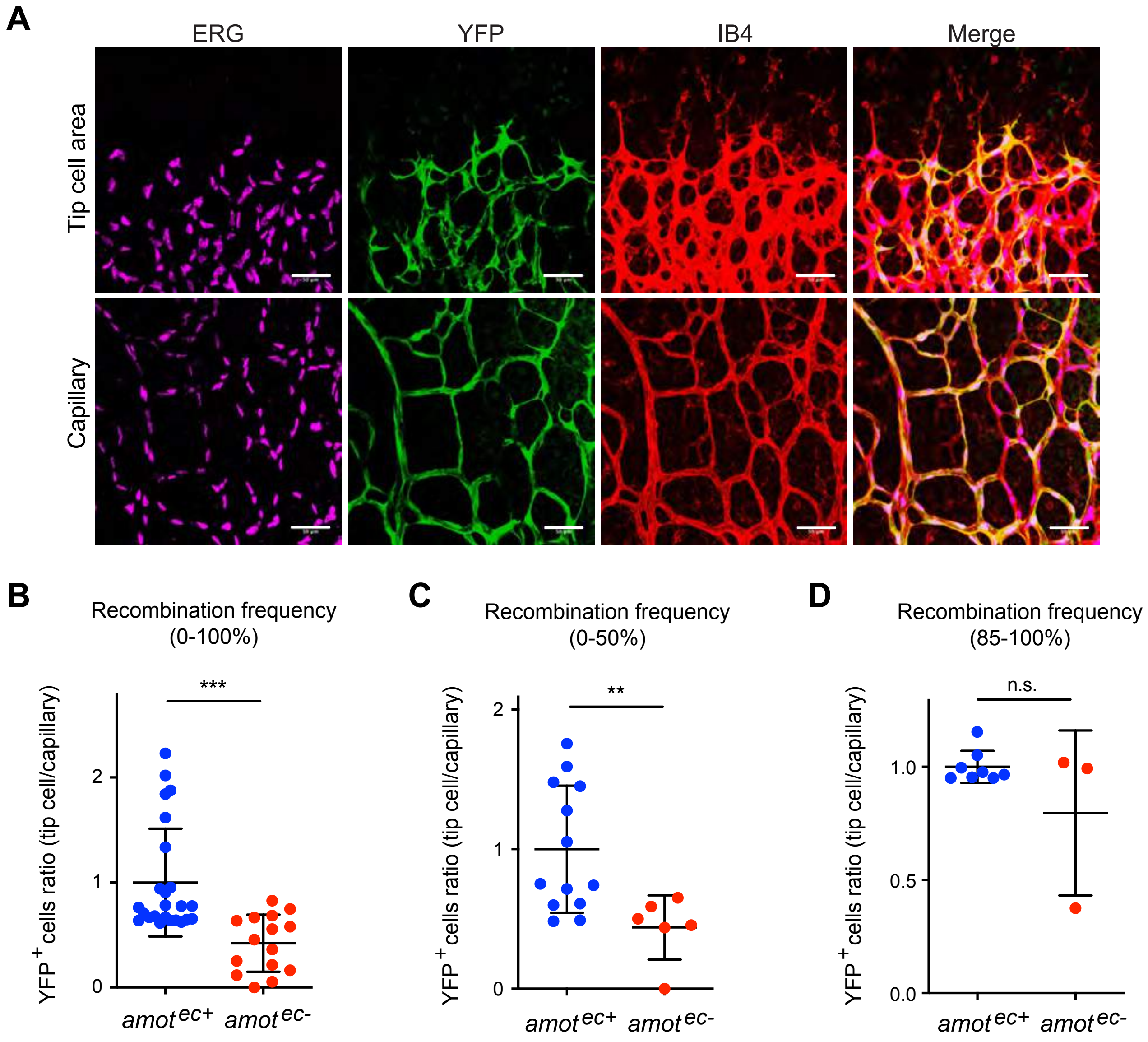
A
ERG
YFP
IB4
Merge
Tip cell area
Capillary
B
Recombination frequency (0-100%)
***
YFP+ cells ratio (tip cell/capillary)
amot ec+
amot ec-
C
Recombination frequency (0-50%)
**
D
Recombination frequency (85-100%)
n.s.

**Figure S3. Amot enables ECs to take tip cell position.**

(A) Vasculature in tip cell position and capillary area was visualized by IB4 staining (in red) in *amot*$^{ec+}$ retinas of P6. ERG (in magenta) as specific nuclear EC marker was stained to facilitate the quantification of Rosa26-EYFP positive (YFP$^{+}$) EC cells (in green). Scale bars, 50 μm. (B-D) Quantification of YFP$^{+}$ EC ratio between tip cell position and capillary in *amot*$^{ec+}$ and *amot*$^{ec-}$ retinas. All *amot*$^{ec+}$ retinas (n=25) and *amot*$^{ec-}$ retinas (n=15) were analyzed for statistical difference in (B). Retinas with recombination frequencies of 0-50% at capillaries (*amot*$^{ec+}$ n=13; *amot*$^{ec-}$ n=6) were analyzed in (C), and high recombination group (85-100%) was compared in (D) (*amot*$^{ec+}$ n=8; *amot*$^{ec-}$ n=3). For each sample, at least three representative images were taken from tip cell and capillary area, respectively. *n.s*, not significant, $^{**}P < 0.01$ and $^{***}P < 0.001$.

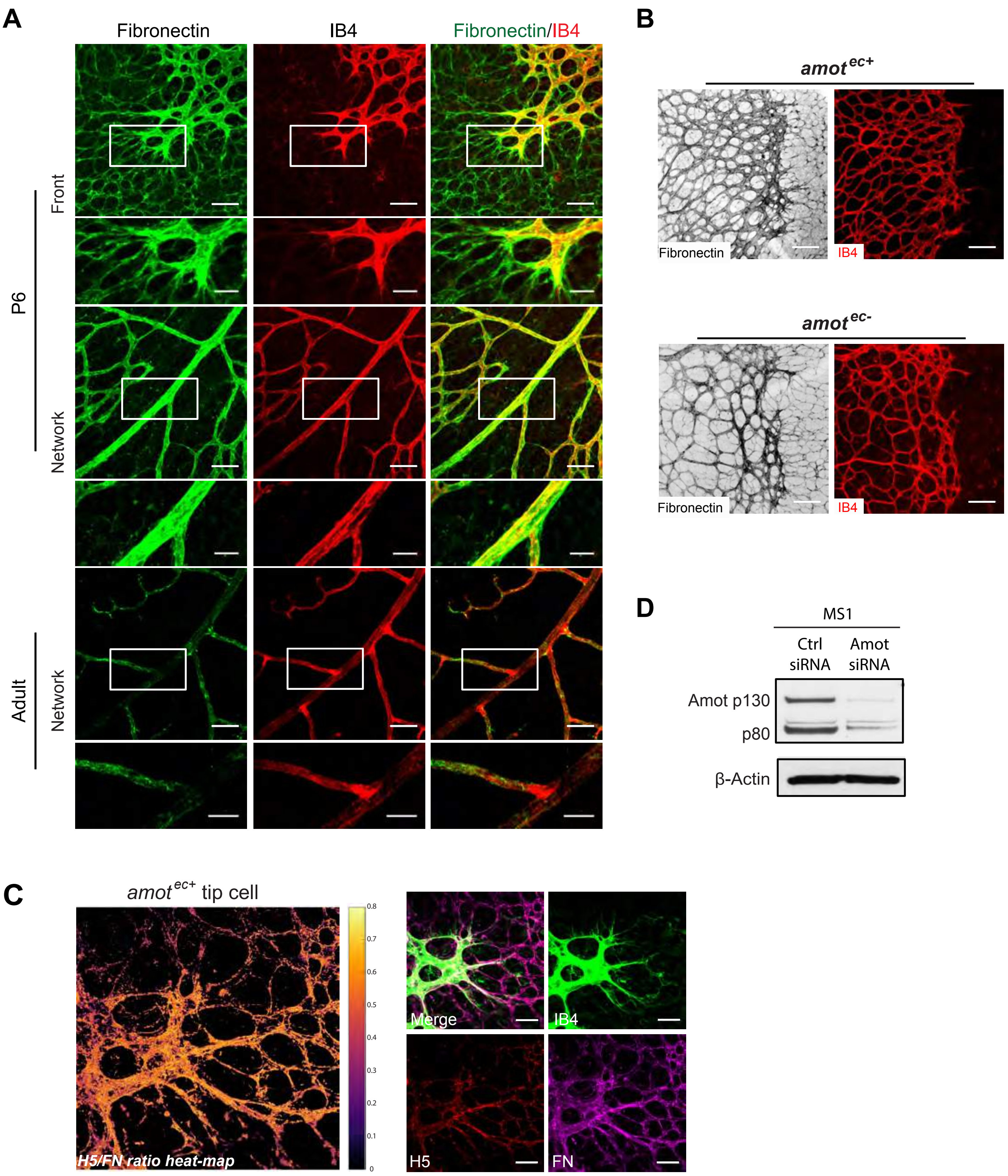
A
Fibronectin
IB4
Fibronectin/IB4
P6
Front
Network
Adult
Network
B
amot^ec+
Fibronectin
IB4
amot^ec-
Fibronectin
IB4
D
MS1
Ctrl siRNA
Amot siRNA
Amot p130
p80
β-Actin
C
amot^ec+ tip cell
H5/FN ratio heat-map
0.8
0.7
0.6
0.5
0.4
0.3
0.2
0.1
0
Merge
IB4
H5
FN

**Figure S4. Amot relays force between EC and Fn.**

(A) Fibronectin (in green) in wild-type retinas from P6 and adult age was visualized by IF staining. Blood vessels were visualized by IB4 staining (in red). Images of vascular fronts and capillaries at P6 are positioned in the upper and middle panels, respectively, while images from adult retina were placed at the bottom panel. White boxes indicate areas shown in higher magnification. (B) Fn expression pattern in $amot^{ec+}$ and $amot^{ec-}$ retinas at P6. Retinas were whole-mount stained with Fn (in grayscale) and IB4 (in red). (C) Heatmap showing the ratio of H5/Fn fluorescent intensity at $amot^{ec+}$ retina at P6. Retinal fluorescent images were stained with H5-myc (in red) showing strained Fn distribution, pan-Fn (in magenta) and IB4 (in green). (D) WB verification of Amot siRNA depletion in MS1 cells. Scale bars, (A) 50 µm, and 25 µm in the magnified images, (B) 100 µm, (C) 50 µm.

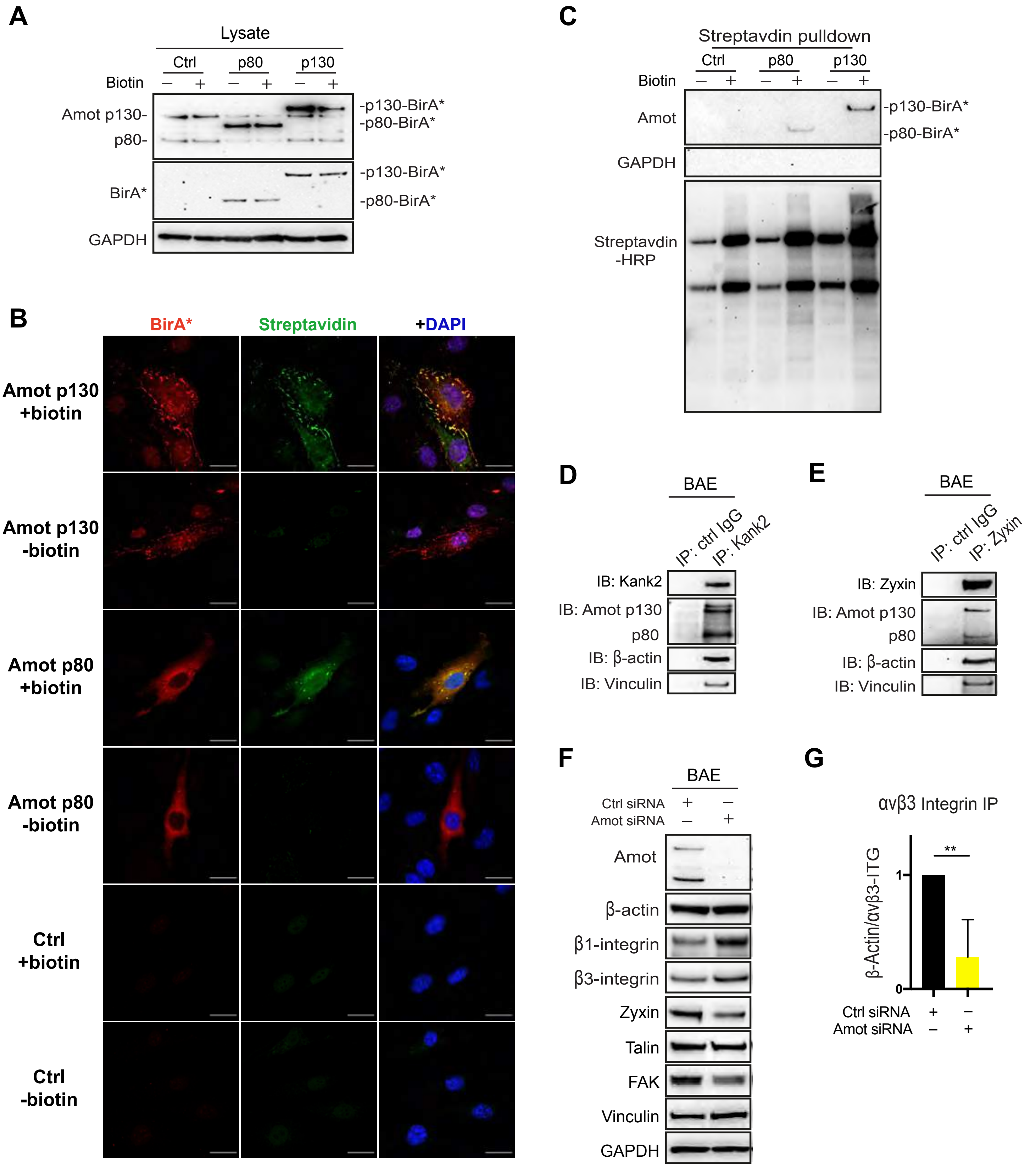
A
Lysate
Ctrl
p80
p130
Biotin
Amot p130-
p80-
-p130-BirA*
-p80-BirA*
BirA*
GAPDH
B
BirA*
Streptavidin
+DAPI
Amot p130 +biotin
Amot p130 -biotin
Amot p80 +biotin
Amot p80 -biotin
Ctrl +biotin
Ctrl -biotin
C
Streptavdin pulldown
Amot
GAPDH
Streptavdin -HRP
D
BAE
IP: ctrl IgG
IP: Kank2
IB: Kank2
IB: Amot p130
p80
IB: β-actin
IB: Vinculin
E
IP: Zyxin
IB: Zyxin
F
Ctrl siRNA
Amot siRNA
β1-integrin
β3-integrin
Zyxin
Talin
FAK
Vinculin
G
αvβ3 Integrin IP
β-Actin/αvβ3-ITG
**

**Figure S5. Amot promotes actin filament formation at focal adhesions**

(A-C) WB analysis BioID-fusion protein expression in transfected MS endothelial cells, including empty vector, Amot-p80 BioID or Amot-p130 BioID. Biotin ligase BirA is activated by 16-hour incubation of of 20nM biotin. (B) IF staining of BirA and Streptavidin in transfected MS1 cells. Scale bar, 20 μm. (C) WB analysis of IP of using streptavidin sepharose, probed with antibodies against Amot and streptavidin-HRP. GAPDH was included as a non-binding negative control. (D-E) WB verification Amot and kank2 (D) and zyxin (E) interaction by Co-IP in sub-confluent BAE cells (40%). Rabbit IgG was included as a negative control. (F) Steady-state levels of expression of focal adhesion related molecules in control or Amot siRNA depleted cells as analyzed by WB. The lysates were harvested from the BAE cells transfected by control or Amot siRNA. (G) Quantification of the intensity of actin using ImageJ in αvβ3-integrin IP samples. The quantities of β-actin were normalized to expression of αvβ3-integrin. All data above were collected from at least three independent experiments. $^{**}P < 0.01$.

**Figure S6. Full length blots to Figure 2B and 6A.**

**Figure S7. Full length blots to Figure 6B, 6C and 6D.**

**Figure S8. Full length blots to Figure S5A, S5C, S5D and S5E.**

**Figure S9. Full length blots to Figure S2A, S4D and S5F.**

**Figure 2B**

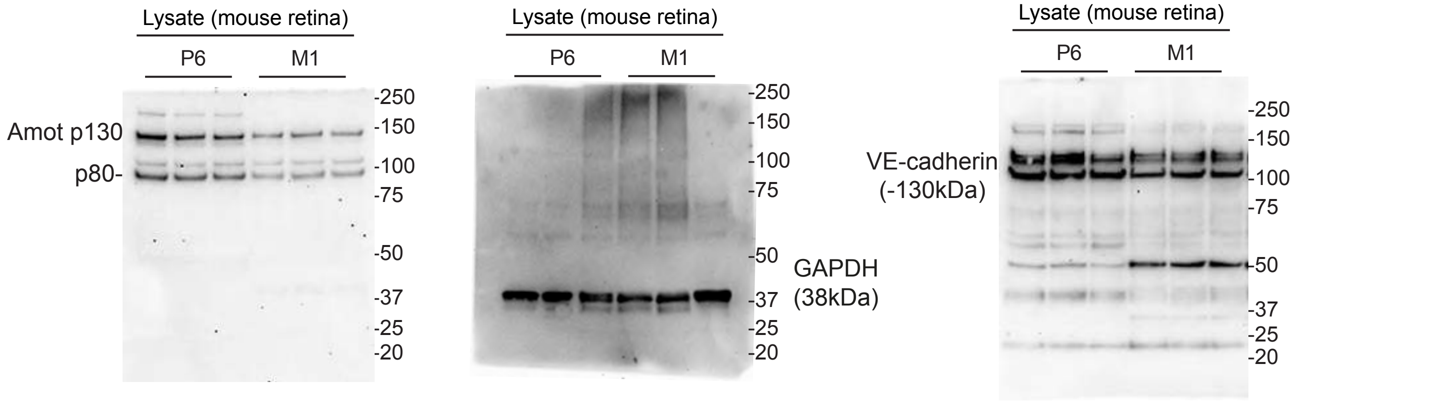

**Figure 6A** All in BAE cells

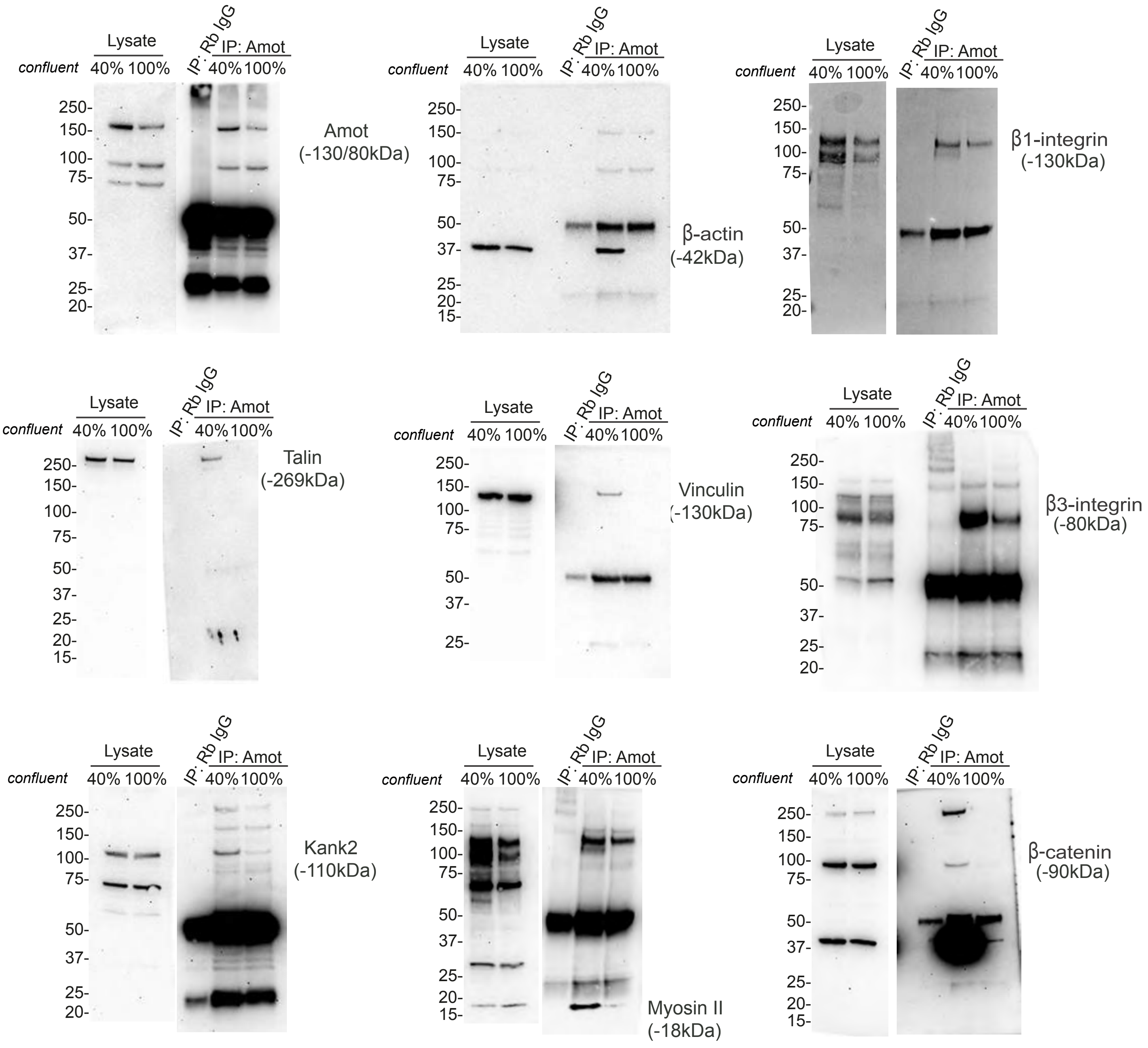

**Figure S6.** Full length blots to Figure 2B and 6A.

All in BAE cells
Note: ITG=integrin

**Figure 6B**

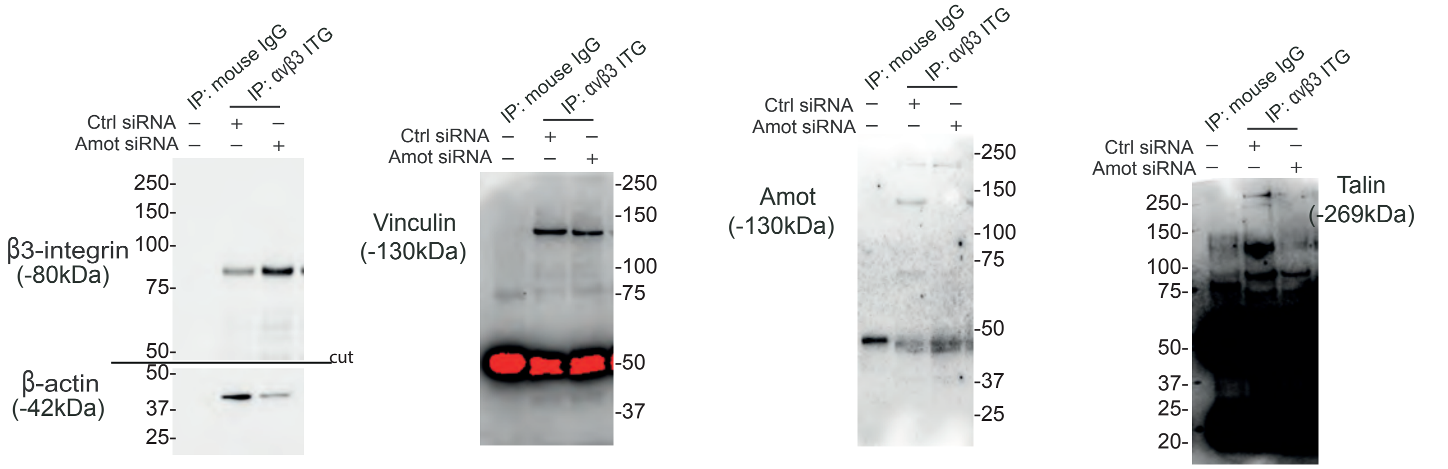

**Figure 6C**

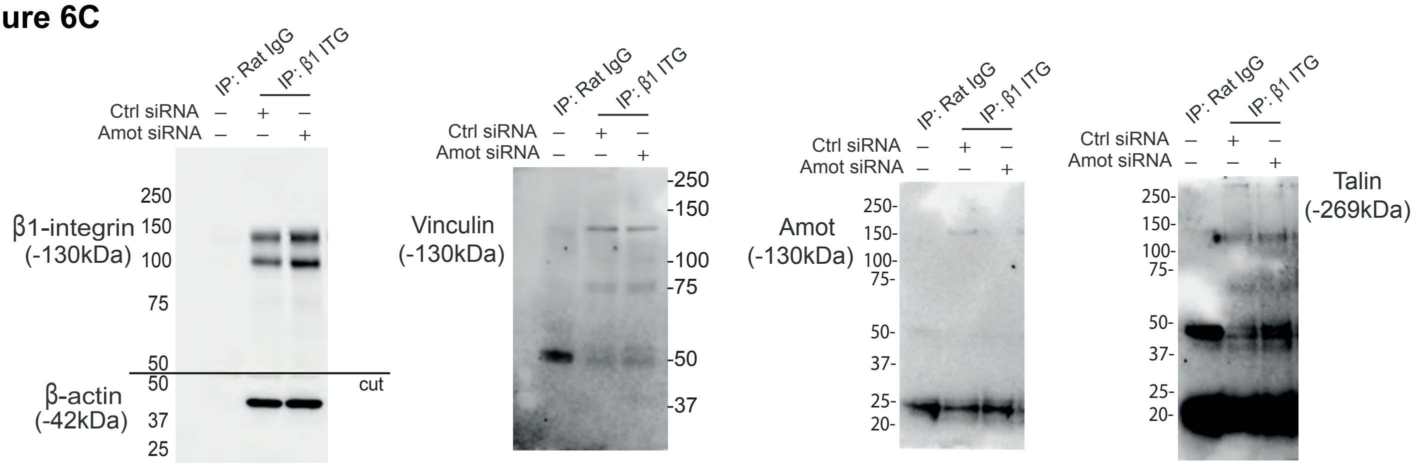

**Figure 6D**

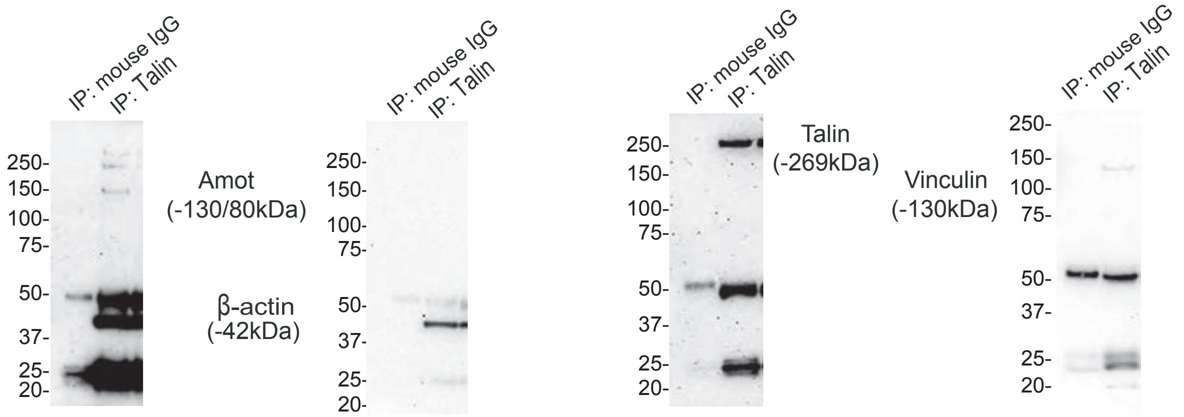

**Figure S7.** Full length blots to Figure 6B, 6C and 6D.

**Figure S5A**

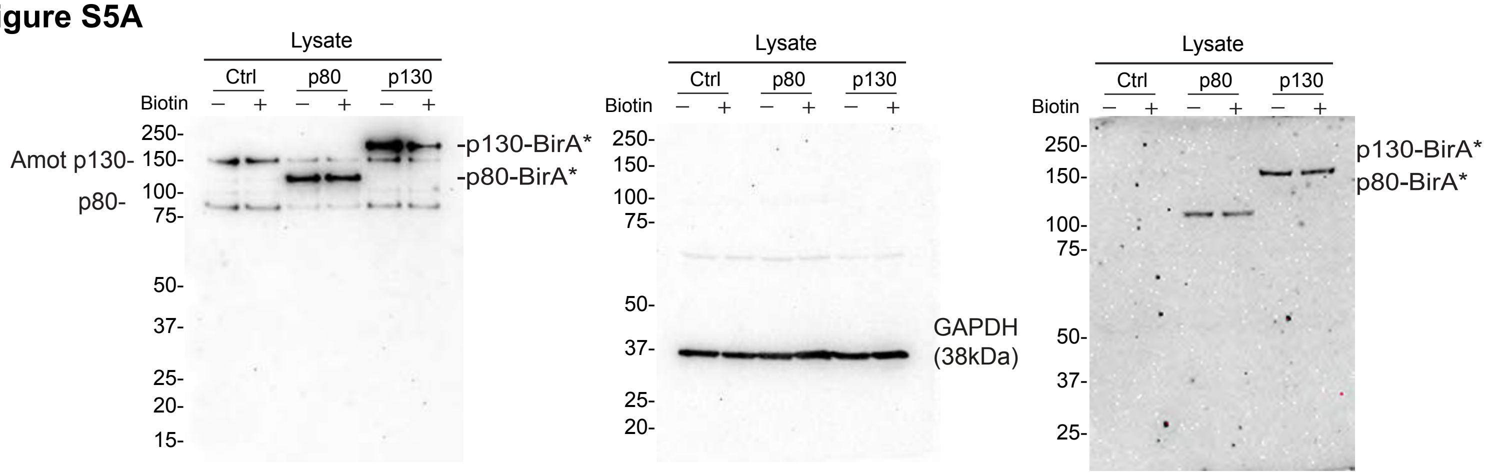

**Figure S5C**

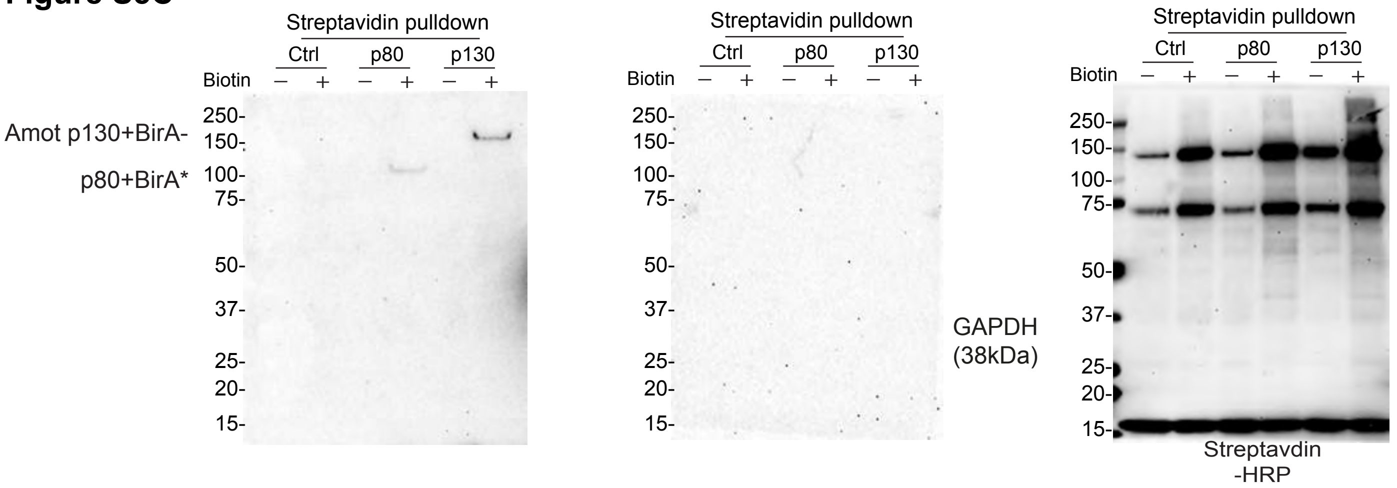

**Figure S5D**

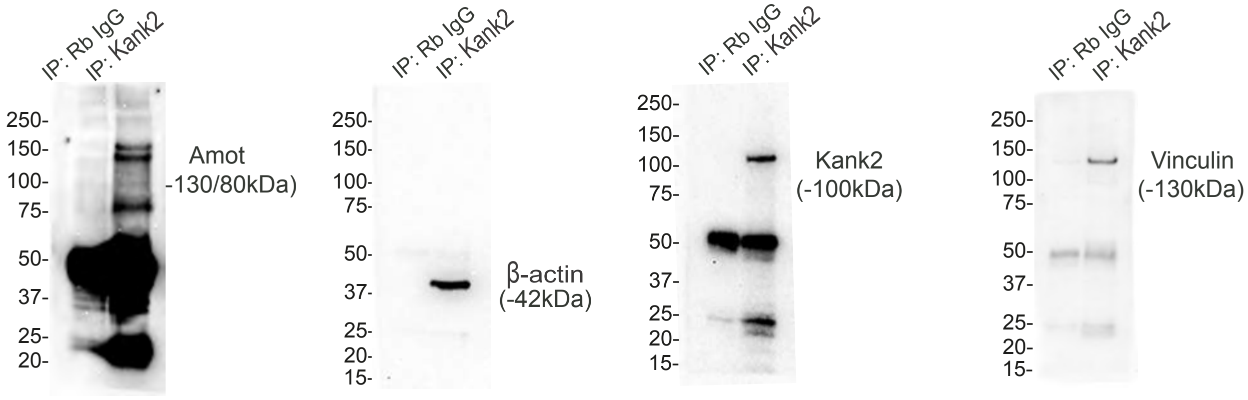

**Figure S5E**

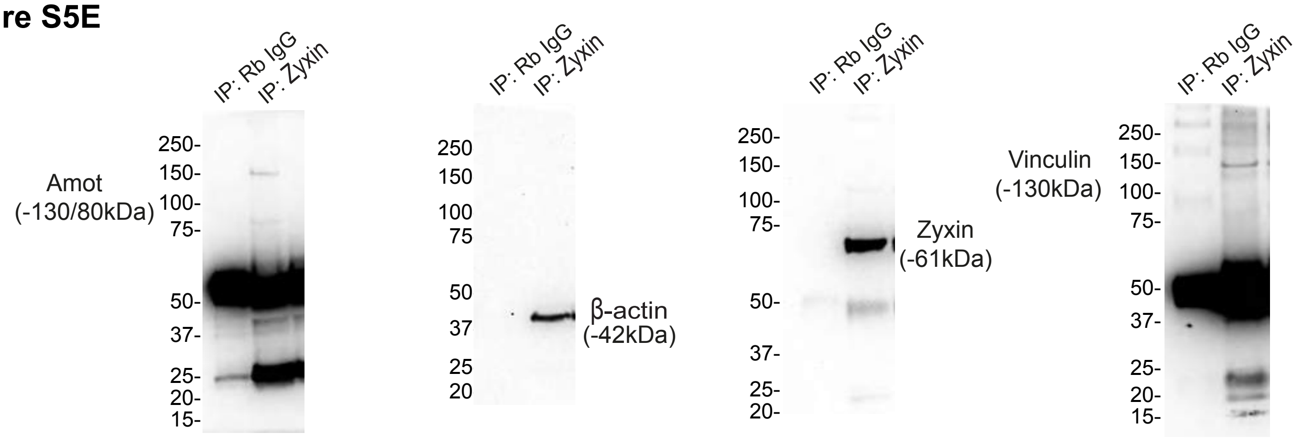

**Figure S8.** Full length blots to Figure S5A, S5C, S5D, and S5E.

**Figure S2A**

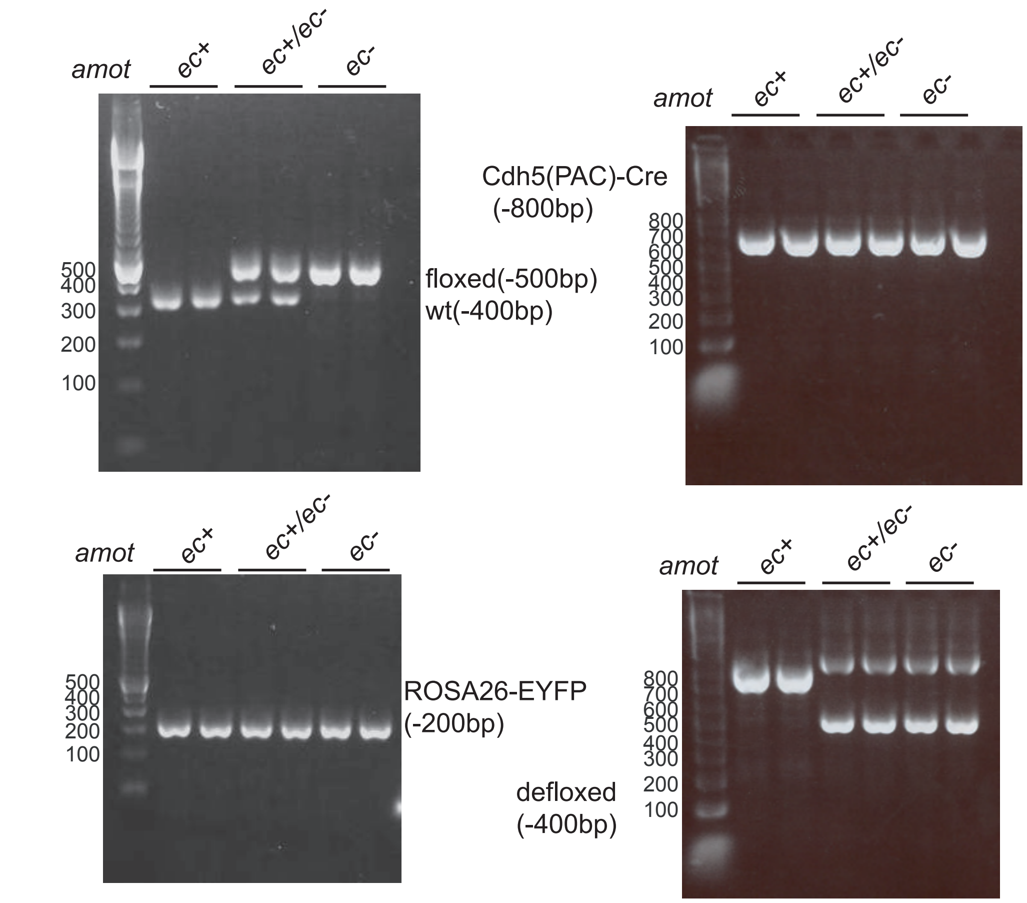

**Figure S4D**

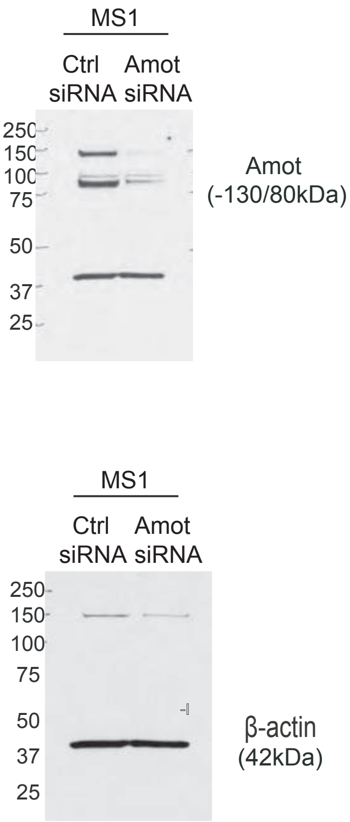

**Figure S5F**

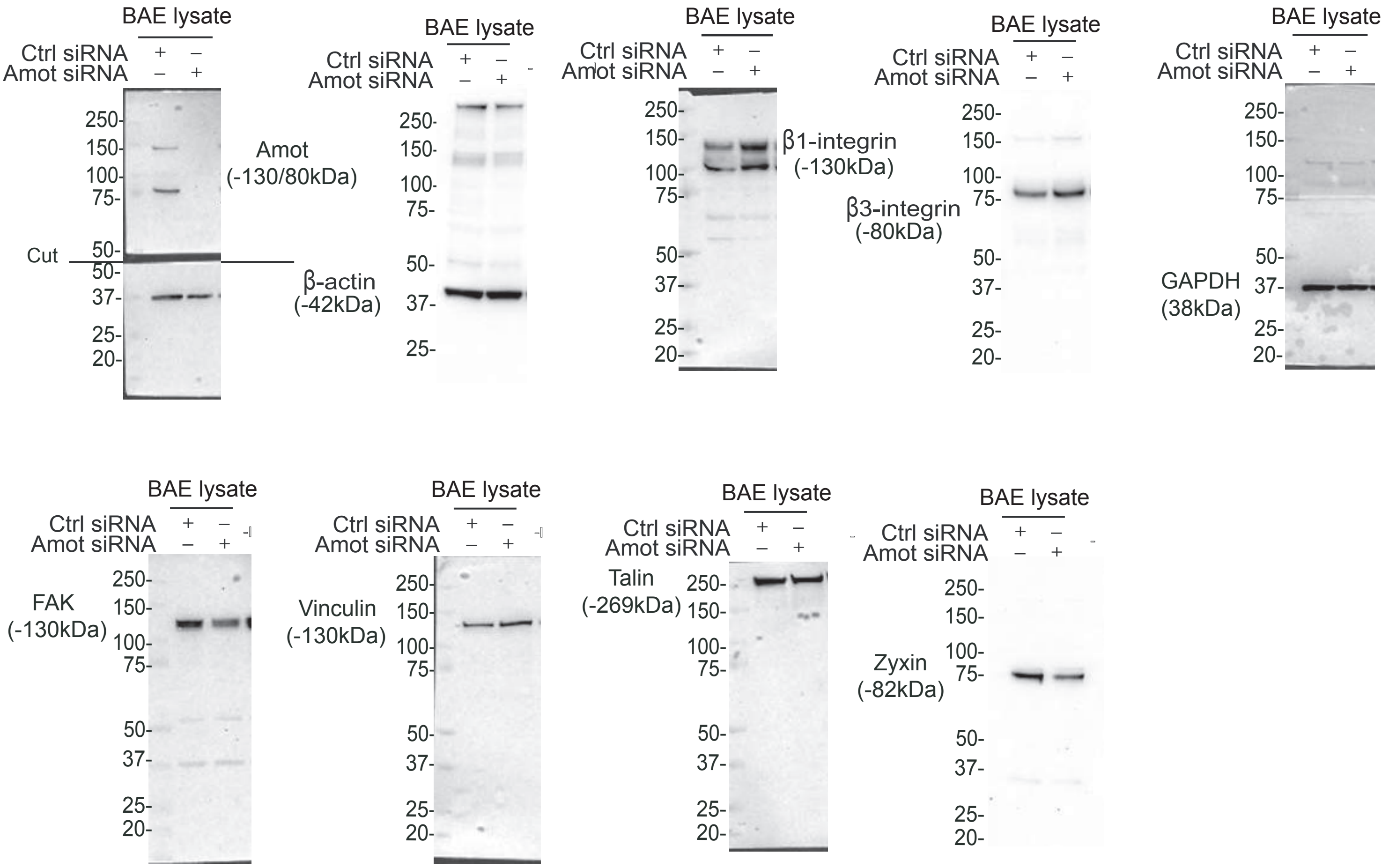

**Figure S9.** Full length blots to Figure S2A, S4D and S5F.

### Table S1

| | Accession | Description | Area_Ctrl_1 | Area_Ctrl_2 | Area_Ctrl_3 | Area_Amot_1 | Area_Amot_2 | Gene_Symbol | control.mean | amot.mean | fold change (fc) | log10 fc |
|---|---|---|---|---|---|---|---|---|---|---|---|---|
| 1 | F1N6C0 | Uncharacterized protein OS=Bos taurus OX=9913 GN=CALM2 PE=4 SV=2 - [F1N6C0_BOVIN] | 0 | 0 | 0 | 5895271043 | 33870539 | CALM2 | 1 | 2964570791 | 2964570791 | 9,471961825 |
| 2 | Q1RMT6 | Drebrin 1 OS=Bos taurus OX=9913 GN=DBN1 PE=1 SV=1 - [Q1RMT6_BOVIN] | 0 | 0 | 0 | 4118044519 | 29792249,5 | DBN1 | 1 | 2073918384 | 2073918384 | 9,316791661 |
| 3 | Q5KR47-2 | Isoform 2 of Tropomyosin alpha-3 chain OS=Bos taurus OX=9913 GN=TPM3 - [TPM3_BOVIN] | 0 | 0 | 0 | 3072624208 | 11039789,75 | TPM3 | 1 | 1541831999 | 1541831999 | 9,188037055 |
| 4 | Q5KR47 | Tropomyosin alpha-3 chain OS=Bos taurus OX=9913 GN=TPM3 PE=2 SV=1 - [TPM3_BOVIN] | 0 | 0 | 0 | 2640094631 | 11039789,75 | TPM3 | 1 | 1325567211 | 1325567211 | 9,122401753 |
| 5 | G3MWI5 | Coronin OS=Bos taurus OX=9913 GN=CORO1C PE=3 SV=1 - [G3MWI5_BOVIN] | 0 | 0 | 0 | 2522559255 | 48670426,89 | CORO1C | 1 | 1285614841 | 1285614841 | 9,109110877 |
| 6 | P79136-2 | rm Beta-2 of F-actin-capping protein subunit beta OS=Bos taurus OX=9913 GN=CAPZB - [CAPZB_BC | 0 | 0 | 0 | 1446576471 | 6211089,75 | CAPZB | 1 | 726393780 | 726393780,5 | 8,861172117 |
| 7 | F1N2Z6 | led-coil alpha-helical rod protein 1 OS=Bos taurus OX=9913 GN=CCHCR1 PE=4 SV=1 - [F1N2Z6_BOV | 0 | 0 | 0 | 717734700 | 538665858,5 | CCHCR1 | 1 | 628200279 | 628200279,3 | 8,798098125 |
| 8 | Q2KJH0 | Tropomodulin 3 (Ubiquitous) OS=Bos taurus OX=9913 GN=TMOD3 PE=2 SV=1 - [Q2KJH0_BOVIN] | 0 | 0 | 0 | 1233408587 | 12048472,5 | TMOD3 | 1 | 622728530 | 622728529,5 | 8,794298763 |
| 9 | A4FUA8 | ctin-capping protein subunit alpha-1 OS=Bos taurus OX=9913 GN=CAPZA1 PE=2 SV=1 - [CAZA1_BO\ | 0 | 0 | 0 | 1114793077 | 36771813,92 | CAPZA1 | 1 | 575782445 | 575782445,5 | 8,76025842 |
| 10 | F1MSZ6 | Antithrombin-III OS=Bos taurus OX=9913 GN=SERPINC1 PE=3 SV=1 - [F1MSZ6_BOVIN] | 0 | 0 | 0 | 745030280,8 | 144866800,3 | SERPINC1 | 1 | 444948541 | 444948540,5 | 8,648309787 |
| 11 | Q5E997 | ctin-capping protein subunit alpha-2 OS=Bos taurus OX=9913 GN=CAPZA2 PE=1 SV=3 - [CAZA2_BO\ | 0 | 0 | 0 | 772069758,9 | 12332887,06 | CAPZA2 | 1 | 392201323 | 392201323 | 8,593509054 |
| 12 | B8Y9S9 | yo-specific fibronectin 1 transcript variant OS=Bos taurus OX=9913 GN=FN1 PE=2 SV=1 - [B8Y9S9_B | 0 | 0 | 0 | 594656739,9 | 93541827,75 | FN1 | 1 | 344099284 | 344099283,8 | 8,536683769 |
| 13 | E1BF59 | Plectin OS=Bos taurus OX=9913 GN=PLEC PE=1 SV=2 - [E1BF59_BOVIN] | 0 | 0 | 0 | 582237422,4 | 49418721,33 | PLEC | 1 | 315828072 | 315828071,9 | 8,499450729 |
| 14 | Q56JX6 | 40S ribosomal protein S28 OS=Bos taurus OX=9913 GN=RPS28 PE=3 SV=1 - [RS28_BOVIN] | 0 | 0 | 0 | 527797550,9 | 94036031,06 | RPS28 | 1 | 310916791 | 310916791 | 8,492644177 |
| 15 | F1MQ65 | Myosin IE OS=Bos taurus OX=9913 GN=MYO1E PE=3 SV=2 - [F1MQ65_BOVIN] | 0 | 0 | 0 | 592275008,1 | 14092709,78 | MYO1E | 1 | 303183859 | 303183859 | 8,481706076 |
| 16 | Q32PB8 | 40S ribosomal protein S21 OS=Bos taurus OX=9913 GN=RPS21 PE=3 SV=1 - [RS21_BOVIN] | 0 | 0 | 0 | 469405122,3 | 130666775 | RPS21 | 1 | 300035949 | 300035948,6 | 8,477173293 |
| 17 | E1BDU8 | LIM domain and actin binding 1 OS=Bos taurus OX=9913 GN=LIMA1 PE=1 SV=2 - [E1BDU8_BOVIN] | 0 | 0 | 0 | 583632615,3 | 2573143,906 | LIMA1 | 1 | 293102880 | 293102879,6 | 8,467020085 |
| 18 | F1MBQ8 | DEAD-box helicase 5 OS=Bos taurus OX=9913 GN=DDX5 PE=3 SV=2 - [F1MBQ8_BOVIN] | 0 | 0 | 0 | 185824344,3 | 362845158,8 | DDX5 | 1 | 274334752 | 274334751,5 | 8,438280826 |
| 19 | A4IFC3 | Polyadenylate-binding protein OS=Bos taurus OX=9913 GN=PABPC4 PE=2 SV=1 - [A4IFC3_BOVIN] | 0 | 0 | 0 | 400142617,6 | 121759127,7 | PABPC4 | 1 | 260950873 | 260950872,6 | 8,416558754 |
| 20 | P61284 | 60S ribosomal protein L12 OS=Bos taurus OX=9913 GN=RPL12 PE=2 SV=1 - [RL12_BOVIN] | 0 | 0 | 0 | 430148451,2 | 57560663,96 | RPL12 | 1 | 243854558 | 243854557,6 | 8,387130877 |
| 21 | Q3T0D5 | 60S ribosomal protein L30 OS=Bos taurus OX=9913 GN=RPL30 PE=3 SV=3 - [RL30_BOVIN] | 0 | 0 | 0 | 321344226,4 | 131409590,1 | RPL30 | 1 | 226376908 | 226376908,2 | 8,354832124 |
| 22 | Q1RMR3 | Cortactin OS=Bos taurus OX=9913 GN=CTTN PE=1 SV=1 - [Q1RMR3_BOVIN] | 0 | 0 | 0 | 441065882,5 | 5696908,172 | CTTN | 1 | 223381395 | 223381395,3 | 8,349046999 |
| 23 | A7E307 | DDX17 protein OS=Bos taurus OX=9913 GN=DDX17 PE=1 SV=1 - [A7E307_BOVIN] | 0 | 0 | 0 | 80729644,33 | 360955385 | DDX17 | 1 | 220842515 | 220842514,7 | 8,344082684 |
| 24 | Q76I81 | 40S ribosomal protein S12 OS=Bos taurus OX=9913 GN=RPS12 PE=2 SV=1 - [RS12_BOVIN] | 0 | 0 | 0 | 388161316,3 | 30865840 | RPS12 | 1 | 209513578 | 209513578,2 | 8,321212174 |
| 25 | Q95140 | 60S acidic ribosomal protein P0 OS=Bos taurus OX=9913 GN=RPLP0 PE=2 SV=3 - [RLA0_BOVIN] | 0 | 0 | 0 | 337047499 | 59162337,13 | RPLP0 | 1 | 198104918 | 198104918,1 | 8,296895257 |
| 26 | Q3ZBH8 | 40S ribosomal protein S20 OS=Bos taurus OX=9913 GN=RPS20 PE=3 SV=1 - [RS20_BOVIN] | 0 | 0 | 0 | 319653158,2 | 74049536,63 | RPS20 | 1 | 196851347 | 196851347,4 | 8,294138392 |
| 27 | Q3SYR7 | 60S ribosomal protein L9 OS=Bos taurus OX=9913 GN=RPL9 PE=2 SV=1 - [RL9_BOVIN] | 0 | 0 | 0 | 379611994 | 10917911,5 | RPL9 | 1 | 195264953 | 195264952,8 | 8,290624301 |
| 28 | P61356 | 60S ribosomal protein L27 OS=Bos taurus OX=9913 GN=RPL27 PE=2 SV=2 - [RL27_BOVIN] | 0 | 0 | 0 | 298844930,3 | 59216827,5 | RPL27 | 1 | 179030879 | 179030878,9 | 8,252927944 |
| 29 | P26452 | 40S ribosomal protein SA OS=Bos taurus OX=9913 GN=RPSA PE=2 SV=4 - [RSSA_BOVIN] | 0 | 0 | 0 | 299090004,9 | 51605463,75 | RPSA | 1 | 175347734 | 175347734,3 | 8,243900159 |
| 30 | F1N076 | Ceruloplasmin OS=Bos taurus OX=9913 GN=CP PE=1 SV=2 - [F1N076_BOVIN] | 0 | 0 | 0 | 307852167,8 | 35271451 | CP | 1 | 171561809 | 171561809,4 | 8,234420618 |
| 31 | Q32PD5 | 40S ribosomal protein S19 OS=Bos taurus OX=9913 GN=RPS19 PE=2 SV=3 - [RS19_BOVIN] | 0 | 0 | 0 | 293628380,5 | 48275897,38 | RPS19 | 1 | 170952139 | 170952138,9 | 8,232874539 |
| 32 | E1BC59 | phosphatase 1 regulatory subunit 12A OS=Bos taurus OX=9913 GN=PPP1R12A PE=4 SV=2 - [E1BC59 | 0 | 0 | 0 | 326633867,6 | 13397689,24 | PPP1R12A | 1 | 170015778 | 170015778,4 | 8,230489228 |
| 33 | P17697 | Clusterin OS=Bos taurus OX=9913 GN=CLU PE=1 SV=1 - [CLUS_BOVIN] | 0 | 0 | 0 | 172362175,5 | 148531286 | CLU | 1 | 160446731 | 160446730,8 | 8,205330872 |
| 34 | E1BCF8 | ger CCCH-type containing, antiviral 1 OS=Bos taurus OX=9913 GN=ZC3HAV1 PE=4 SV=1 - [E1BCF8_ | 0 | 0 | 0 | 300876474,6 | 17574557,63 | ZC3HAV1 | 1 | 159225516 | 159225516,1 | 8,202012665 |
| 35 | Q5E988 | 40S ribosomal protein S5 OS=Bos taurus OX=9913 GN=RPS5 PE=2 SV=3 - [RS5_BOVIN] | 0 | 0 | 0 | 270802420 | 32257671,16 | RPS5 | 1 | 151530046 | 151530045,6 | 8,180498754 |
| 36 | Q5E9E6 | 60S ribosomal protein L10a OS=Bos taurus OX=9913 GN=RPL10A PE=2 SV=3 - [RL10A_BOVIN] | 0 | 0 | 0 | 184011313,6 | 118676460,7 | RPL10A | 1 | 151343887 | 151343887,1 | 8,179964884 |
| 37 | O18789 | 40S ribosomal protein S2 OS=Bos taurus OX=9913 GN=RPS2 PE=2 SV=2 - [RS2_BOVIN] | 0 | 0 | 0 | 258973767,3 | 35476091,13 | RPS2 | 1 | 147224929 | 147224929,2 | 8,167981354 |
| 38 | P00735 | Prothrombin OS=Bos taurus OX=9913 GN=F2 PE=1 SV=2 - [THRB_BOVIN] | 0 | 0 | 0 | 213754285,1 | 74987805,06 | F2 | 1 | 144371045 | 144371045,1 | 8,1594801 |
| 39 | Q862K7 | Dynein light chain (Fragment) OS=Bos taurus OX=9913 PE=2 SV=1 - [Q862K7_BOVIN] | 0 | 0 | 0 | 68361048,25 | 217777150,8 | DYNLL | 1 | 143069100 | 143069099,5 | 8,155545844 |
| 40 | E1BDB1 | Ras interacting protein 1 OS=Bos taurus OX=9913 GN=RASIP1 PE=4 SV=1 - [E1BDB1_BOVIN] | 0 | 0 | 0 | 257013352 | 24438793,21 | RASIP1 | 1 | 140726073 | 140726072,6 | 8,148374568 |
| 41 | Q58DM9 | Ribosomal protein L27a OS=Bos taurus OX=9913 GN=RPL27A PE=2 SV=1 - [Q58DM9_BOVIN] | 0 | 0 | 0 | 273698407 | 5021160,5 | RPL27A | 1 | 139359784 | 139359783,8 | 8,144137464 |
| 42 | E1BIB4 | ogeneous nuclear ribonucleoprotein L OS=Bos taurus OX=9913 GN=HNRNPL PE=1 SV=2 - [E1BIB4_E | 0 | 0 | 0 | 123381043,4 | 150949363,7 | HNRNPL | 1 | 137165204 | 137165203,5 | 8,137243952 |
| 43 | Q58DT1 | 60S ribosomal protein L7 OS=Bos taurus OX=9913 GN=RPL7 PE=2 SV=1 - [RL7_BOVIN] | 0 | 0 | 0 | 236236978,3 | 33281894,28 | RPL7 | 1 | 134759436 | 134759436,3 | 8,129559186 |
| 44 | Q3ZBL0 | Small nuclear ribonucleoprotein G OS=Bos taurus OX=9913 GN=SNRPG PE=3 SV=1 - [RUXG_BOVIN] | 0 | 0 | 0 | 153055457,8 | 113707659,3 | SNRPG | 1 | 133381559 | 133381558,5 | 8,125095788 |
| 45 | A2VDN7 | ribonucleoprotein U (Scaffold attachment factor A) OS=Bos taurus OX=9913 GN=HNRNPU PE=2 S\ | 0 | 0 | 0 | 179554959,1 | 63358966,87 | HNRNPU | 1 | 121456963 | 121456963 | 8,084422417 |
| 46 | Q56JX3 | 60S ribosomal protein L31 OS=Bos taurus OX=9913 GN=RPL31 PE=2 SV=1 - [RL31_BOVIN] | 0 | 0 | 0 | 215433439,7 | 21930357,13 | RPL31 | 1 | 118681898 | 118681898,4 | 8,074384485 |
| 47 | Q56JZ1 | 60S ribosomal protein L13 OS=Bos taurus OX=9913 GN=RPL13 PE=2 SV=3 - [RL13_BOVIN] | 0 | 0 | 0 | 203113727,6 | 31128950,5 | RPL13 | 1 | 117121339 | 117121339,1 | 8,068636029 |
| 48 | Q76I82 | 40S ribosomal protein S15a OS=Bos taurus OX=9913 GN=RPS15A PE=2 SV=1 - [RS15A_BOVIN] | 0 | 0 | 0 | 188533262,9 | 26571723,41 | RPS15A | 1 | 107552493 | 107552493,2 | 8,031620482 |
| 49 | Q56JU9-2 | Isoform 2 of 40S ribosomal protein S24 OS=Bos taurus OX=9913 GN=RPS24 - [RS24_BOVIN] | 0 | 0 | 0 | 199535632 | 9332225 | RPS24 | 1 | 104433929 | 104433928,5 | 8,018841615 |
| 50 | Q6LBN7 | Lactoferrin (Fragment) OS=Bos taurus OX=9913 PE=2 SV=1 - [Q6LBN7_BOVIN] | 0 | 0 | 0 | 188729027,8 | 19730122,19 | LF | 1 | 104229575 | 104229575 | 8,017990967 |
| 51 | Q58DW5 | 60S ribosomal protein L5 OS=Bos taurus OX=9913 GN=RPL5 PE=2 SV=3 - [RL5_BOVIN] | 0 | 0 | 0 | 196811230,4 | 8539606 | RPL5 | 1 | 102675418 | 102675418,2 | 8,01146648 |
| 52 | A6QPP2 | SERPIND1 protein OS=Bos taurus OX=9913 GN=SERPIND1 PE=2 SV=1 - [A6QPP2_BOVIN] | 0 | 0 | 0 | 83597258 | 115814540,9 | SERPIND1 | 1 | 99705899,4 | 99705899,44 | 7,998720856 |
| 53 | F1N690 | e component of pyruvate dehydrogenase complex OS=Bos taurus OX=9913 GN=DLAT PE=1 SV=1 - [ | 0 | 0 | 0 | 103916441,5 | 81256895,75 | DLAT | 1 | 92586668,6 | 92586668,63 | 7,966548458 |
| 54 | E1BK63 | Ribosomal protein L15 OS=Bos taurus OX=9913 PE=3 SV=1 - [E1BK63_BOVIN] | 0 | 0 | 0 | 180090424 | 3237253,063 | RPL15 | 1 | 91663838,5 | 91663838,53 | 7,96219804 |
| 55 | Q0V8H6 | Putative helicase MOV-10 OS=Bos taurus OX=9913 GN=MOV10 PE=2 SV=1 - [MOV10_BOVIN] | 0 | 0 | 0 | 132513128,3 | 37334636,74 | MOV10 | 1 | 84923882,5 | 84923882,52 | 7,929029841 |
| 56 | Q56K10 | 40S ribosomal protein S15 OS=Bos taurus OX=9913 GN=RPS15 PE=2 SV=3 - [RS15_BOVIN] | 0 | 0 | 0 | 91096662,98 | 76354824,94 | RPS15 | 1 | 83725744 | 83725743,96 | 7,922859015 |
| 57 | P15246 | oaspartate(D-aspartate) O-methyltransferase OS=Bos taurus OX=9913 GN=PCMT1 PE=1 SV=2 - [PII | 0 | 0 | 0 | 116760015,9 | 50146912,95 | PCMT1 | 1 | 83453464,4 | 83453464,41 | 7,92144437 |
| 58 | Q56JV9 | 40S ribosomal protein S3a OS=Bos taurus OX=9913 GN=RPS3A PE=2 SV=3 - [RS3A_BOVIN] | 0 | 0 | 0 | 128614803,5 | 36054471,1 | RPS3A | 1 | 82334637,3 | 82334637,3 | 7,915582577 |
| 59 | A1L532 | us nuclear ribonucleoprotein A0 (Fragment) OS=Bos taurus OX=9913 GN=HNRPA0 PE=2 SV=1 - [A1I | 0 | 0 | 0 | 79312284,75 | 75665820,21 | HNRPA0 | 1 | 77489052,5 | 77489052,48 | 7,88924035 |
| 60 | F1N301 | Uncharacterized protein OS=Bos taurus OX=9913 GN=RPL22 PE=1 SV=2 - [F1N301_BOVIN] | 0 | 0 | 0 | 140693632,6 | 13860380,59 | RPL22 | 1 | 77277006,6 | 77277006,61 | 7,888050291 |
| 61 | E1BAK6 | DAZ associated protein 1 OS=Bos taurus OX=9913 GN=DAZAP1 PE=4 SV=2 - [E1BAK6_BOVIN] | 0 | 0 | 0 | 92822204 | 59616712,75 | DAZAP1 | 1 | 76219458,4 | 76219458,38 | 7,882065858 |
| 62 | F1MBG0 | r associated arginine methyltransferase 1 OS=Bos taurus OX=9913 GN=CARM1 PE=3 SV=2 - [F1MB( | 0 | 0 | 0 | 73633889,5 | 75216610,19 | CARM1 | 1 | 74425249,8 | 74425249,84 | 7,871720301 |
| 63 | F1N169 | Filamin A OS=Bos taurus OX=9913 GN=FLNA PE=1 SV=2 - [F1N169_BOVIN] | 0 | 0 | 0 | 133763768,6 | 7454906,375 | FLNA | 1 | 70609337,5 | 70609337,48 | 7,848862137 |
| 64 | 0A1B0Z5 | Heat shock 27 kDa protein 1 OS=Bos taurus OX=9913 GN=HspB1 PE=3 SV=1 - [A0A1B0Z542_BOVIN | 0 | 0 | 0 | 73217856,4 | 66530829,1 | HspB1 | 1 | 69874342,7 | 69874342,75 | 7,844317736 |
| 65 | E1BNQ9 | Uncharacterized protein OS=Bos taurus OX=9913 GN=HNRNPUL2 PE=1 SV=1 - [E1BNQ9_BOVIN] | 0 | 0 | 0 | 132073857,5 | 4103695,313 | HNRNPUL2 | 1 | 68088776,4 | 68088776,41 | 7,83307553 |
| 66 | P39872 | 60S ribosomal protein L3 OS=Bos taurus OX=9913 GN=RPL3 PE=2 SV=2 - [RL3_BOVIN] | 0 | 0 | 0 | 75747932,33 | 52634673,38 | RPL3 | 1 | 64191302,9 | 64191302,85 | 7,80747619 |
| 67 | Q32LC7 | Pase-activating protein-binding protein 1 OS=Bos taurus OX=9913 GN=G3BP1 PE=2 SV=1 - [G3BP1_ | 0 | 0 | 0 | 114265111 | 14025739 | G3BP1 | 1 | 64145425 | 64145425,01 | 7,807165687 |
| 68 | Q7YRV4 | 3 ubiquitin-protein ligase TRIM21 OS=Bos taurus OX=9913 GN=TRIM21 PE=2 SV=1 - [RO52_BOVIN | 0 | 0 | 0 | 59693725,46 | 67800647,01 | TRIM21 | 1 | 63747186,2 | 63747186,23 | 7,80446102 |
| 69 | A7YWC4 | family AAA domain-containing protein 3 OS=Bos taurus OX=9913 GN=ATAD3 PE=2 SV=1 - [ATAD3_ | 0 | 0 | 0 | 79515453,24 | 47495991,4 | ATAD3 | 1 | 63505722,3 | 63505722,32 | 7,80281286 |
| 70 | F1MUZ9 | Da heat shock protein, mitochondrial OS=Bos taurus OX=9913 GN=HSPD1 PE=1 SV=1 - [F1MUZ9_BC | 0 | 0 | 0 | 63688259,71 | 54148201,58 | HSPD1 | 1 | 58918230,6 | 58918230,65 | 7,770249696 |
| 71 | P61287 | -protein phosphatase PP1-gamma catalytic subunit OS=Bos taurus OX=9913 GN=PPP1CC PE=2 SV=1 | 0 | 0 | 0 | 109838856 | 4347077,469 | PPP1CC | 1 | 57092966,7 | 57092966,73 | 7,756582611 |
| 72 | A6QQF3 | TJP2 protein OS=Bos taurus OX=9913 GN=TJP2 PE=2 SV=1 - [A6QQF3_BOVIN] | 0 | 0 | 0 | 76485954,63 | 36447794,46 | TJP2 | 1 | 56466874,5 | 56466874,54 | 7,75179375 |
| 73 | Q5E949 | nRNP methyltransferase-like 2 isoform 3 OS=Bos taurus OX=9913 GN=PRMT1 PE=2 SV=1 - [Q5E949_ | 0 | 0 | 0 | 78967238,31 | 33044272,39 | PRMT1 | 1 | 56005755,3 | 56005755,35 | 7,748232659 |
| 74 | E1BKT0 | Leucine zipper protein 1 OS=Bos taurus OX=9913 GN=LUZP1 PE=4 SV=2 - [E1BKT0_BOVIN] | 0 | 0 | 0 | 83997015,66 | 27541888,29 | LUZP1 | 1 | 55769452 | 55769451,98 | 7,746396377 |
| 75 | P13909 | Plasminogen activator inhibitor 1 OS=Bos taurus OX=9913 GN=SERPINE1 PE=1 SV=1 - [PAI1_BOVIN | 0 | 0 | 0 | 94008039,33 | 15333253,5 | SERPINE1 | 1 | 54670646,4 | 54670646,42 | 7,737754209 |
| 76 | Q58DW0 | 60S ribosomal protein L4 OS=Bos taurus OX=9913 GN=RPL4 PE=2 SV=3 - [RL4_BOVIN] | 0 | 0 | 0 | 94940626,75 | 12892108,48 | RPL4 | 1 | 53916367,6 | 53916367,62 | 7,731720626 |
| 77 | F1MBN2 | Tight junction protein 2 OS=Bos taurus OX=9913 GN=TJP2 PE=1 SV=2 - [F1MBN2_BOVIN] | 0 | 0 | 0 | 76485954,63 | 31000393,96 | TJP2 | 1 | 53743174,3 | 53743174,29 | 7,730323314 |
| 78 | Q1RMR2 | nall nuclear ribonucleoprotein 70 kDa OS=Bos taurus OX=9913 GN=SNRNP70 PE=2 SV=1 - [RU17_B( | 0 | 0 | 0 | 80737367,35 | 22839500,75 | SNRNP70 | 1 | 51788434,1 | 51788434,05 | 7,714232779 |
| 79 | Q862E3 | Similar to ribosomal protein L17 (Fragment) OS=Bos taurus OX=9913 PE=2 SV=1 - [Q862E3_BOVIN] | 0 | 0 | 0 | 55198525,53 | 47477485,81 | RPL17 | 1 | 51338005,7 | 51338005,67 | 7,710438994 |
| 80 | F1MZ00 | ll nuclear ribonucleoprotein Sm D3 OS=Bos taurus OX=9913 GN=SNRPD3 PE=3 SV=2 - [F1MZ00_BO | 0 | 0 | 0 | 79545112,41 | 23058348,82 | SNRPD3 | 1 | 51301730,6 | 51301730,61 | 7,710132016 |
| 81 | Q7M2Y2 | c transplantation antigen P198 homolog p23 (Fragment) OS=Bos taurus OX=9913 PE=3 SV=1 - [Q7M | 0 | 0 | 0 | 90702791,77 | 5488746,25 | TSTAP198 | 1 | 48095769 | 48095769,01 | 7,682106873 |
| 82 | Q3T0B6 | nent 1 Q subcomponent-binding protein, mitochondrial OS=Bos taurus OX=9913 GN=C1QBP PE=2 S\ | 0 | 0 | 0 | 28024823,58 | 68018568,75 | C1QBP | 1 | 48021696,2 | 48021696,17 | 7,681437496 |
| 83 | Q0QEV5 | ibosomal protein L18 (Fragment) OS=Bos taurus OX=9913 GN=RPL18 PE=2 SV=1 - [Q0QEV5_BOVIN | 0 | 0 | 0 | 81410737,81 | 13830464,25 | RPL18 | 1 | 47620601 | 47620601,03 | 7,677794872 |
| 84 | E1BEK9 | UPF1, RNA helicase and ATPase OS=Bos taurus OX=9913 GN=UPF1 PE=4 SV=2 - [E1BEK9_BOVIN] | 0 | 0 | 0 | 69394039,42 | 22976608,4 | UPF1 | 1 | 46185323,9 | 46185323,91 | 7,664503994 |
| 85 | F1MZU1 | nucleoprotein domain family member 1 OS=Bos taurus OX=9913 GN=LARP1 PE=4 SV=2 - [F1MZU1_ | 0 | 0 | 0 | 50165343 | 37373147,94 | LARP1 | 1 | 43769245,5 | 43769245,47 | 7,64116906 |
| 86 | Q3T0V4 | 40S ribosomal protein S11 OS=Bos taurus OX=9913 GN=RPS11 PE=2 SV=3 - [RS11_BOVIN] | 0 | 0 | 0 | 72453069 | 14609350,16 | RPS11 | 1 | 43531209,6 | 43531209,58 | 7,638800735 |
| 87 | A8E4M9 | PRPF39 protein OS=Bos taurus OX=9913 GN=PRPF39 PE=2 SV=1 - [A8E4M9_BOVIN] | 0 | 0 | 0 | 46350407,46 | 40548386,39 | PRPF39 | 1 | 43449396,9 | 43449396,92 | 7,637983753 |
| 88 | G3X757 | nsitional endoplasmic reticulum ATPase OS=Bos taurus OX=9913 GN=VCP PE=3 SV=1 - [G3X757_BO' | 0 | 0 | 0 | 47116721,33 | 39603695,83 | VCP | 1 | 43360208,6 | 43360208,58 | 7,637091363 |
| 89 | F1MYI2 | Uncharacterized protein OS=Bos taurus OX=9913 GN=HNRNPM PE=1 SV=2 - [F1MYI2_BOVIN] | 0 | 0 | 0 | 34316056,83 | 51646209,25 | HNRNPM | 1 | 42981133 | 42981133,04 | 7,63327786 |
| 90 | Q0IIK5 | ATP-dependent RNA helicase DDX1 OS=Bos taurus OX=9913 GN=DDX1 PE=2 SV=1 - [DDX1_BOVIN] | 0 | 0 | 0 | 73001823,25 | 12864306,45 | DDX1 | 1 | 42933064,8 | 42933064,85 | 7,632791892 |
| 91 | Q5E9T9 | tRNA-splicing ligase RtcB homolog OS=Bos taurus OX=9913 GN=RTCB PE=2 SV=1 - [RTCB_BOVIN] | 0 | 0 | 0 | 39838618,98 | 40603885,08 | RTCB | 1 | 40221252 | 40221252,03 | 7,604455585 |
| 92 | F1MD77 | Laminin subunit gamma 1 OS=Bos taurus OX=9913 GN=LAMC1 PE=1 SV=2 - [F1MD77_BOVIN] | 0 | 0 | 0 | 71458905,92 | 5555930,188 | LAMC1 | 1 | 38507418,1 | 38507418,05 | 7,5855444 |
| 93 | F1MY44 | Uncharacterized protein OS=Bos taurus OX=9913 GN=HNRNPM PE=1 SV=2 - [F1MY44_BOVIN] | 0 | 0 | 0 | 45002788 | 31416474,59 | HNRNPM | 1 | 38209631,3 | 38209631,3 | 7,582172847 |
| 94 | F1MKZ5 | 40S ribosomal protein S6 OS=Bos taurus OX=9913 PE=3 SV=1 - [F1MKZ5_BOVIN] | 0 | 0 | 0 | 62191729,77 | 11172503,81 | RPS6 | 1 | 36682116,8 | 36682116,79 | 7,564454389 |
| 95 | E3SAZ8 | Nucleophosmin OS=Bos taurus OX=9913 GN=NPM1 PE=4 SV=1 - [E3SAZ8_BOVIN] | 0 | 0 | 0 | 65024704,61 | 8262945 | NPM1 | 1 | 36643824,8 | 36643824,8 | 7,564000798 |
| 96 | A0JN82 | rbin and SH3 domain containing 2 OS=Bos taurus OX=9913 GN=SORBS2 PE=2 SV=1 - [A0JN82_BOVI | 0 | 0 | 0 | 50631549 | 20452867 | SORBS2 | 1 | 35542208 | 35542208 | 7,550744404 |
| 97 | F1MKX7 | 5'-3' exoribonuclease OS=Bos taurus OX=9913 GN=XRN2 PE=3 SV=1 - [F1MKX7_BOVIN] | 0 | 0 | 0 | 22702570,78 | 46243599,5 | XRN2 | 1 | 34473085,1 | 34473085,14 | 7,537480152 |
| 98 | F1ME65 | Cytoskeleton associated protein 4 OS=Bos taurus OX=9913 GN=CKAP4 PE=1 SV=1 - [F1ME65_BOVIN | 0 | 0 | 0 | 63723017,55 | 3521373,492 | CKAP4 | 1 | 33622195,5 | 33622195,52 | 7,526626069 |
| 99 | G3N3U9 | bosomal protein S22, mitochondrial OS=Bos taurus OX=9913 GN=MRPS22 PE=4 SV=1 - [G3N3U9_B | 0 | 0 | 0 | 39705586,33 | 27066282,84 | MRPS22 | 1 | 33385934,6 | 33385934,59 | 7,523563538 |
| 100 | F1MXX0 | MICOS complex subunit MIC60 OS=Bos taurus OX=9913 GN=IMMT PE=1 SV=1 - [F1MXX0_BOVIN] | 0 | 0 | 0 | 61275768,96 | 5400903,594 | IMMT | 1 | 33338336,3 | 33338336,28 | 7,522943923 |
| 101 | Q3T171 | 60S ribosomal protein L36 OS=Bos taurus OX=9913 GN=RPL36 PE=3 SV=3 - [RL36_BOVIN] | 0 | 0 | 0 | 52345235,97 | 13403826,88 | RPL36 | 1 | 32874531,4 | 32874531,42 | 7,516859571 |
| 102 | F1MTN1 | Integrin beta OS=Bos taurus OX=9913 GN=ITGB3 PE=3 SV=1 - [F1MTN1_BOVIN] | 0 | 0 | 0 | 58790171,96 | 5185097,021 | ITGB3 | 1 | 31987634,5 | 31987634,49 | 7,504982125 |
| 103 | Q5E9D3 | MICOS complex subunit MIC19 OS=Bos taurus OX=9913 GN=CHCHD3 PE=2 SV=1 - [MIC19_BOVIN] | 0 | 0 | 0 | 50431747,81 | 12569060,97 | CHCHD3 | 1 | 31500404,4 | 31500404,39 | 7,498316129 |
| 104 | Q0VCY7 | Serine/arginine-rich splicing factor 1 OS=Bos taurus OX=9913 GN=SRSF1 PE=2 SV=1 - [SRSF1_BOVIN | 0 | 0 | 0 | 52902313,11 | 10073178 | SRSF1 | 1 | 31487745,6 | 31487745,56 | 7,498141567 |
| 105 | Q3ZCE2 | ELAV-like protein OS=Bos taurus OX=9913 GN=ELAVL1 PE=2 SV=1 - [Q3ZCE2_BOVIN] | 0 | 0 | 0 | 39280992,73 | 21780716,34 | ELAVL1 | 1 | 30530854,5 | 30530854,54 | 7,48473896 |
| 106 | E1BP85 | Uridine-cytidine kinase OS=Bos taurus OX=9913 GN=UCKL1 PE=3 SV=1 - [E1BP85_BOVIN] | 0 | 0 | 0 | 16948066,62 | 43610834,88 | UCKL1 | 1 | 30279450,7 | 30279450,75 | 7,481147993 |
| 107 | Q3T122 | otic translation initiation factor 3 subunit D OS=Bos taurus OX=9913 GN=EIF3D PE=2 SV=1 - [EIF3D_ | 0 | 0 | 0 | 37112535,47 | 21458564,44 | EIF3D | 1 | 29285550 | 29285549,95 | 7,466653384 |
| 108 | F1N2D3 | Tight junction protein 1 OS=Bos taurus OX=9913 GN=TJP1 PE=3 SV=2 - [F1N2D3_BOVIN] | 0 | 0 | 0 | 46345329,1 | 11069774,49 | TJP1 | 1 | 28707551,8 | 28707551,8 | 7,457996157 |
| 109 | Q0VCV0 | Replication protein A subunit OS=Bos taurus OX=9913 GN=RPA1 PE=2 SV=1 - [Q0VCV0_BOVIN] | 0 | 0 | 0 | 34831646,75 | 22117340,5 | RPA1 | 1 | 28474493,6 | 28474493,63 | 7,45445601 |
| 110 | E1BPW2 | AB6-interacting/CAST family member 1 OS=Bos taurus OX=9913 GN=ERC1 PE=4 SV=2 - [E1BPW2_E | 0 | 0 | 0 | 38266563,71 | 18476640,75 | ERC1 | 1 | 28371602,2 | 28371602,23 | 7,452883862 |
| 111 | Q7YR75 | ribosomal protein L12, mitochondrial OS=Bos taurus OX=9913 GN=MRPL12 PE=1 SV=1 - [RM12_BO | 0 | 0 | 0 | 18513263,69 | 35557196,56 | MRPL12 | 1 | 27035230,1 | 27035230,13 | 7,431930071 |
| 112 | Q0VCD2 | Tubulin gamma-1 chain OS=Bos taurus OX=9913 GN=TUBG1 PE=2 SV=1 - [TBG1_BOVIN] | 0 | 0 | 0 | 41097990 | 12141068,68 | TUBG1 | 1 | 26619529,3 | 26619529,34 | 7,425200372 |
| 113 | A7MB80 | General transcription factor II-I OS=Bos taurus OX=9913 GN=GTF2I PE=2 SV=1 - [GTF2I_BOVIN] | 0 | 0 | 0 | 24785585,61 | 28359407,06 | GTF2I | 1 | 26572496,3 | 26572496,33 | 7,424432356 |
| 114 | E1BN58 | OTU deubiquitinase 4 OS=Bos taurus OX=9913 GN=OTUD4 PE=4 SV=2 - [E1BN58_BOVIN] | 0 | 0 | 0 | 28207550,88 | 22922927,35 | OTUD4 | 1 | 25565239,1 | 25565239,11 | 7,407649859 |
| 115 | F1MXX2 | gonaute 1, RISC catalytic component OS=Bos taurus OX=9913 GN=AGO1 PE=3 SV=1 - [F1MXX2_BO\ | 0 | 0 | 0 | 27267412,88 | 23711325,44 | AGO1 | 1 | 25489369,2 | 25489369,16 | 7,406359087 |
| 116 | E1B7R4 | tic translation initiation factor 3 subunit A OS=Bos taurus OX=9913 GN=EIF3A PE=3 SV=2 - [E1B7R4_ | 0 | 0 | 0 | 28923809,67 | 22041304,19 | EIF3A | 1 | 25482556,9 | 25482556,93 | 7,406243003 |
| 117 | A5D989 | Elongation factor 1-delta OS=Bos taurus OX=9913 GN=EEF1D PE=2 SV=2 - [EF1D_BOVIN] | 0 | 0 | 0 | 44234153,94 | 6390049,875 | EEF1D | 1 | 25312101,9 | 25312101,91 | 7,40332821 |
| 118 | G5E5N3 | ATP-dependent RNA helicase A OS=Bos taurus OX=9913 GN=DHX9 PE=4 SV=1 - [G5E5N3_BOVIN] | 0 | 0 | 0 | 23132777,58 | 26765027,94 | DHX9 | 1 | 24948902,8 | 24948902,76 | 7,39705145 |
| 119 | P68509 | 14-3-3 protein eta OS=Bos taurus OX=9913 GN=YWHAH PE=1 SV=2 - [1433F_BOVIN] | 0 | 0 | 0 | 22476427,69 | 27396156,31 | YWHAH | 1 | 24936292 | 24936292 | 7,396831875 |
| 120 | E1BNR0 | Apolipoprotein B OS=Bos taurus OX=9913 GN=APOB PE=1 SV=2 - [E1BNR0_BOVIN] | 0 | 0 | 0 | 38039264 | 11488055,5 | APOB | 1 | 24763659,8 | 24763659,75 | 7,393814828 |
| 121 | Q1JPF8 | 2-associated athanogene 4 (Fragment) OS=Bos taurus OX=9913 GN=BAG4 PE=2 SV=1 - [Q1JPF8_BO | 0 | 0 | 0 | 22043425,75 | 26518316,5 | BAG4 | 1 | 24280871,1 | 24280871,13 | 7,385264264 |
| 122 | Q2KJ42 | OU domain containing octamer binding OS=Bos taurus OX=9913 GN=NONO PE=2 SV=1 - [Q2KJ42_B | 0 | 0 | 0 | 27830085,5 | 20696267 | NONO | 1 | 24263176,3 | 24263176,25 | 7,384947653 |
| 123 | Q3T040 | S ribosomal protein S7, mitochondrial OS=Bos taurus OX=9913 GN=MRPS7 PE=1 SV=1 - [RT07_BOVI | 0 | 0 | 0 | 22060426,5 | 26390642,72 | MRPS7 | 1 | 24225534,6 | 24225534,61 | 7,38427337 |
| 124 | F1MB60 | 40S ribosomal protein S26 OS=Bos taurus OX=9913 GN=RPS26 PE=3 SV=1 - [F1MB60_BOVIN] | 0 | 0 | 0 | 44645382 | 3560193,063 | RPS26 | 1 | 24102787,5 | 24102787,53 | 7,382067272 |
| 125 | 0A140T8 | rotease homolog, mitochondrial OS=Bos taurus OX=9913 GN=LONP1 PE=3 SV=1 - [A0A140T842_BC | 0 | 0 | 0 | 12316086,5 | 33968246,88 | LONP1 | 1 | 23142166,7 | 23142166,69 | 7,364404017 |
| 126 | F1N0S9 | with sequence similarity 98 member A OS=Bos taurus OX=9913 GN=FAM98A PE=4 SV=2 - [F1N0S9_ | 0 | 0 | 0 | 34429693,72 | 11804878,41 | FAM98A | 1 | 23117286,1 | 23117286,06 | 7,363936847 |
| 127 | E1B7F5 | eat and HMG-box DNA binding protein 1 OS=Bos taurus OX=9913 GN=WDHD1 PE=4 SV=2 - [E1B7F5 | 0 | 0 | 0 | 21579148,67 | 23512206,24 | WDHD1 | 1 | 22545677,5 | 22545677,45 | 7,35306329 |
| 128 | Q0V8Q2 | ase activating protein 24 (Fragment) OS=Bos taurus OX=9913 GN=ARHGAP24 PE=2 SV=1 - [Q0V8Q2 | 0 | 0 | 0 | 24385710,36 | 19946731,63 | ARHGAP24 | 1 | 22166221 | 22166220,99 | 7,345691659 |
| 129 | Q148N0 | glutarate dehydrogenase, mitochondrial OS=Bos taurus OX=9913 GN=OGDH PE=2 SV=1 - [ODO1_B | 0 | 0 | 0 | 24702429,09 | 19543486,21 | OGDH | 1 | 22122957,7 | 22122957,65 | 7,344843188 |
| 130 | E1BDM8 | DEAD-box helicase 6 OS=Bos taurus OX=9913 GN=DDX6 PE=1 SV=1 - [E1BDM8_BOVIN] | 0 | 0 | 0 | 42369627,56 | 1395017,18 | DDX6 | 1 | 21882322,4 | 21882322,37 | 7,340093412 |
| 131 | Q58D08 | Pyrroline-5-carboxylate reductase 3 OS=Bos taurus OX=9913 GN=PYCR3 PE=2 SV=1 - [P5CR3_BOVIN | 0 | 0 | 0 | 22870718 | 19041428,56 | PYCR3 | 1 | 20956073,3 | 20956073,28 | 7,321309908 |
| 132 | G8JKY0 | 40S ribosomal protein S8 OS=Bos taurus OX=9913 GN=RPS8 PE=3 SV=1 - [G8JKY0_BOVIN] | 0 | 0 | 0 | 1644100,125 | 40012276,55 | RPS8 | 1 | 20828188,3 | 20828188,34 | 7,318651496 |
| 133 | F1N4M9 | ental retardation syndrome-related protein 1 OS=Bos taurus OX=9913 GN=FXR1 PE=4 SV=1 - [F1N4I | 0 | 0 | 0 | 32344197,69 | 8777155,625 | FXR1 | 1 | 20560676,7 | 20560676,66 | 7,313037403 |
| 134 | E1BGI2 | Centrosomal protein 170 OS=Bos taurus OX=9913 GN=CEP170 PE=4 SV=2 - [E1BGI2_BOVIN] | 0 | 0 | 0 | 25446065 | 15661584,71 | CEP170 | 1 | 20553824,9 | 20553824,85 | 7,312892651 |
| 135 | Q08E01 | Adducin 3 (Gamma) OS=Bos taurus OX=9913 GN=ADD3 PE=2 SV=1 - [Q08E01_BOVIN] | 0 | 0 | 0 | 29457088,5 | 10924995,34 | ADD3 | 1 | 20191041,9 | 20191041,92 | 7,30515873 |
| 136 | G1K134 | lear ribonucleoprotein-associated protein OS=Bos taurus OX=9913 GN=SNRPB PE=3 SV=1 - [G1K134 | 0 | 0 | 0 | 20391337,44 | 15301750,45 | SNRPB | 1 | 17846543,9 | 17846543,95 | 7,251554126 |
| 137 | Q3T0S7 | ome 14 open reading frame 166 ortholog OS=Bos taurus OX=9913 GN=RTRAF PE=2 SV=1 - [Q3T0S7 | 0 | 0 | 0 | 24257578,75 | 10935415,13 | RTRAF | 1 | 17596496,9 | 17596496,94 | 7,245426218 |
| 138 | A1A4J9 | sp40) homolog, subfamily A, member 3 OS=Bos taurus OX=9913 GN=DNAJA3 PE=2 SV=1 - [A1A4J9_ | 0 | 0 | 0 | 30158709,13 | 4950338,313 | DNAJA3 | 1 | 17554523,7 | 17554523,72 | 7,244389051 |
| 139 | A6H7H1 | coactivator of PPAR-gamma-like protein 1 OS=Bos taurus OX=9913 GN=FAM120A PE=2 SV=1 - [F12 | 0 | 0 | 0 | 15734566,05 | 17544431,46 | FAM120A | 1 | 16639498,8 | 16639498,76 | 7,22114024 |
| 140 | E1BQ37 | cing factor proline and glutamine rich OS=Bos taurus OX=9913 GN=SFPQ PE=1 SV=1 - [E1BQ37_BO\ | 0 | 0 | 0 | 26179435,8 | 7010466,625 | SFPQ | 1 | 16594951,2 | 16594951,21 | 7,21997598 |
| 141 | Q3SYV6 | Importin subunit alpha OS=Bos taurus OX=9913 GN=KPNA2 PE=1 SV=1 - [Q3SYV6_BOVIN] | 0 | 0 | 0 | 30644212,13 | 2487874,875 | KPNA2 | 1 | 16566043,5 | 16566043,5 | 7,219218797 |
| 142 | B0JYP5 | Protein argonaute-2 OS=Bos taurus OX=9913 GN=AGO2 PE=2 SV=1 - [B0JYP5_BOVIN] | 0 | 0 | 0 | 19694242,48 | 13024547,86 | AGO2 | 1 | 16359395,2 | 16359395,17 | 7,213767243 |
| 143 | Q5E966 | yotic translation initiation factor 3 subunit I OS=Bos taurus OX=9913 GN=EIF3I PE=2 SV=1 - [EIF3I_B | 0 | 0 | 0 | 18210260,89 | 14438844,28 | EIF3I | 1 | 16324552,6 | 16324552,59 | 7,212841287 |
| 144 | F1MCK2 | AHNAK nucleoprotein OS=Bos taurus OX=9913 GN=AHNAK PE=1 SV=2 - [F1MCK2_BOVIN] | 0 | 0 | 0 | 17568476,5 | 14860772,56 | AHNAK | 1 | 16214624,5 | 16214624,53 | 7,209906897 |
| 145 | Q1RMU4 | Erlin-2 OS=Bos taurus OX=9913 GN=ERLIN2 PE=2 SV=1 - [ERLN2_BOVIN] | 0 | 0 | 0 | 29970885,25 | 2377332,25 | ERLIN2 | 1 | 16174108,8 | 16174108,75 | 7,208820359 |
| 146 | 0A088Q0 | hock protein 90kDa alpha OS=Bos taurus x Bos grunniens OX=1167295 PE=2 SV=1 - [A0A088Q0F1_ | 0 | 0 | 0 | 23818960,15 | 8016960,375 | HSP90AA1 | 1 | 15917960,3 | 15917960,26 | 7,201887416 |
| 147 | Q5E947 | Peroxiredoxin-1 OS=Bos taurus OX=9913 GN=PRDX1 PE=2 SV=1 - [PRDX1_BOVIN] | 0 | 0 | 0 | 19236585,75 | 12227095,38 | PRDX1 | 1 | 15731840,6 | 15731840,56 | 7,196779536 |
| 148 | A4IFJ8 | ACTL6A protein OS=Bos taurus OX=9913 GN=ACTL6A PE=2 SV=1 - [A4IFJ8_BOVIN] | 0 | 0 | 0 | 23391426,66 | 7550402,5 | ACTL6A | 1 | 15470914,6 | 15470914,58 | 7,189515988 |
| 149 | Q2TA49 | asodilator-stimulated phosphoprotein OS=Bos taurus OX=9913 GN=VASP PE=2 SV=3 - [VASP_BOVIN | 0 | 0 | 0 | 4810699 | 25273589 | VASP | 1 | 15042144 | 15042144 | 7,177309742 |
| 150 | E1BNE5 | 311 family interacting protein 5 OS=Bos taurus OX=9913 GN=RAB11FIP5 PE=4 SV=2 - [E1BNE5_BO\ | 0 | 0 | 0 | 20555684,61 | 8985261,219 | RAB11FIP5 | 1 | 14770472,9 | 14770472,91 | 7,169394401 |
| 151 | Q3MIC0 | 60S ribosomal protein L37a OS=Bos taurus OX=9913 GN=RPL37A PE=3 SV=3 - [RL37A_BOVIN] | 0 | 0 | 0 | 21058411,5 | 7776261,703 | RPL37A | 1 | 14417336,6 | 14417336,6 | 7,158885038 |
| 152 | F1MZD9 | Nucleoporin 50 OS=Bos taurus OX=9913 GN=NUP50 PE=4 SV=1 - [F1MZD9_BOVIN] | 0 | 0 | 0 | 5619492,734 | 22653086,75 | NUP50 | 1 | 14136289,7 | 14136289,74 | 7,150335438 |
| 153 | G3X7R1 | e triphosphate triphosphohydrolase SAMHD1 OS=Bos taurus OX=9913 GN=SAMHD1 PE=4 SV=1 - [G | 0 | 0 | 0 | 8407368,563 | 19761902,5 | SAMHD1 | 1 | 14084635,5 | 14084635,53 | 7,148745613 |
| 154 | Q3SYW6 | otic translation initiation factor 3 subunit C OS=Bos taurus OX=9913 GN=EIF3C PE=2 SV=1 - [EIF3C_I | 0 | 0 | 0 | 12897815,11 | 15266658,43 | EIF3C | 1 | 14082236,8 | 14082236,77 | 7,148671642 |
| 155 | Q3T102 | otic translation initiation factor 3 subunit E OS=Bos taurus OX=9913 GN=EIF3E PE=2 SV=1 - [EIF3E_I | 0 | 0 | 0 | 10508030,88 | 16874415,44 | EIF3E | 1 | 13691223,2 | 13691223,16 | 7,136442249 |
| 156 | A4IF82 | MATR3 protein OS=Bos taurus OX=9913 GN=MATR3 PE=2 SV=1 - [A4IF82_BOVIN] | 0 | 0 | 0 | 10694572,31 | 16459216,2 | MATR3 | 1 | 13576894,3 | 13576894,26 | 7,132800436 |

| | | | | | | | | | | | | |
|---|---|---|---|---|---|---|---|---|---|---|---|---|
| 157 | A7MBB7 | NFKB2 protein OS=Bos taurus OX=9913 GN=NFKB2 PE=2 SV=1 - [A7MBB7_BOVIN] | 0 | 0 | 0 | 17117034,75 | 10017512,84 | NFKB2 | 1 | 13567273,8 | 13567273,8 | 7,132492589 |
| 158 | Q8WN55 | olypyrimidine tract-binding protein 1 OS=Bos taurus OX=9913 GN=PTBP1 PE=2 SV=1 - [PTBP1_BOVI | 0 | 0 | 0 | 10090301,75 | 16958120 | PTBP1 | 1 | 13524210,9 | 13524210,88 | 7,131111934 |
| 159 | Q9BGI2 | Peroxiredoxin-4 OS=Bos taurus OX=9913 GN=PRDX4 PE=2 SV=1 - [PRDX4_BOVIN] | 0 | 0 | 0 | 14599210,75 | 11930718 | PRDX4 | 1 | 13264964,4 | 13264964,38 | 7,122706088 |
| 160 | F1MRZ0 | Angiomotin OS=Bos taurus OX=9913 GN=AMOT PE=4 SV=2 - [F1MRZ0_BOVIN] | 0 | 0 | 0 | 14937752,75 | 10268887,14 | AMOT | 1 | 12603319,9 | 12603319,95 | 7,100484961 |
| 161 | F1N206 | Dihydrolipoyl dehydrogenase OS=Bos taurus OX=9913 GN=DLD PE=1 SV=2 - [F1N206_BOVIN] | 0 | 0 | 0 | 2213048,422 | 22748805,88 | DLD | 1 | 12480927,1 | 12480927,15 | 7,096246848 |
| 162 | E1BEK3 | uclear organelle associated protein 2 OS=Bos taurus OX=9913 GN=GEMIN2 PE=4 SV=1 - [E1BEK3_E | 0 | 0 | 0 | 5574329 | 16991781,56 | GEMIN2 | 1 | 11283055,3 | 11283055,28 | 7,052426716 |
| 163 | A5D7P5 | nma-tubulin complex component OS=Bos taurus OX=9913 GN=TUBGCP2 PE=2 SV=1 - [A5D7P5_BO | 0 | 0 | 0 | 13550875,46 | 8987545 | TUBGCP2 | 1 | 11269210,2 | 11269210,23 | 7,051893481 |
| 164 | A6QLN8 | SCYL2 protein OS=Bos taurus OX=9913 GN=SCYL2 PE=2 SV=1 - [A6QLN8_BOVIN] | 0 | 0 | 0 | 6237056,844 | 15784443,22 | SCYL2 | 1 | 11010750 | 11010750,03 | 7,041816903 |
| 165 | E1BNG7 | nma-tubulin complex component OS=Bos taurus OX=9913 GN=TUBGCP3 PE=3 SV=1 - [E1BNG7_BO | 0 | 0 | 0 | 15617109,26 | 6239693,219 | TUBGCP3 | 1 | 10928401,2 | 10928401,24 | 7,038556632 |
| 166 | Q2KIR1 | 1 small nuclear ribonucleoprotein A OS=Bos taurus OX=9913 GN=SNRPA PE=2 SV=1 - [SNRPA_BOVI | 0 | 0 | 0 | 17498072,44 | 4063535,25 | SNRPA | 1 | 10780803,8 | 10780803,84 | 7,032651144 |
| 167 | E1BEQ5 | Angiomotin-like protein 1 OS=Bos taurus OX=9913 GN=AMOTL1 PE=3 SV=1 - [AMOL1_BOVIN] | 0 | 0 | 0 | 11950088,35 | 9161458,625 | AMOTL1 | 1 | 10555773,5 | 10555773,49 | 7,023490062 |
| 168 | A1L545 | Gemin4 (Fragment) OS=Bos taurus OX=9913 GN=GEMIN4 PE=2 SV=1 - [A1L545_BOVIN] | 0 | 0 | 0 | 8388074,375 | 12381267,92 | GEMIN4 | 1 | 10384671,1 | 10384671,15 | 7,016392748 |
| 169 | E1BFI4 | growth factor 2 mRNA binding protein 2 OS=Bos taurus OX=9913 GN=IGF2BP2 PE=4 SV=2 - [E1BFI | 0 | 0 | 0 | 11951398,38 | 8321546,938 | IGF2BP2 | 1 | 10136472,7 | 10136472,66 | 7,005886853 |
| 170 | G3X6H0 | Centrosomal protein 170B OS=Bos taurus OX=9913 GN=CEP170B PE=4 SV=1 - [G3X6H0_BOVIN] | 0 | 0 | 0 | 11709250,33 | 7425810,172 | CEP170B | 1 | 9567530,25 | 9567530,25 | 6,980799844 |
| 171 | Q769I5 | Hepatocyte growth factor receptor OS=Bos taurus OX=9913 GN=MET PE=2 SV=1 - [MET_BOVIN] | 0 | 0 | 0 | 16993906,63 | 1787210,781 | MET | 1 | 9390558,7 | 9390558,703 | 6,972691432 |
| 172 | Q1JQD2 | te-rich WD repeat-containing protein 1 OS=Bos taurus OX=9913 GN=GRWD1 PE=2 SV=1 - [GRWD1 | 0 | 0 | 0 | 15651130,25 | 2624229,328 | GRWD1 | 1 | 9137679,79 | 9137679,789 | 6,960835935 |
| 173 | F1MSE4 | Gephyrin OS=Bos taurus OX=9913 GN=GPHN PE=4 SV=2 - [F1MSE4_BOVIN] | 0 | 0 | 0 | 12593524,78 | 4581652,25 | GPHN | 1 | 8587588,52 | 8587588,516 | 6,933871227 |
| 174 | A6QNM0 | Septin 7 OS=Bos taurus OX=9913 GN=SEPT7 PE=2 SV=1 - [A6QNM0_BOVIN] | 0 | 0 | 0 | 3874664,313 | 13284385,19 | SEPT7 | 1 | 8579524,75 | 8579524,75 | 6,933463231 |
| 175 | Q5E954 | aJ homolog subfamily A member 1 OS=Bos taurus OX=9913 GN=DNAJA1 PE=2 SV=2 - [DNJA1_BOVI | 0 | 0 | 0 | 8184309,352 | 8400365 | DNAJA1 | 1 | 8292337,18 | 8292337,176 | 6,918676953 |
| 176 | A6QLV1 | d1 and UNC84 domain containing 2 OS=Bos taurus OX=9913 GN=SUN2 PE=2 SV=1 - [A6QLV1_BOVI | 0 | 0 | 0 | 12204367,38 | 4376061,9 | SUN2 | 1 | 8290214,64 | 8290214,638 | 6,918565775 |
| 177 | F1MX25 | inucleotide repeat containing 6A OS=Bos taurus OX=9913 GN=TNRC6A PE=4 SV=1 - [F1MX25_BOVI | 0 | 0 | 0 | 12896977,35 | 3079126,453 | TNRC6A | 1 | 7988051,9 | 7988051,902 | 6,902440878 |
| 178 | Q1JPA2 | nslation elongation factor 1 gamma (Fragment) OS=Bos taurus OX=9913 GN=EEF1G PE=2 SV=1 - [Q | 0 | 0 | 0 | 12486790,81 | 3251411 | EEF1G | 1 | 7869100,91 | 7869100,906 | 6,895925114 |
| 179 | E1BIQ0 | bonucleotide nucleotidyltransferase 1 OS=Bos taurus OX=9913 GN=PNPT1 PE=4 SV=2 - [E1BIQ0_B( | 0 | 0 | 0 | 5089803,875 | 10041502,52 | PNPT1 | 1 | 7565653,2 | 7565653,195 | 6,87884643 |
| 180 | F1MK44 | Integrin alpha-5 OS=Bos taurus OX=9913 GN=ITGA5 PE=3 SV=2 - [F1MK44_BOVIN] | 0 | 0 | 0 | 10382866,25 | 4613218,594 | ITGA5 | 1 | 7498042,42 | 7498042,422 | 6,874947893 |
| 181 | Q58DT4 | :-5-carboxylate reductase 1, mitochondrial OS=Bos taurus OX=9913 GN=PYCR1 PE=2 SV=1 - [P5CR1 | 0 | 0 | 0 | 4429208,688 | 10412831,88 | PYCR1 | 1 | 7421020,28 | 7421020,281 | 6,870463618 |
| 182 | A7MBG8 | RuvB-like helicase OS=Bos taurus OX=9913 GN=RUVBL1 PE=1 SV=1 - [A7MBG8_BOVIN] | 0 | 0 | 0 | 7635109,5 | 6221790,602 | RUVBL1 | 1 | 6928450,05 | 6928450,051 | 6,84063609 |
| 183 | E1BLT0 | Uncharacterized protein OS=Bos taurus OX=9913 GN=GOLGA2 PE=4 SV=2 - [E1BLT0_BOVIN] | 0 | 0 | 0 | 4401249,109 | 9177094,703 | GOLGA2 | 1 | 6789171,91 | 6789171,906 | 6,831816805 |
| 184 | E1BJG5 | Tripartite motif containing 28 OS=Bos taurus OX=9913 GN=TRIM28 PE=4 SV=2 - [E1BJG5_BOVIN] | 0 | 0 | 0 | 9416128,375 | 3806240,375 | TRIM28 | 1 | 6611184,38 | 6611184,375 | 6,820279269 |
| 185 | F1N1S1 | ibosomal protein S27, mitochondrial OS=Bos taurus OX=9913 GN=MRPS27 PE=4 SV=1 - [F1N1S1_B( | 0 | 0 | 0 | 2375571,688 | 10589136,63 | MRPS27 | 1 | 6482354,16 | 6482354,156 | 6,811732755 |
| 186 | A2VE34 | PABPN1 protein OS=Bos taurus OX=9913 GN=PABPN1 PE=2 SV=1 - [A2VE34_BOVIN] | 0 | 0 | 0 | 7852942,375 | 4530189 | PABPN1 | 1 | 6191565,69 | 6191565,688 | 6,791800485 |
| 187 | E1BG23 | Tripartite motif containing 47 OS=Bos taurus OX=9913 GN=TRIM47 PE=4 SV=2 - [E1BG23_BOVIN] | 0 | 0 | 0 | 8823930,625 | 2986474,813 | TRIM47 | 1 | 5905202,72 | 5905202,719 | 6,771234811 |
| 188 | A5D7D9 | DEAH-box helicase 15 OS=Bos taurus OX=9913 GN=DHX15 PE=1 SV=1 - [A5D7D9_BOVIN] | 0 | 0 | 0 | 8691191,5 | 3054883,625 | DHX15 | 1 | 5873037,56 | 5873037,563 | 6,768862778 |
| 189 | A7E3W2 | Galectin-3-binding protein OS=Bos taurus OX=9913 GN=LGALS3BP PE=1 SV=1 - [LG3BP_BOVIN] | 0 | 0 | 0 | 2527031,586 | 9195251,422 | LGALS3BP | 1 | 5861141,5 | 5861141,504 | 6,767982207 |
| 190 | A6QQ70 | Aicrotubule-associated protein 1S OS=Bos taurus OX=9913 GN=MAP1S PE=2 SV=1 - [MAP1S_BOVIN | 0 | 0 | 0 | 3096446,25 | 7813659,75 | MAP1S | 1 | 5455053 | 5455053 | 6,736798974 |
| 191 | E1BBK8 | Uncharacterized protein OS=Bos taurus OX=9913 GN=PPFIA1 PE=4 SV=2 - [E1BBK8_BOVIN] | 0 | 0 | 0 | 5421264,188 | 5436805,938 | PPFIA1 | 1 | 5429035,06 | 5429035,063 | 6,734722646 |
| 192 | Q3SZI4 | 14-3-3 protein theta OS=Bos taurus OX=9913 GN=YWHAQ PE=2 SV=1 - [1433T_BOVIN] | 0 | 0 | 0 | 4425293,25 | 6203959,688 | YWHAQ | 1 | 5314626,47 | 5314626,469 | 6,725472746 |
| 193 | E1B970 | Golgin A3 OS=Bos taurus OX=9913 GN=GOLGA3 PE=4 SV=1 - [E1B970_BOVIN] | 0 | 0 | 0 | 6082440,391 | 3996696,5 | GOLGA3 | 1 | 5039568,45 | 5039568,445 | 6,702393348 |
| 194 | E1BQ03 | Nucleus accumbens associated 1 OS=Bos taurus OX=9913 GN=NACC1 PE=4 SV=1 - [E1BQ03_BOVIN] | 0 | 0 | 0 | 6127413,469 | 3598887,25 | NACC1 | 1 | 4863150,36 | 4863150,359 | 6,686917697 |
| 195 | Q17QN2 | Target of EGR1 protein 1 OS=Bos taurus OX=9913 GN=TOE1 PE=2 SV=1 - [TOE1_BOVIN] | 0 | 0 | 0 | 7184137,016 | 2490147,125 | TOE1 | 1 | 4837142,07 | 4837142,07 | 6,684588843 |
| 196 | Q58DQ5- | form 3 of 28S ribosomal protein S9, mitochondrial OS=Bos taurus OX=9913 GN=MRPS9 - [RT09_BOV | 0 | 0 | 0 | 4740711,875 | 4731404,438 | MRPS9 | 1 | 4736058,16 | 4736058,156 | 6,675417027 |
| 197 | P63103 | 14-3-3 protein zeta/delta OS=Bos taurus OX=9913 GN=YWHAZ PE=1 SV=1 - [1433Z_BOVIN] | 0 | 0 | 0 | 5558586,25 | 2142611,438 | YWHAZ | 1 | 3850598,84 | 3850598,844 | 6,585528276 |
| 198 | A7MB01 | SART3 protein OS=Bos taurus OX=9913 GN=SART3 PE=2 SV=1 - [A7MB01_BOVIN] | 0 | 0 | 0 | 2293788,313 | 4634186,348 | SART3 | 1 | 3463987,33 | 3463987,33 | 6,539576295 |
| 199 | E1BP72 | C42 binding protein kinase beta OS=Bos taurus OX=9913 GN=CDC42BPB PE=3 SV=2 - [E1BP72_BOV | 0 | 0 | 0 | 3529094,875 | 2293387,703 | CDC42BPB | 1 | 2911241,29 | 2911241,289 | 6,464078202 |
| 200 | F1N1G7 | Kinesin-like protein OS=Bos taurus OX=9913 GN=KIF5B PE=1 SV=2 - [F1N1G7_BOVIN] | 0 | 0 | 0 | 4323816,875 | 1352252,875 | KIF5B | 1 | 2838034,88 | 2838034,875 | 6,453017728 |
| 201 | Q3ZCK1 | otic translation initiation factor 3 subunit L OS=Bos taurus OX=9913 GN=EIF3L PE=2 SV=1 - [EIF3L_E | 0 | 0 | 0 | 3342069,073 | 1918102 | EIF3L | 1 | 2630085,54 | 2630085,536 | 6,419969873 |
| 202 | F1MX04 | c translation initiation factor 4 gamma 1 OS=Bos taurus OX=9913 GN=EIF4G1 PE=1 SV=2 - [F1MX04 | 0 | 0 | 0 | 3816766,875 | 1003856,906 | EIF4G1 | 1 | 2410311,89 | 2410311,891 | 6,382073243 |
| 203 | Q95KE5 | ribosomal protein L43, mitochondrial OS=Bos taurus OX=9913 GN=MRPL43 PE=2 SV=2 - [RM43_BO | 0 | 0 | 0 | 2002083,469 | 2699837 | MRPL43 | 1 | 2350960,23 | 2350960,234 | 6,371245283 |
| 204 | F1MLI8 | Podocalyxin like OS=Bos taurus OX=9913 GN=PODXL PE=4 SV=2 - [F1MLI8_BOVIN] | 0 | 0 | 0 | 878634,3281 | 2680341,688 | PODXL | 1 | 1779488,01 | 1779488,008 | 6,250295066 |
| 205 | Q27966-3 | Isoform 3 of Unconventional myosin-Ic OS=Bos taurus OX=9913 GN=MYO1C - [MYO1C_BOVIN] | 1184507,438 | 0 | 0 | 3582143005 | 27152420,42 | MYO1C | 394835,81 | 1804647713 | 4570,628235 | 3,659975898 |
| 206 | A1L5B0 | regulatory light polypeptide 9 (Fragment) OS=Bos taurus OX=9913 GN=MYL9 PE=2 SV=1 - [A1L5B0_ | 4386960,75 | 0 | 0 | 8596551877 | 32350351,5 | MYL9 | 1462320,3 | 4314451114 | 2950,414668 | 3,469883058 |
| 207 | A4IF97 | Ayosin regulatory light chain 12B OS=Bos taurus OX=9913 GN=MYL12B PE=2 SV=1 - [ML12B_BOVIN | 10034338,33 | 0 | 0 | 14557775994 | 657173033,6 | MYL12B | 3344779,4 | 7607474514 | 2274,432333 | 3,356873021 |
| 208 | P60661-2 | form Smooth muscle of Myosin light polypeptide 6 OS=Bos taurus OX=9913 GN=MYL6 - [MYL6_BOV | 7341425,607 | 13542839,03 | 0 | 13378911295 | 138188667,2 | MYL6 | 6961421,5 | 6758549981 | 970,8577388 | 2,987155597 |
| 209 | P60661 | Myosin light polypeptide 6 OS=Bos taurus OX=9913 GN=MYL6 PE=2 SV=2 - [MYL6_BOVIN] | 7341425,607 | 13542839,03 | 0 | 13378911295 | 67713106,02 | MYL6 | 6961421,5 | 6723312200 | 965,7958731 | 2,984885345 |
| 210 | E1BNE7 | Caveolae associated protein 1 OS=Bos taurus OX=9913 GN=CAVIN1 PE=1 SV=1 - [E1BNE7_BOVIN] | 5894706,469 | 0 | 0 | 2200492298 | 44470787,78 | CAVIN1 | 1964902,2 | 1122481543 | 571,2658717 | 2,756838279 |
| 211 | F1MQ37 | Myosin heavy chain 9 OS=Bos taurus OX=9913 GN=MYH9 PE=1 SV=2 - [F1MQ37_BOVIN] | 60201715,69 | 4101663,109 | 6815719,375 | 23196978930 | 256242696,8 | MYH9 | 23706366 | 1,1727E+10 | 494,6608344 | 2,694307526 |
| 212 | F1MM57 | Myosin heavy chain 14 OS=Bos taurus OX=9913 GN=MYH14 PE=1 SV=2 - [F1MM57_BOVIN] | 31814905,5 | 0 | 0 | 10109789782 | 18138518,34 | MYH14 | 10604969 | 5063964150 | 477,5086461 | 2,67898124 |
| 213 | P48616 | Vimentin OS=Bos taurus OX=9913 GN=VIM PE=1 SV=3 - [VIME_BOVIN] | 17732996,18 | 30572050,47 | 0 | 12978723068 | 1058743913 | VIM | 16101682 | 7018733490 | 435,9006342 | 2,639387501 |
| 214 | O02717 | Non-muscle myosin heavy chain (Fragment) OS=Bos taurus OX=9913 PE=2 SV=2 - [O02717_BOVIN] | 54574194,31 | 0 | 7721628,313 | 14557834532 | 89733942,56 | NMMHC | 20765274 | 7323784237 | 352,693837 | 2,547397871 |
| 215 | P60712 | Actin, cytoplasmic 1 OS=Bos taurus OX=9913 GN=ACTB PE=1 SV=1 - [ACTB_BOVIN] | 216428994,6 | 20689487,5 | 0 | 50898753883 | 1438088556 | ACTB | 79039494 | 2,6168E+10 | 331,080323 | 2,51993337 |
| 216 | F1MRD0 | Actin, cytoplasmic 1 OS=Bos taurus OX=9913 GN=ACTB PE=3 SV=2 - [F1MRD0_BOVIN] | 216428994,6 | 20689487,5 | 0 | 42233747621 | 1438088556 | ACTB | 79039494 | 2,1836E+10 | 276,2659143 | 2,441327305 |
| 217 | F1N3A1 | Thrombospondin-1 OS=Bos taurus OX=9913 GN=THBS1 PE=4 SV=1 - [F1N3A1_BOVIN] | 968187,5313 | 0 | 0 | 109786689,9 | 34909483,81 | THBS1 | 322729,18 | 72348086,9 | 224,1758477 | 2,350588821 |
| 218 | Q27991 | Myosin-10 OS=Bos taurus OX=9913 GN=MYH10 PE=2 SV=2 - [MYH10_BOVIN] | 56625179,96 | 1986835,5 | 0 | 7693202318 | 62874653,5 | MYH10 | 19537338 | 3878038486 | 198,4936939 | 2,297746714 |
| 219 | )A0F6QNI | Complement component 3 OS=Bos taurus OX=9913 GN=C3 PE=2 SV=1 - [A0A0F6QNP7_BOVIN] | 9582357,266 | 0 | 0 | 476223491 | 538893442,9 | C3 | 3194119,1 | 507558467 | 158,9040524 | 2,201134973 |
| 220 | Q7SIH1 | Alpha-2-macroglobulin OS=Bos taurus OX=9913 GN=A2M PE=1 SV=2 - [A2MG_BOVIN] | 11365333,78 | 3012582,875 | 0 | 702455107,6 | 790788609,9 | A2M | 4792638,9 | 746621859 | 155,7851273 | 2,192525993 |
| 221 | A4IFM8 | Actin, alpha 1, skeletal muscle OS=Bos taurus OX=9913 GN=ACTA1 PE=2 SV=1 - [A4IFM8_BOVIN] | 242576162,8 | 20689487,5 | 0 | 26550230012 | 681182520,9 | ACTA1 | 87755217 | 1,3616E+10 | 155,1555197 | 2,19076723 |
| 222 | P11179 | ferase component of 2-oxoglutarate dehydrogenase complex, mitochondrial OS=Bos taurus OX=99: | 0 | 2748994,266 | 0 | 114924907,1 | 157214007,3 | DLST | 916331,42 | 136069457 | 148,4937152 | 2,171708073 |
| 223 | Q3T087 | 60S ribosomal protein L11 OS=Bos taurus OX=9913 GN=RPL11 PE=2 SV=3 - [RL11_BOVIN] | 10168574,75 | 0 | 0 | 883062340,5 | 34047638,5 | RPL11 | 3389524,9 | 458554990 | 135,2859179 | 2,131252593 |
| 224 | Q28194 | Thrombospondin-1 (Fragment) OS=Bos taurus OX=9913 PE=2 SV=1 - [Q28194_BOVIN] | 968187,5313 | 0 | 0 | 64597779 | 17486781,25 | THBS1 | 322729,18 | 41042280,1 | 127,1725119 | 2,10439325 |
| 225 | P61286 | Polyadenylate-binding protein 1 OS=Bos taurus OX=9913 GN=PABPC1 PE=2 SV=1 - [PABP1_BOVIN] | 11659910,1 | 0 | 0 | 642506450,5 | 247961754 | PABPC1 | 3886636,7 | 445234102 | 114,5551119 | 2,059014474 |
| 226 | F1MJH1 | Gelsolin OS=Bos taurus OX=9913 GN=GSN PE=4 SV=1 - [F1MJH1_BOVIN] | 12086506,5 | 0 | 0 | 746119472,3 | 174787785,8 | GSN | 4028835,5 | 460453629 | 114,2895085 | 2,058006365 |
| 227 | Q3T0U2 | 60S ribosomal protein L14 OS=Bos taurus OX=9913 GN=RPL14 PE=2 SV=3 - [RL14_BOVIN] | 0 | 4570460 | 0 | 281233892,6 | 19706536 | RPL14 | 1523486,7 | 150470214 | 98,7670044 | 1,994611882 |
| 228 | A5D7R6 | ITIH2 protein OS=Bos taurus OX=9913 GN=ITIH2 PE=2 SV=1 - [A5D7R6_BOVIN] | 4037421,5 | 0 | 0 | 225777609,2 | 11115438,88 | ITIH2 | 1345807,2 | 118446524 | 88,01151231 | 1,944539484 |
| 229 | P63243 | :eptor of activated protein C kinase 1 OS=Bos taurus OX=9913 GN=RACK1 PE=2 SV=3 - [RACK1_BOV | 7575398,027 | 0 | 0 | 407666437,5 | 34883619,13 | RACK1 | 2525132,7 | 221275028 | 87,62907012 | 1,942648203 |
| 230 | P00974 | Pancreatic trypsin inhibitor OS=Bos taurus OX=9913 PE=1 SV=2 - [BPT1_BOVIN] | 0 | 0 | 37649122,5 | 1978943599 | 39691626 | PSTI | 12549708 | 1009317612 | 80,42558858 | 1,905394248 |
| 231 | Q3T0X6 | 40S ribosomal protein S16 OS=Bos taurus OX=9913 GN=RPS16 PE=2 SV=3 - [RS16_BOVIN] | 6253575,75 | 0 | 0 | 217885540 | 29552092,19 | RPS16 | 2084525,3 | 123718816 | 59,35107579 | 1,773428595 |
| 232 | F1MT45 | Histone H2A OS=Bos taurus OX=9913 GN=H2AFZ PE=3 SV=2 - [F1MT45_BOVIN] | 4534252,828 | 0 | 0 | 136121315 | 5144049,438 | H2AFZ | 1511417,6 | 70632682,2 | 46,73273739 | 1,669621221 |
| 233 | Q4VQD6 | Actin (Fragment) OS=Cryptosporidium sp. ex Bos taurus OX=310049 PE=3 SV=1 - [Q4VQD6_9CRYT] | 120310954,5 | 0 | 0 | 3479561322 | 150332566,2 | ACT | 40103652 | 1814946944 | 45,25640125 | 1,655680016 |
| 234 | Q3T169 | 40S ribosomal protein S3 OS=Bos taurus OX=9913 GN=RPS3 PE=2 SV=1 - [RS3_BOVIN] | 8535379,75 | 8992976,938 | 5060503,953 | 443603839,7 | 200373988,4 | RPS3 | 7529620,2 | 321988914 | 42,76296877 | 1,631067848 |
| 235 | P09867 | geneous nuclear ribonucleoprotein A1 OS=Bos taurus OX=9913 GN=HNRNPA1 PE=1 SV=2 - [ROA1_ | 8950052,078 | 0 | 0 | 121004247,5 | 109672578,5 | HNRNPA1 | 2983350,7 | 115338413 | 38,6606956 | 1,587269664 |
| 236 | Q862B7 | Similar to ribosomal protein S14 (Fragment) OS=Bos taurus OX=9913 PE=2 SV=1 - [Q862B7_BOVIN] | 4963636,5 | 0 | 11375034,75 | 304357132,4 | 104684950,1 | RPL14 | 5446223,8 | 204521041 | 37,55281653 | 1,574642515 |
| 237 | P42899 | 60S acidic ribosomal protein P2 OS=Bos taurus OX=9913 GN=RPLP2 PE=3 SV=1 - [RLA2_BOVIN] | 297320,3125 | 2451246,031 | 17569122,13 | 467909737,5 | 35540378,79 | RPLP2 | 6772562,8 | 251725058 | 37,16836074 | 1,570173408 |
| 238 | Q862S8 | Similar to ribosomal protein S18 (Fragment) OS=Bos taurus OX=9913 PE=2 SV=1 - [Q862S8_BOVIN] | 0 | 0 | 8944804,75 | 181415429,8 | 39344590,94 | RPS18 | 2981601,6 | 110380010 | 37,02037555 | 1,56844082 |
| 239 | E1BH06 | Uncharacterized protein OS=Bos taurus OX=9913 GN=C4A PE=1 SV=2 - [E1BH06_BOVIN] | 4555315,125 | 0 | 0 | 67475660 | 35590448,05 | C4A | 1518438,4 | 51533054 | 33,9381926 | 1,53068871 |
| 240 | A7MBI5 | DPYSL3 protein OS=Bos taurus OX=9913 GN=DPYSL3 PE=1 SV=1 - [A7MBI5_BOVIN] | 0 | 1642842,711 | 7065867,563 | 119179414,4 | 69431978,23 | DPYSL3 | 2902903,4 | 94305696,3 | 32,48668059 | 1,511705338 |
| 241 | P32007 | ADP/ATP translocase 3 OS=Bos taurus OX=9913 GN=SLC25A6 PE=1 SV=3 - [ADT3_BOVIN] | 15409452,16 | 0 | 0 | 270198657,9 | 57955327,89 | SLC25A6 | 5136484,1 | 164076993 | 31,94344443 | 1,504381744 |
| 242 | P19120 | Heat shock cognate 71 kDa protein OS=Bos taurus OX=9913 GN=HSPA8 PE=1 SV=2 - [HSP7C_BOVIN | 20630150,78 | 8923548,125 | 16364886,64 | 592181739,1 | 354444661,9 | HSPA8 | 15306195 | 473313200 | 30,92298215 | 1,49028137 |
| 243 | Q8SQH5 | ADP/ATP translocase 2 OS=Bos taurus OX=9913 GN=SLC25A5 PE=2 SV=3 - [ADT2_BOVIN] | 19705901,15 | 0 | 0 | 326705654,3 | 57955327,89 | SLC25A5 | 6568633,7 | 192330491 | 29,28013639 | 1,466573095 |
| 244 | A8E651 | EWS RNA binding protein 1 OS=Bos taurus OX=9913 GN=EWSR1 PE=2 SV=1 - [A8E651_BOVIN] | 127320961,5 | 10406270,7 | 42506541,9 | 2571883455 | 774709963 | EWSR1 | 60077925 | 1673296709 | 27,8521057 | 1,444858035 |
| 245 | A7E355 | SYNCRIP protein (Fragment) OS=Bos taurus OX=9913 GN=SYNCRIP PE=2 SV=1 - [A7E355_BOVIN] | 0 | 0 | 17149812,25 | 250827063 | 59532666,42 | SYNCRIP | 5716604,1 | 155179865 | 27,14546301 | 1,433697254 |
| 246 | Q2KJD0 | Tubulin beta-5 chain OS=Bos taurus OX=9913 GN=TUBB5 PE=2 SV=1 - [TBB5_BOVIN] | 22199391,44 | 0 | 0 | 147686932,1 | 210989220,8 | TUBB5 | 7399797,1 | 179338076 | 24,23553955 | 1,384452693 |
| 247 | Q3T0D0 | geneous nuclear ribonucleoprotein K OS=Bos taurus OX=9913 GN=HNRNPK PE=2 SV=1 - [HNRPK_E | 10811448,13 | 0 | 0 | 105012347,8 | 68809073,95 | HNRNPK | 3603816 | 86910710,9 | 24,11630057 | 1,382310688 |
| 248 | Q32PB9 | 60S ribosomal protein L38 OS=Bos taurus OX=9913 GN=RPL38 PE=3 SV=4 - [RL38_BOVIN] | 3762463,5 | 0 | 0 | 35789661 | 24058167,25 | RPL38 | 1254154,5 | 29923914,1 | 23,85983077 | 1,377667359 |
| 249 | E1BPR4 | molog B, COPII coat complex component OS=Bos taurus OX=9913 GN=SEC24B PE=4 SV=1 - [E1BPR4 | 8101564 | 0 | 0 | 79523618,13 | 48907037,48 | SEC24B | 2700521,3 | 64215327,8 | 23,77886337 | 1,376191091 |
| 250 | F2Z4C1 | Tubulin alpha chain OS=Bos taurus OX=9913 GN=TUBA1A PE=3 SV=1 - [F2Z4C1_BOVIN] | 23495108,25 | 0 | 0 | 205713593,1 | 140429900 | TUBA1A | 7831702,8 | 173071747 | 22,09886561 | 1,344369981 |
| 251 | P81947 | Tubulin alpha-1B chain OS=Bos taurus OX=9913 PE=1 SV=2 - [TBA1B_BOVIN] | 23495108,25 | 0 | 0 | 205713593,1 | 140429900 | TUBA1B | 7831702,8 | 173071747 | 22,09886561 | 1,344369981 |
| 252 | E1B9K1 | Polyubiquitin-C OS=Bos taurus OX=9913 GN=UBC PE=4 SV=2 - [E1B9K1_BOVIN] | 28036531 | 0 | 0 | 364310184,5 | 41854107,03 | UBC | 9345510,3 | 203082146 | 21,73045008 | 1,337068721 |
| 253 | Q3ZBX9 | Histone H2A.J OS=Bos taurus OX=9913 GN=H2AFJ PE=2 SV=1 - [H2AJ_BOVIN] | 9879948,734 | 0 | 0 | 136121315 | 5144049,438 | H2AFJ | 3293316,2 | 70632682,2 | 21,447282 | 1,331372262 |
| 254 | Q3ZBH3 | CD151 antigen OS=Bos taurus OX=9913 GN=CD151 PE=2 SV=1 - [CD151_BOVIN] | 18285378,13 | 0 | 0 | 169784229 | 74636893,25 | CD151 | 6095126 | 122210561 | 20,05053879 | 1,302126047 |
| 255 | E1BEG2 | eneous nuclear ribonucleoprotein A3 OS=Bos taurus OX=9913 GN=HNRNPA3 PE=1 SV=2 - [E1BEG2_ | 33832059,09 | 0 | 0 | 89535057,56 | 349174283,4 | HNRNPA3 | 11277353 | 219354671 | 19,45090039 | 1,28893971 |
| 256 | F1MZ92 | ase-sensitive element-binding protein 1 OS=Bos taurus OX=9913 GN=YBX1 PE=4 SV=2 - [F1MZ92_B | 26195468,61 | 0 | 54100940,53 | 671664679,8 | 362809566,9 | YBX1 | 26765470 | 517237123 | 19,32479156 | 1,286114818 |
| 257 | F1N6K7 | Protein transport protein sec16 OS=Bos taurus OX=9913 GN=SEC16A PE=3 SV=2 - [F1N6K7_BOVIN] | 10235103,28 | 0 | 8997560,5 | 126440371,1 | 109026773,6 | SEC16A | 6410887,9 | 117733572 | 18,36462806 | 1,263982137 |
| 258 | Q3ZC44 | :neous nuclear ribonucleoprotein A/B OS=Bos taurus OX=9913 GN=HNRNPAB PE=1 SV=1 - [Q3ZC44 | 30972526,28 | 0 | 0 | 207672614,8 | 165947029,9 | HNRNPAB | 10324175 | 186809822 | 18,09440605 | 1,257544332 |
| 259 | E1BJB1 | Tubulin beta chain OS=Bos taurus OX=9913 GN=TUBB2A PE=1 SV=1 - [E1BJB1_BOVIN] | 28686769,88 | 0 | 0 | 130775923,3 | 210989220,8 | TUBB2A | 9562256,6 | 170882572 | 17,87052772 | 1,252137377 |
| 260 | A5D7Q4 | CSDA protein OS=Bos taurus OX=9913 GN=YBX3 PE=1 SV=1 - [A5D7Q4_BOVIN] | 30003877,98 | 0 | 0 | 284965667,9 | 56687237,42 | YBX3 | 10001293 | 170826453 | 17,08043734 | 1,232498987 |
| 261 | Q3ZCH0 | Stress-70 protein, mitochondrial OS=Bos taurus OX=9913 GN=HSPA9 PE=2 SV=1 - [GRP75_BOVIN] | 28553294,39 | 10997162,68 | 20405363,07 | 460409037,7 | 214454946,9 | HSPA9 | 19985273 | 337431992 | 16,88403185 | 1,227476163 |
| 262 | Q3MHM5 | Tubulin beta-4B chain OS=Bos taurus OX=9913 GN=TUBB4B PE=2 SV=1 - [TBB4B_BOVIN] | 28686769,88 | 0 | 0 | 133680207,9 | 177610056,4 | TUBB4B | 9562256,6 | 155645132 | 16,27702939 | 1,211575148 |
| 263 | A1A4R1 | Histone H2A type 2-C OS=Bos taurus OX=9913 GN=HIST2H2AC PE=2 SV=1 - [H2A2C_BOVIN] | 15256373,06 | 0 | 0 | 136121315 | 5144049,438 | HIST2H2AC | 5085457,7 | 70632682,2 | 13,88914952 | 1,142675653 |
| 264 | G5E631 | Uncharacterized protein OS=Bos taurus OX=9913 GN=DDX3X PE=1 SV=1 - [G5E631_BOVIN] | 174143462,3 | 5158042 | 0 | 881104468,2 | 758908529,3 | DDX3X | 59767168 | 820006499 | 13,72001594 | 1,137354616 |
| 265 | A5D9H5 | geneous nuclear ribonucleoprotein D OS=Bos taurus OX=9913 GN=HNRNPD PE=1 SV=1 - [A5D9H5_ | 43719434,52 | 0 | 0 | 182654283,4 | 210099099,7 | HNRNPD | 14573145 | 196376692 | 13,47524462 | 1,129536658 |
| 266 | Q2HJH4 | HNRPDL protein (Fragment) OS=Bos taurus OX=9913 GN=HNRPDL PE=2 SV=1 - [Q2HJH4_BOVIN] | 43612850,28 | 0 | 0 | 145089243,3 | 215326110,2 | HNRPDL | 14537617 | 180207677 | 12,39595731 | 1,093280072 |
| 267 | F1MI47 | RNA-binding protein 14 OS=Bos taurus OX=9913 GN=RBM14 PE=4 SV=1 - [F1MI47_BOVIN] | 0 | 18139425,88 | 0 | 116015448,8 | 23520894,41 | RBM14 | 6046475,3 | 69768171,6 | 11,53865157 | 1,062155059 |
| 268 | P53712 | Integrin beta-1 OS=Bos taurus OX=9913 GN=ITGB1 PE=1 SV=3 - [ITB1_BOVIN] | 80155968,99 | 41704926,55 | 77228971,63 | 923237890,1 | 608074785,6 | ITGB1 | 66363289 | 765656338 | 11,53734766 | 1,06210598 |
| 269 | A5PKE2 | J homolog subfamily C member 10 OS=Bos taurus OX=9913 GN=DNAJC10 PE=2 SV=1 - [A5PKE2_BO | 0 | 5677757 | 4686253,75 | 57307119,65 | 21783545,02 | DNAJC10 | 3454670,3 | 39545332,3 | 11,44691952 | 1,05868863 |
| 270 | F1MBA3 | Integrin alpha-2 OS=Bos taurus OX=9913 GN=ITGA2 PE=3 SV=2 - [F1MBA3_BOVIN] | 15313465,34 | 0 | 24168847,38 | 212114469 | 69053611,63 | ITGA2 | 13160771 | 140584040 | 10,68205209 | 1,028654691 |
| 271 | F1MVW5 | rotein transport protein SEC23 OS=Bos taurus OX=9913 GN=SEC23A PE=3 SV=2 - [F1MVW5_BOVIN | 30385509,44 | 0 | 0 | 65424057,4 | 150906926,3 | SEC23A | 10128503 | 108165492 | 10,67931661 | 1,028543462 |
| 272 | A4FUC2 | HNRNPUL1 protein OS=Bos taurus OX=9913 GN=HNRNPUL1 PE=1 SV=1 - [A4FUC2_BOVIN] | 62446778,17 | 68104453,67 | 0 | 628140830,4 | 298880322,6 | HNRNPUL1 | 43517077 | 463510576 | 10,65123408 | 1,027399929 |
| 273 | P22439 | ate dehydrogenase protein X component OS=Bos taurus OX=9913 GN=PDHX PE=1 SV=3 - [ODPX_B( | 3101652,305 | 0 | 0 | 10946863,5 | 10956165,03 | PDHX | 1033884,1 | 10951514,3 | 10,59259374 | 1,025002316 |
| 274 | 0A140T86 | RNA-binding protein FUS OS=Bos taurus OX=9913 GN=FUS PE=4 SV=1 - [A0A140T861_BOVIN] | 908774748,9 | 0 | 555923732,7 | 8707052535 | 1597151600 | FUS | 488232827 | 5152102067 | 10,55255153 | 1,023357482 |
| 275 | Q5E9J1 | geneous nuclear ribonucleoprotein F OS=Bos taurus OX=9913 GN=HNRNPF PE=2 SV=3 - [HNRPF_B | 234328262,4 | 0 | 0 | 762534311,3 | 849791190 | HNRNPF | 78109421 | 806162751 | 10,32094135 | 1,01371931 |
| 276 | Q3SX36 | TFG protein OS=Bos taurus OX=9913 GN=TFG PE=2 SV=1 - [Q3SX36_BOVIN] | 742470520,9 | 185934071,6 | 80835871,81 | 5154822953 | 1706622608 | TFG | 336413488 | 3430722781 | 10,19793469 | 1,008512226 |
| 277 | Q29RQ2 | nain containing, signal transduction associated 1 OS=Bos taurus OX=9913 GN=KHDRBS1 PE=2 SV=1 | 30713823,88 | 0 | 0 | 63630268,39 | 137359898,1 | KHDRBS1 | 10237941 | 100495083 | 9,81594643 | 0,99193218 |
| 278 | A5D9B4 | eneous nuclear ribonucleoprotein H2 OS=Bos taurus OX=9913 GN=HNRNPH2 PE=2 SV=1 - [A5D9B4 | 377409529,8 | 36343305,75 | 34807281,5 | 1984309501 | 881746133,7 | HNRNPH2 | 149520039 | 1433027817 | 9,584185682 | 0,981555219 |
| 279 | E1BF20 | eneous nuclear ribonucleoprotein H1 OS=Bos taurus OX=9913 GN=HNRNPH1 PE=1 SV=2 - [E1BF20_ | 377409529,8 | 36343305,75 | 102237708,1 | 2199636895 | 1050757610 | HNRNPH1 | 171996848 | 1625197253 | 9,4489944 | 0,975385592 |
| 280 | E1BH80 | methyladenosine RNA binding protein 3 OS=Bos taurus OX=9913 GN=YTHDF3 PE=4 SV=2 - [E1BH80 | 0 | 0 | 7659923,516 | 9070706,848 | 34819009,84 | YTHDF3 | 2553307,8 | 21944858,3 | 8,594677858 | 0,934229603 |
| 281 | F1N1A9 | Integrin alpha-V OS=Bos taurus OX=9913 GN=ITGAV PE=3 SV=2 - [F1N1A9_BOVIN] | 6603196 | 0 | 0 | 22268613,41 | 14774031,39 | ITGAV | 2201065,3 | 18521322,4 | 8,414708149 | 0,925039058 |
| 282 | F1MY06 | inucleotide repeat containing 6B OS=Bos taurus OX=9913 GN=TNRC6B PE=4 SV=2 - [F1MY06_BOVI | 9599428,625 | 0 | 13785247,51 | 85664730,46 | 42595270,75 | TNRC6B | 7794892 | 64130000,6 | 8,227182652 | 0,915251139 |
| 283 | E1BB17 | eneous nuclear ribonucleoprotein H3 OS=Bos taurus OX=9913 GN=HNRNPH3 PE=1 SV=1 - [E1BB17_ | 89448720 | 0 | 14434277,95 | 375924637,9 | 100390389,7 | HNRNPH3 | 34627666 | 238157514 | 6,877665792 | 0,837441068 |
| 284 | Q3SZN2 | Protein transport protein Sec23B OS=Bos taurus OX=9913 GN=SEC23B PE=2 SV=1 - [SC23B_BOVIN] | 25257948,88 | 0 | 0 | 21363720,85 | 92205642,67 | SEC23B | 8419316,3 | 56784681,8 | 6,744571625 | 0,828954371 |
| 285 | Q0VCX2 | Endoplasmic reticulum chaperone BiP OS=Bos taurus OX=9913 GN=HSPA5 PE=2 SV=1 - [BIP_BOVIN | 20508060,84 | 0 | 29759496,53 | 150834543,5 | 57611856,81 | HSPA5 | 16755852 | 104223200 | 6,220107298 | 0,793797876 |
| 286 | E1BIU0 | molog C, COPII coat complex component OS=Bos taurus OX=9913 GN=SEC24C PE=1 SV=1 - [E1BIU0 | 12785460,13 | 11979179,19 | 11057865,27 | 78284627,4 | 62331503,56 | SEC24C | 11940835 | 70308065,5 | 5,888035996 | 0,769970456 |
| 287 | 0A140T89 | Serum albumin OS=Bos taurus OX=9913 GN=ALB PE=4 SV=1 - [A0A140T897_BOVIN] | 154345486,3 | 0 | 34532674,84 | 541872648 | 196991870,7 | ALB | 62959387 | 369432259 | 5,867786786 | 0,768474325 |
| 288 | Q2HJ81 | Tubulin beta-6 chain OS=Bos taurus OX=9913 GN=TUBB6 PE=2 SV=1 - [TBB6_BOVIN] | 45193400 | 0 | 0 | 72861402,14 | 94913710,82 | TUBB6 | 15064467 | 83887556,5 | 5,568571283 | 0,745743783 |
| 289 | F1N6W6 | Sorting nexin OS=Bos taurus OX=9913 GN=SNX9 PE=3 SV=2 - [F1N6W6_BOVIN] | 41817927,13 | 63886600,25 | 5096681,625 | 309419453 | 76955496,83 | SNX9 | 36933736 | 193187475 | 5,230650731 | 0,718555722 |
| 290 | F1MMN6 | Integrin subunit alpha 1 OS=Bos taurus OX=9913 GN=ITGA1 PE=3 SV=2 - [F1MMN6_BOVIN] | 20868443,75 | 0 | 0 | 38860774,34 | 30908186,33 | ITGA1 | 6956147,9 | 34884480,3 | 5,014913535 | 0,700263449 |
| 291 | F1MMS9 | Integrin alpha-3 OS=Bos taurus OX=9913 GN=ITGA3 PE=3 SV=1 - [ITA3_BOVIN] | 29085121,54 | 37321222,65 | 39975131,57 | 300003181,8 | 37622803,15 | ITGA3 | 35460492 | 168812992 | 4,760593645 | 0,677661113 |
| 292 | Q0VBZ6 | oxic granule-associated RNA binding protein OS=Bos taurus OX=9913 GN=TIA1 PE=2 SV=1 - [Q0VBZ | 32589879 | 0 | 68247788,75 | 98617582,25 | 187065675,8 | TIA1 | 33612556 | 142841629 | 4,249650915 | 0,628353257 |
| 293 | A3KMV6 | HNRNPR protein OS=Bos taurus OX=9913 GN=HNRNPR PE=1 SV=1 - [A3KMV6_BOVIN] | 0 | 0 | 17149812,25 | 22975673,28 | 24662022,75 | HNRNPR | 5716604,1 | 23818848 | 4,166607949 | 0,619782638 |
| 294 | G3N0V2 | Keratin 1 OS=Bos taurus OX=9913 GN=KRT1 PE=1 SV=1 - [G3N0V2_BOVIN] | 348896488,4 | 31466160,28 | 16339995,98 | 982605534,3 | 108850315,4 | KRT1 | 132234215 | 545727925 | 4,126979733 | 0,615632336 |
| 295 | K9ZRL4 | hock 70kDa protein 1A OS=Bos indicus x Bos taurus OX=30522 GN=HSPA1A PE=2 SV=1 - [K9ZRL4_B | 34458059,75 | 0 | 29759496,53 | 62441302,5 | 108740268,9 | HSPA1A | 21405852 | 85590785,7 | 3,998475993 | 0,601894493 |
| 296 | Q2HJ60 | eous nuclear ribonucleoproteins A2/B1 OS=Bos taurus OX=9913 GN=HNRNPA2B1 PE=2 SV=1 - [RO/ | 300449360 | 5764215,813 | 166484685,2 | 762078954,3 | 453427315,3 | HNRNPA2B1 | 157566087 | 607753135 | 3,85713161 | 0,586264458 |
| 297 | E1BGX1 | box binding protein associated factor 15 OS=Bos taurus OX=9913 GN=TAF15 PE=4 SV=2 - [E1BGX1_I | 270601926,5 | 5179309,844 | 555923732,7 | 1231238215 | 671032792,4 | TAF15 | 277234990 | 951135504 | 3,430791707 | 0,535394352 |
| 298 | F2Z4H8 | c granule associated RNA binding protein like 1 OS=Bos taurus OX=9913 GN=TIAL1 PE=4 SV=2 - [F2Z | 32589879 | 0 | 61968810,38 | 32794070,73 | 165834341,8 | TIAL1 | 31519563 | 99314206,3 | 3,150875089 | 0,498431187 |
| 299 | Q3SZ32 | Elongin B OS=Bos taurus OX=9913 GN=TCEB2 PE=2 SV=1 - [Q3SZ32_BOVIN] | 14737899,5 | 0 | 0 | 8446719,375 | 3242776,34 | TCEB2 | 4912633,2 | 5844747,86 | 1,189738305 | 0,075451444 |
| 300 | A4IFP2 | KRT4 protein OS=Bos taurus OX=9913 GN=KRT4 PE=2 SV=1 - [A4IFP2_BOVIN] | 530729784,1 | 43708715 | 0 | 231319613,6 | 103214599 | KRT4 | 191479500 | 167267106 | -1,144752868 | -0,05871174 |
| 301 | M0QVZ6 | Keratin, type II cytoskeletal 5 OS=Bos taurus OX=9913 GN=KRT5 PE=3 SV=1 - [M0QVZ6_BOVIN] | 1294734587 | 26098394,97 | 28301463,13 | 132447740,4 | 424797256 | KRT5 | 449711482 | 278622498 | -1,614053009 | -0,207917794 |
| 302 | A6QNZ7 | olytic hyperkeratosis; keratosis palmaris et plantaris) OS=Bos taurus OX=9913 GN=KRT10 PE=2 SV=: | 567143112 | 514488840,7 | 22698209,48 | 351977563,2 | 54400752,08 | KRT10 | 368110054 | 203189158 | -1,811661893 | -0,258077149 |
| 303 | F1MC11 | Keratin, type I cytoskeletal 14 OS=Bos taurus OX=9913 GN=KRT14 PE=1 SV=2 - [F1MC11_BOVIN] | 411133564,8 | 63664366,8 | 293854319,8 | 92332942,21 | 27467909,78 | KRT14 | 256217417 | 59900426 | -4,277388898 | -0,631178738 |

Note: Integrin subunits

Proteins with fold change (Amot IP substrates Ctrl IP) less than 0, which were exclueded from the analysis

### Table S2

| Term | Overlap | P-value | Adjusted P-value | Log10 adjusted P-vaue | Genes |
|---|---|---|---|---|---|
| Cytoskeletal regulation by Rho GTPase_Homo sapiens_P00016 | 9/70 | 7,948E-07 | 2,54336E-05 | 4,594592163 | VASP;ACTA1;TUBB6;TUBB2A;MYH14;MYH9;TUBB4B;MYH10;ACTB |
| Integrin signalling pathway_Homo sapiens_P00034 | 11/156 | 1,90226E-05 | 0,000304361 | 3,516610998 | VASP;ITGB1;ITGA3;ITGA2;ITGA1;FN1;FLNA;ITGAV;LAMC1;ITGA5;ACTB |
| Parkinson disease_Homo sapiens_P00049 | 7/81 | 0,000181887 | 0,001940124 | 2,712170422 | HSPA9;HSPA8;HSPA5;YWHAQ;YWHAZ;YWHAH;HSPA1A |
| Nicotinic acetylcholine receptor signaling pathway_Homo sapiens_P00044 | 6/68 | 0,000471379 | 0,003771034 | 2,423539552 | ACTA1;MYO1E;MYH14;MYH9;MYH10;ACTB |
| Apoptosis signaling pathway_Homo sapiens_P00006 | 5/102 | 0,01693816 | 0,094896565 | 1,022749509 | HSPA8;BAG4;HSPA5;HSPA1A;NFKB2 |
| Inflammation mediated by chemokine and cytokine signaling pathway_Homo sapiens_P00031 | 7/188 | 0,020758624 | 0,094896565 | 1,022749509 | ITGB1;ACTA1;MYH14;MYH9;MYH10;ACTB;NFKB2 |
| Blood coagulation_Homo sapiens_P00011 | 3/38 | 0,01782967 | 0,094896565 | 1,022749509 | SERPINC1;SERPINE1;F2 |
| CCKR signaling map ST_Homo sapiens_P06959 | 6/165 | 0,034421897 | 0,126208838 | 0,898910233 | ITGB1;SERPINE1;RPS6;HSPB1;ITGAV;CLU |
| Huntington disease_Homo sapiens_P00029 | 5/124 | 0,035496236 | 0,126208838 | 0,898910233 | ACTA1;TUBB6;TUBB2A;TUBB4B;ACTB |
| Dopamine receptor mediated signaling pathway_Homo sapiens_P05912 | 2/52 | 0,175863725 | 0,401974228 | 0,39580179 | PPP1CC;FLNA |
| EGF receptor signaling pathway_Homo sapiens_P00018 | 3/109 | 0,213337208 | 0,426674416 | 0,369903397 | YWHAQ;YWHAZ;YWHAH |
| p53 pathway_Homo sapiens_P00059 | 2/71 | 0,27780961 | 0,493883752 | 0,306375262 | SERPINE1;THBS1 |
| FGF signaling pathway_Homo sapiens_P00021 | 2/99 | 0,425218704 | 0,566958271 | 0,246448904 | YWHAQ;YWHAZ |
| Alzheimer disease-presenilin pathway_Homo sapiens_P00004 | 2/99 | 0,425218704 | 0,566958271 | 0,246448904 | ACTA1;ACTB |
| VEGF signaling pathway_Homo sapiens_P00056 | 1/54 | 0,54854247 | 0,649236528 | 0,187597053 | HSPB1 |
| Cadherin signaling pathway_Homo sapiens_P00012 | 2/150 | 0,646221165 | 0,702837194 | 0,153145264 | ACTA1;ACTB |
| T cell activation_Homo sapiens_P00053 | 1/73 | 0,65890987 | 0,702837194 | 0,153145264 | NFKB2 |
| Angiogenesis_Homo sapiens_P00005 | 1/142 | 0,877044778 | 0,905336545 | 0,043189949 | HSPB1 |
| Wnt signaling pathway_Homo sapiens_P00057 | 1/278 | 0,983715036 | 0,983715036 | 0,00713069 | ACTA1 |

**299 proteins listed in Table S1**
**Top 20 enrtiched pathways**
Statistical information
http://amp.pharm.mssm.edu/Enrichr

### Table S3

| | Original readout | | | | | | | | | | | | |
|---|---|---|---|---|---|---|---|---|---|---|---|---|---|
| **Protein** | **Ctrl -Biotin** | | **p80 -Biotin** | | **p130 -Biotin** | | **Ctrl +Biotin** | | **p80 +Biotin** | | **p130 +Biotin** | | **Log10 (fold change)** |
| Amot | 0,000E0 | 0,000E0 | 0,000E0 | 0,000E0 | 0,000E0 | 0,000E0 | 0,000E0 | 0,000E0 | 9,416E7 | 4,498E7 | 4,722E8 | 3,525E8 | 8,61530 |
| Tjp1 | 0,000E0 | 0,000E0 | 0,000E0 | 0,000E0 | 0,000E0 | 0,000E0 | 0,000E0 | 0,000E0 | 1,069E8 | 7,358E7 | 1,346E9 | 1,572E9 | 9,16402 |
| Afdn | 0,000E0 | 0,000E0 | 0,000E0 | 0,000E0 | 0,000E0 | 0,000E0 | 0,000E0 | 0,000E0 | 1,803E7 | 2,910E7 | 2,590E8 | 3,108E8 | 8,45472 |
| Jcad | 0,000E0 | 0,000E0 | 0,000E0 | 0,000E0 | 0,000E0 | 0,000E0 | 0,000E0 | 0,000E0 | 1,540E7 | 1,345E7 | 1,475E8 | 2,498E8 | 8,29810 |
| Shroom4 | 0,000E0 | 0,000E0 | 0,000E0 | 0,000E0 | 0,000E0 | 0,000E0 | 0,000E0 | 0,000E0 | 7,815E7 | 4,982E7 | 1,518E8 | 1,881E8 | 8,23040 |
| Lpp | 0,000E0 | 0,000E0 | 0,000E0 | 0,000E0 | 0,000E0 | 0,000E0 | 0,000E0 | 0,000E0 | 5,242E6 | 6,804E6 | 2,746E7 | 1,037E8 | 7,81665 |
| Tjp2 | 0,000E0 | 0,000E0 | 0,000E0 | 0,000E0 | 0,000E0 | 0,000E0 | 0,000E0 | 0,000E0 | 6,473E6 | 4,853E6 | 3,019E7 | 9,357E7 | 7,79154 |
| Magi1 | 0,000E0 | 0,000E0 | 0,000E0 | 0,000E0 | 0,000E0 | 0,000E0 | 0,000E0 | 0,000E0 | 0,000E0 | 0,000E0 | 5,558E7 | 4,671E7 | 7,70881 |
| Shroom2 | 0,000E0 | 0,000E0 | 0,000E0 | 0,000E0 | 0,000E0 | 0,000E0 | 0,000E0 | 0,000E0 | 7,310E6 | 1,320E7 | 5,022E7 | 3,776E7 | 7,64336 |
| Cgnl1 | 0,000E0 | 0,000E0 | 0,000E0 | 0,000E0 | 0,000E0 | 0,000E0 | 0,000E0 | 0,000E0 | 5,471E6 | 1,290E6 | 4,686E7 | 2,561E7 | 7,55910 |
| Synpo | 0,000E0 | 0,000E0 | 0,000E0 | 0,000E0 | 0,000E0 | 0,000E0 | 0,000E0 | 0,000E0 | 0,000E0 | 0,000E0 | 3,450E7 | 3,592E7 | 7,54668 |
| Amotl1 | 0,000E0 | 0,000E0 | 0,000E0 | 0,000E0 | 0,000E0 | 0,000E0 | 0,000E0 | 0,000E0 | 3,914E7 | 3,990E7 | 1,654E7 | 1,887E7 | 7,24812 |
| Mpdz | 0,000E0 | 0,000E0 | 0,000E0 | 0,000E0 | 0,000E0 | 0,000E0 | 0,000E0 | 0,000E0 | 1,008E7 | 1,129E7 | 1,336E7 | 1,857E7 | 7,20313 |
| Yap1 | 0,000E0 | 0,000E0 | 0,000E0 | 0,000E0 | 0,000E0 | 0,000E0 | 0,000E0 | 0,000E0 | 6,568E6 | 6,665E6 | 9,906E6 | 2,096E7 | 7,18846 |
| Crkl | 0,000E0 | 0,000E0 | 0,000E0 | 0,000E0 | 0,000E0 | 0,000E0 | 0,000E0 | 0,000E0 | 0,000E0 | 7,002E6 | 2,910E7 | 8,951E5 | 7,17608 |
| Sipa1 | 0,000E0 | 0,000E0 | 0,000E0 | 0,000E0 | 0,000E0 | 0,000E0 | 0,000E0 | 0,000E0 | 2,996E6 | 0,000E0 | 1,372E7 | 1,494E7 | 7,15622 |
| Tp53bp2 | 0,000E0 | 0,000E0 | 0,000E0 | 0,000E0 | 0,000E0 | 0,000E0 | 0,000E0 | 0,000E0 | 2,075E7 | 6,426E6 | 1,561E7 | 1,301E7 | 7,15564 |
| Ly6c1 | 0,000E0 | 0,000E0 | 0,000E0 | 0,000E0 | 0,000E0 | 0,000E0 | 0,000E0 | 0,000E0 | 3,000E7 | 1,403E7 | 5,471E6 | 2,284E7 | 7,15088 |
| Pdlim7 | 0,000E0 | 0,000E0 | 0,000E0 | 0,000E0 | 0,000E0 | 0,000E0 | 0,000E0 | 0,000E0 | 0,000E0 | 0,000E0 | 8,944E6 | 1,795E7 | 7,12867 |
| Pdlim5 | 0,000E0 | 0,000E0 | 0,000E0 | 0,000E0 | 0,000E0 | 0,000E0 | 0,000E0 | 0,000E0 | 0,000E0 | 0,000E0 | 1,696E7 | 7,680E6 | 7,09062 |
| Zyx | 0,000E0 | 0,000E0 | 0,000E0 | 0,000E0 | 0,000E0 | 0,000E0 | 0,000E0 | 0,000E0 | 0,000E0 | 0,000E0 | 7,413E6 | 1,525E7 | 7,05431 |
| Cttn | 0,000E0 | 0,000E0 | 0,000E0 | 0,000E0 | 0,000E0 | 0,000E0 | 0,000E0 | 0,000E0 | 9,298E6 | 7,480E6 | 1,827E7 | 3,848E6 | 7,04379 |
| Kif14 | 0,000E0 | 0,000E0 | 0,000E0 | 0,000E0 | 0,000E0 | 0,000E0 | 0,000E0 | 0,000E0 | 0,000E0 | 1,475E6 | 7,520E6 | 1,352E7 | 7,02200 |
| Rassf8 | 0,000E0 | 0,000E0 | 0,000E0 | 0,000E0 | 0,000E0 | 0,000E0 | 0,000E0 | 0,000E0 | 4,587E6 | 0,000E0 | 1,132E7 | 6,952E6 | 6,96076 |
| Dlg5 | 0,000E0 | 0,000E0 | 0,000E0 | 0,000E0 | 0,000E0 | 0,000E0 | 0,000E0 | 0,000E0 | 3,285E6 | 1,196E6 | 3,201E6 | 1,312E7 | 6,91171 |
| Ctnnd1 | 0,000E0 | 0,000E0 | 0,000E0 | 0,000E0 | 0,000E0 | 0,000E0 | 0,000E0 | 0,000E0 | 0,000E0 | 8,505E5 | 9,953E6 | 5,037E6 | 6,87478 |
| Flna | 0,000E0 | 0,000E0 | 0,000E0 | 0,000E0 | 0,000E0 | 0,000E0 | 0,000E0 | 0,000E0 | 3,235E6 | 3,604E6 | 8,537E6 | 5,856E6 | 6,85713 |
| Sorbs2 | 0,000E0 | 0,000E0 | 0,000E0 | 0,000E0 | 0,000E0 | 0,000E0 | 0,000E0 | 0,000E0 | 0,000E0 | 0,000E0 | 5,034E6 | 8,404E6 | 6,82731 |
| Gars | 0,000E0 | 0,000E0 | 0,000E0 | 0,000E0 | 0,000E0 | 0,000E0 | 0,000E0 | 0,000E0 | 4,338E6 | 3,546E6 | 8,423E6 | 4,998E6 | 6,82675 |
| Fbl | 0,000E0 | 0,000E0 | 0,000E0 | 0,000E0 | 0,000E0 | 0,000E0 | 0,000E0 | 0,000E0 | 8,468E6 | 2,231E6 | 3,599E6 | 9,606E6 | 6,81972 |
| Swap70 | 0,000E0 | 0,000E0 | 0,000E0 | 0,000E0 | 0,000E0 | 0,000E0 | 0,000E0 | 0,000E0 | 9,452E6 | 3,962E6 | 6,543E6 | 5,429E6 | 6,77717 |
| Ccdc85c | 0,000E0 | 0,000E0 | 0,000E0 | 0,000E0 | 0,000E0 | 0,000E0 | 0,000E0 | 0,000E0 | 0,000E0 | 0,000E0 | 6,466E6 | 4,970E6 | 6,75722 |
| Flnb | 0,000E0 | 0,000E0 | 0,000E0 | 0,000E0 | 0,000E0 | 0,000E0 | 0,000E0 | 0,000E0 | 5,100E6 | 4,609E6 | 6,327E6 | 4,587E6 | 6,73696 |
| Copa | 0,000E0 | 0,000E0 | 0,000E0 | 0,000E0 | 0,000E0 | 0,000E0 | 0,000E0 | 0,000E0 | 2,440E6 | 2,543E6 | 4,164E6 | 5,347E6 | 6,67721 |
| Tln1 | 0,000E0 | 0,000E0 | 0,000E0 | 0,000E0 | 0,000E0 | 0,000E0 | 0,000E0 | 0,000E0 | 3,006E6 | 1,186E6 | 4,176E6 | 5,283E6 | 6,67482 |
| Baz1b | 0,000E0 | 0,000E0 | 0,000E0 | 0,000E0 | 0,000E0 | 0,000E0 | 0,000E0 | 0,000E0 | 0,000E0 | 0,000E0 | 3,559E6 | 5,449E6 | 6,65356 |
| Cse1l | 0,000E0 | 0,000E0 | 0,000E0 | 0,000E0 | 0,000E0 | 0,000E0 | 0,000E0 | 0,000E0 | 2,415E6 | 1,656E6 | 5,505E6 | 3,454E6 | 6,65123 |
| Micall2 | 0,000E0 | 0,000E0 | 0,000E0 | 0,000E0 | 0,000E0 | 0,000E0 | 0,000E0 | 0,000E0 | 0,000E0 | 0,000E0 | 5,400E6 | 2,932E6 | 6,61970 |
| Crk | 0,000E0 | 0,000E0 | 0,000E0 | 0,000E0 | 0,000E0 | 0,000E0 | 0,000E0 | 0,000E0 | 1,353E6 | 3,142E6 | 7,801E5 | 7,383E6 | 6,61083 |
| Actr2 | 0,000E0 | 0,000E0 | 0,000E0 | 0,000E0 | 0,000E0 | 0,000E0 | 0,000E0 | 0,000E0 | 1,058E6 | 4,738E6 | 2,992E6 | 5,076E6 | 6,60573 |
| Wwc2 | 0,000E0 | 0,000E0 | 0,000E0 | 0,000E0 | 0,000E0 | 0,000E0 | 0,000E0 | 0,000E0 | 1,941E6 | 0,000E0 | 5,451E6 | 2,528E6 | 6,60089 |
| Dvl2 | 0,000E0 | 0,000E0 | 0,000E0 | 0,000E0 | 0,000E0 | 0,000E0 | 0,000E0 | 0,000E0 | 3,992E6 | 0,000E0 | 2,747E6 | 5,142E6 | 6,59601 |
| Nckap1 | 0,000E0 | 0,000E0 | 0,000E0 | 0,000E0 | 0,000E0 | 0,000E0 | 0,000E0 | 0,000E0 | 1,508E6 | 5,671E5 | 2,553E6 | 5,163E6 | 6,58636 |
| Pard3 | 0,000E0 | 0,000E0 | 0,000E0 | 0,000E0 | 0,000E0 | 0,000E0 | 0,000E0 | 0,000E0 | 0,000E0 | 0,000E0 | 2,118E6 | 5,223E6 | 6,56469 |
| Ddx39a | 0,000E0 | 0,000E0 | 0,000E0 | 0,000E0 | 0,000E0 | 0,000E0 | 0,000E0 | 0,000E0 | 1,205E7 | 3,246E6 | 4,607E6 | 2,665E6 | 6,56064 |
| Atp1a1 | 0,000E0 | 0,000E0 | 0,000E0 | 0,000E0 | 0,000E0 | 0,000E0 | 0,000E0 | 0,000E0 | 7,579E5 | 0,000E0 | 2,867E6 | 4,078E6 | 6,54066 |
| Akap2 | 0,000E0 | 0,000E0 | 0,000E0 | 0,000E0 | 0,000E0 | 0,000E0 | 0,000E0 | 0,000E0 | 2,148E6 | 5,235E5 | 1,704E6 | 5,055E6 | 6,52888 |
| Rps24 | 0,000E0 | 0,000E0 | 0,000E0 | 0,000E0 | 0,000E0 | 0,000E0 | 0,000E0 | 0,000E0 | 0,000E0 | 7,616E6 | 4,814E6 | 1,817E6 | 6,52056 |
| Tpm3 | 0,000E0 | 0,000E0 | 0,000E0 | 0,000E0 | 0,000E0 | 0,000E0 | 0,000E0 | 0,000E0 | 0,000E0 | 2,739E6 | 3,329E6 | 2,736E6 | 6,48180 |
| Fxr2 | 0,000E0 | 0,000E0 | 0,000E0 | 0,000E0 | 0,000E0 | 0,000E0 | 0,000E0 | 0,000E0 | 0,000E0 | 0,000E0 | 9,801E5 | 4,737E6 | 6,45618 |
| Kank2 | 0,000E0 | 0,000E0 | 0,000E0 | 0,000E0 | 0,000E0 | 0,000E0 | 0,000E0 | 0,000E0 | 7,248E5 | 0,000E0 | 3,607E6 | 1,650E6 | 6,41968 |
| Parp1 | 0,000E0 | 0,000E0 | 0,000E0 | 0,000E0 | 0,000E0 | 0,000E0 | 0,000E0 | 0,000E0 | 0,000E0 | 0,000E0 | 1,131E6 | 4,096E6 | 6,41724 |
| Dync1h1 | 0,000E0 | 0,000E0 | 0,000E0 | 0,000E0 | 0,000E0 | 0,000E0 | 0,000E0 | 0,000E0 | 3,440E6 | 1,999E6 | 3,107E6 | 1,590E6 | 6,37080 |
| Nf2 | 0,000E0 | 0,000E0 | 0,000E0 | 0,000E0 | 0,000E0 | 0,000E0 | 0,000E0 | 0,000E0 | 4,492E6 | 1,335E7 | 1,142E5 | 4,509E6 | 6,36396 |
| Rpn1 | 0,000E0 | 0,000E0 | 0,000E0 | 0,000E0 | 0,000E0 | 0,000E0 | 0,000E0 | 0,000E0 | 0,000E0 | 0,000E0 | 1,285E6 | 3,189E6 | 6,34960 |
| Nars | 0,000E0 | 0,000E0 | 0,000E0 | 0,000E0 | 0,000E0 | 0,000E0 | 0,000E0 | 0,000E0 | 0,000E0 | 2,485E6 | 2,322E6 | 2,115E6 | 6,34604 |
| Chtop | 0,000E0 | 0,000E0 | 0,000E0 | 0,000E0 | 0,000E0 | 0,000E0 | 0,000E0 | 0,000E0 | 0,000E0 | 0,000E0 | 1,630E6 | 2,701E6 | 6,33548 |
| Ranbp2 | 0,000E0 | 0,000E0 | 0,000E0 | 0,000E0 | 0,000E0 | 0,000E0 | 0,000E0 | 0,000E0 | 3,113E6 | 2,448E6 | 7,706E5 | 3,425E6 | 6,32178 |
| Magi3 | 0,000E0 | 0,000E0 | 0,000E0 | 0,000E0 | 0,000E0 | 0,000E0 | 0,000E0 | 0,000E0 | 0,000E0 | 0,000E0 | 2,208E6 | 1,974E6 | 6,32034 |
| Sptbn1 | 0,000E0 | 0,000E0 | 0,000E0 | 0,000E0 | 0,000E0 | 0,000E0 | 0,000E0 | 0,000E0 | 2,701E6 | 2,132E6 | 2,558E6 | 1,430E6 | 6,29966 |
| Snd1 | 0,000E0 | 0,000E0 | 0,000E0 | 0,000E0 | 0,000E0 | 0,000E0 | 0,000E0 | 0,000E0 | 4,075E6 | 1,826E6 | 2,587E6 | 1,238E6 | 6,28167 |
| Eif3a | 0,000E0 | 0,000E0 | 0,000E0 | 0,000E0 | 0,000E0 | 0,000E0 | 0,000E0 | 0,000E0 | 0,000E0 | 0,000E0 | 6,156E5 | 2,730E6 | 6,22341 |
| Tpx2 | 0,000E0 | 0,000E0 | 0,000E0 | 0,000E0 | 0,000E0 | 0,000E0 | 0,000E0 | 0,000E0 | 0,000E0 | 0,000E0 | 2,223E6 | 1,029E6 | 6,21112 |
| Wdr1 | 0,000E0 | 0,000E0 | 0,000E0 | 0,000E0 | 0,000E0 | 0,000E0 | 0,000E0 | 0,000E0 | 2,509E6 | 3,096E6 | 7,852E5 | 2,430E6 | 6,20614 |
| Asns | 0,000E0 | 0,000E0 | 0,000E0 | 0,000E0 | 0,000E0 | 0,000E0 | 0,000E0 | 0,000E0 | 0,000E0 | 0,000E0 | 1,140E6 | 2,061E6 | 6,20417 |
| Nat10 | 0,000E0 | 0,000E0 | 0,000E0 | 0,000E0 | 0,000E0 | 0,000E0 | 0,000E0 | 0,000E0 | 0,000E0 | 0,000E0 | 8,053E5 | 2,359E6 | 6,19928 |
| Cyfip2 | 0,000E0 | 0,000E0 | 0,000E0 | 0,000E0 | 0,000E0 | 0,000E0 | 0,000E0 | 0,000E0 | 8,012E5 | 1,143E6 | 2,029E5 | 2,587E6 | 6,14461 |
| Flnc | 0,000E0 | 0,000E0 | 0,000E0 | 0,000E0 | 0,000E0 | 0,000E0 | 0,000E0 | 0,000E0 | 1,144E6 | 0,000E0 | 2,339E6 | 1,631E5 | 6,09726 |
| Fau | 0,000E0 | 0,000E0 | 0,000E0 | 0,000E0 | 0,000E0 | 0,000E0 | 0,000E0 | 0,000E0 | 0,000E0 | 0,000E0 | 8,549E5 | 1,102E6 | 5,99049 |
| Pdcd11 | 0,000E0 | 0,000E0 | 0,000E0 | 0,000E0 | 0,000E0 | 0,000E0 | 0,000E0 | 0,000E0 | 0,000E0 | 0,000E0 | 6,227E5 | 8,145E5 | 5,85649 |
| Prpf8 | 0,000E0 | 0,000E0 | 0,000E0 | 0,000E0 | 0,000E0 | 0,000E0 | 0,000E0 | 0,000E0 | 0,000E0 | 0,000E0 | 6,448E5 | 7,501E5 | 5,84351 |
| Actr3 | 0,000E0 | 0,000E0 | 0,000E0 | 0,000E0 | 0,000E0 | 0,000E0 | 0,000E0 | 0,000E0 | 0,000E0 | 0,000E0 | 1,175E5 | 4,395E5 | 5,44483 |
| Eif3l | 0,000E0 | 0,000E0 | 0,000E0 | 0,000E0 | 0,000E0 | 4,616E4 | 0,000E0 | 0,000E0 | 1,646E6 | 2,378E6 | 1,785E6 | 2,744E6 | 2,59380 |
| Dynll1 | 0,000E0 | 0,000E0 | 0,000E0 | 7,622E4 | 0,000E0 | 0,000E0 | 0,000E0 | 0,000E0 | 4,357E6 | 4,878E6 | 3,801E6 | 2,563E6 | 2,52371 |
| Cav1 | 0,000E0 | 0,000E0 | 0,000E0 | 0,000E0 | 1,878E6 | 0,000E0 | 0,000E0 | 0,000E0 | 9,938E6 | 9,421E6 | 2,253E7 | 1,353E7 | 1,88524 |
| Nedd4 | 0,000E0 | 0,000E0 | 0,000E0 | 0,000E0 | 0,000E0 | 3,602E5 | 1,190E6 | 0,000E0 | 2,292E6 | 0,000E0 | 2,632E6 | 1,391E7 | 1,63031 |
| Atp2a2 | 0,000E0 | 0,000E0 | 0,000E0 | 0,000E0 | 0,000E0 | 7,218E5 | 0,000E0 | 0,000E0 | 2,196E6 | 5,478E5 | 3,190E6 | 4,167E6 | 1,61036 |
| H2afz | 4,405E5 | 2,472E7 | 1,692E7 | 8,344E5 | 3,858E7 | 7,069E6 | 5,100E7 | 0,000E0 | 2,122E6 | 1,434E7 | 1,406E8 | 1,178E9 | 1,57735 |
| Smc4 | 0,000E0 | 0,000E0 | 0,000E0 | 0,000E0 | 0,000E0 | 1,864E6 | 0,000E0 | 0,000E0 | 1,353E6 | 4,003E6 | 8,556E6 | 8,100E6 | 1,55315 |
| Abcf2 | 0,000E0 | 0,000E0 | 0,000E0 | 0,000E0 | 0,000E0 | 4,524E5 | 0,000E0 | 0,000E0 | 3,616E6 | 3,094E6 | 1,513E6 | 2,101E6 | 1,50446 |
| Ywhab | 1,928E6 | 0,000E0 | 0,000E0 | 0,000E0 | 0,000E0 | 0,000E0 | 0,000E0 | 0,000E0 | 5,409E6 | 1,717E7 | 1,213E7 | 3,149E6 | 1,50101 |
| Rrbp1 | 0,000E0 | 0,000E0 | 0,000E0 | 7,381E5 | 0,000E0 | 1,064E6 | 0,000E0 | 0,000E0 | 2,696E6 | 1,716E6 | 6,773E6 | 7,426E6 | 1,49858 |
| Slc25a13 | 0,000E0 | 0,000E0 | 0,000E0 | 0,000E0 | 0,000E0 | 0,000E0 | 5,181E5 | 0,000E0 | 0,000E0 | 0,000E0 | 1,546E6 | 2,150E6 | 1,45533 |
| Psat1 | 0,000E0 | 0,000E0 | 0,000E0 | 0,000E0 | 0,000E0 | 0,000E0 | 9,430E5 | 0,000E0 | 3,756E6 | 2,909E6 | 4,454E6 | 1,801E6 | 1,42377 |
| Ywhae | 1,928E6 | 0,000E0 | 0,000E0 | 0,000E0 | 0,000E0 | 0,000E0 | 0,000E0 | 0,000E0 | 4,989E6 | 2,879E6 | 8,585E6 | 3,990E6 | 1,41641 |
| Lbr | 0,000E0 | 0,000E0 | 0,000E0 | 0,000E0 | 0,000E0 | 2,279E6 | 0,000E0 | 0,000E0 | 6,634E6 | 0,000E0 | 2,506E6 | 1,225E7 | 1,41319 |
| Hist1h2ah | 2,149E7 | 1,380E7 | 3,213E7 | 7,339E6 | 2,282E7 | 3,576E7 | 4,141E7 | 0,000E0 | 5,241E7 | 1,996E7 | 1,512E8 | 8,570E8 | 1,36324 |
| Hist2h2ac | 1,795E7 | 1,617E7 | 3,300E7 | 9,178E6 | 1,614E7 | 3,808E7 | 4,607E7 | 0,000E0 | 6,645E7 | 2,035E7 | 1,356E8 | 8,721E8 | 1,35842 |
| Hnrnpf | 0,000E0 | 0,000E0 | 0,000E0 | 0,000E0 | 0,000E0 | 2,688E6 | 0,000E0 | 0,000E0 | 0,000E0 | 0,000E0 | 5,478E6 | 9,737E6 | 1,35488 |
| Actn4 | 1,574E6 | 0,000E0 | 0,000E0 | 0,000E0 | 0,000E0 | 2,131E6 | 1,414E6 | 0,000E0 | 5,572E6 | 2,355E6 | 1,043E7 | 1,846E7 | 1,35353 |
| Myof | 0,000E0 | 0,000E0 | 1,109E6 | 0,000E0 | 5,501E5 | 0,000E0 | 0,000E0 | 0,000E0 | 4,798E6 | 3,975E6 | 4,662E6 | 4,642E6 | 1,35098 |

| | | | | | | | | | | | | | |
|---|---|---|---|---|---|---|---|---|---|---|---|---|---|
| Rpl26 | 0,000E0 | 1,318E7 | 0,000E0 | 0,000E0 | 0,000E0 | 4,210E6 | 0,000E0 | 0,000E0 | 1,007E7 | 1,790E6 | 5,645E7 | 4,020E7 | 1,34707 |
| Alyref | 0,000E0 | 0,000E0 | 0,000E0 | 0,000E0 | 0,000E0 | 1,741E6 | 0,000E0 | 0,000E0 | 0,000E0 | 0,000E0 | 4,263E6 | 5,063E6 | 1,33097 |
| Mki67 | 0,000E0 | 0,000E0 | 0,000E0 | 0,000E0 | 0,000E0 | 1,615E6 | 0,000E0 | 0,000E0 | 0,000E0 | 0,000E0 | 2,836E6 | 5,557E6 | 1,31787 |
| Capn2 | 0,000E0 | 0,000E0 | 0,000E0 | 0,000E0 | 0,000E0 | 2,631E6 | 0,000E0 | 0,000E0 | 0,000E0 | 0,000E0 | 5,836E6 | 7,679E6 | 1,31284 |
| Actn1 | 1,574E6 | 0,000E0 | 0,000E0 | 0,000E0 | 0,000E0 | 3,258E6 | 1,458E6 | 0,000E0 | 3,011E6 | 3,145E6 | 1,333E7 | 1,557E7 | 1,26429 |
| Kcnk3 | 0,000E0 | 0,000E0 | 0,000E0 | 0,000E0 | 1,391E7 | 0,000E0 | 0,000E0 | 0,000E0 | 9,993E6 | 0,000E0 | 2,780E7 | 3,331E7 | 1,24497 |
| Mcm5 | 0,000E0 | 0,000E0 | 0,000E0 | 7,741E5 | 2,409E6 | 0,000E0 | 2,578E6 | 0,000E0 | 5,950E6 | 8,675E6 | 1,395E7 | 1,117E7 | 1,24160 |
| Lrpprc | 0,000E0 | 0,000E0 | 0,000E0 | 0,000E0 | 0,000E0 | 0,000E0 | 2,066E6 | 0,000E0 | 9,282E6 | 5,736E6 | 2,457E6 | 6,188E6 | 1,22366 |
| Coro1b | 0,000E0 | 0,000E0 | 0,000E0 | 0,000E0 | 0,000E0 | 0,000E0 | 5,625E6 | 0,000E0 | 3,063E7 | 7,908E6 | 6,998E6 | 1,569E7 | 1,20776 |
| Rpl23 | 0,000E0 | 0,000E0 | 0,000E0 | 0,000E0 | 0,000E0 | 8,693E5 | 1,540E6 | 0,000E0 | 3,112E6 | 2,196E6 | 5,209E6 | 4,267E6 | 1,19682 |
| Ckap4 | 0,000E0 | 0,000E0 | 0,000E0 | 8,464E5 | 0,000E0 | 0,000E0 | 2,791E6 | 0,000E0 | 3,229E6 | 5,223E6 | 4,849E6 | 9,443E6 | 1,19635 |
| Nop58 | 0,000E0 | 0,000E0 | 0,000E0 | 0,000E0 | 0,000E0 | 2,994E6 | 0,000E0 | 0,000E0 | 2,977E6 | 0,000E0 | 3,106E6 | 7,870E6 | 1,16632 |
| Camk2d | 0,000E0 | 0,000E0 | 0,000E0 | 0,000E0 | 0,000E0 | 0,000E0 | 4,344E5 | 0,000E0 | 0,000E0 | 4,337E6 | 9,658E5 | 5,365E5 | 1,14089 |
| Bub3 | 0,000E0 | 0,000E0 | 0,000E0 | 0,000E0 | 0,000E0 | 0,000E0 | 7,867E5 | 0,000E0 | 0,000E0 | 0,000E0 | 8,859E5 | 1,742E6 | 1,12582 |
| Ddx3y | 0,000E0 | 0,000E0 | 0,000E0 | 0,000E0 | 0,000E0 | 1,365E6 | 7,895E5 | 0,000E0 | 0,000E0 | 2,531E6 | 5,683E6 | 1,496E6 | 1,12479 |
| Fasn | 7,091E5 | 1,037E6 | 0,000E0 | 0,000E0 | 0,000E0 | 0,000E0 | 7,059E5 | 0,000E0 | 1,492E6 | 1,633E6 | 3,665E6 | 4,262E6 | 1,11162 |
| Rps16 | 2,666E6 | 3,349E6 | 2,013E6 | 4,925E6 | 8,053E6 | 1,481E7 | 4,754E5 | 0,000E0 | 4,550E7 | 7,904E6 | 1,893E7 | 9,364E7 | 1,09372 |
| Ddx21 | 5,399E5 | 0,000E0 | 3,352E6 | 1,319E6 | 0,000E0 | 4,019E6 | 1,598E6 | 0,000E0 | 6,244E6 | 2,921E6 | 1,654E7 | 1,620E7 | 1,08254 |
| Psmd2 | 0,000E0 | 0,000E0 | 0,000E0 | 0,000E0 | 2,884E6 | 8,561E5 | 0,000E0 | 0,000E0 | 4,032E6 | 4,651E6 | 5,687E6 | 5,119E6 | 1,06285 |
| Slc25a5 | 5,057E6 | 5,689E6 | 2,100E6 | 3,666E6 | 8,061E5 | 8,691E6 | 3,208E6 | 0,000E0 | 1,906E7 | 4,418E7 | 4,618E7 | 3,665E7 | 1,05459 |
| Rps9 | 3,103E6 | 1,698E6 | 3,781E6 | 1,213E6 | 6,571E5 | 4,158E6 | 6,362E6 | 0,000E0 | 2,831E7 | 3,773E6 | 3,076E7 | 2,852E7 | 1,05332 |
| Tars | 1,512E6 | 0,000E0 | 1,206E6 | 0,000E0 | 0,000E0 | 0,000E0 | 0,000E0 | 0,000E0 | 8,958E5 | 1,824E6 | 2,140E6 | 5,488E6 | 1,05013 |
| Rhoa | 4,031E6 | 1,554E6 | 0,000E0 | 0,000E0 | 1,249E6 | 2,195E6 | 0,000E0 | 0,000E0 | 1,208E7 | 7,005E6 | 6,114E6 | 1,813E7 | 1,03108 |
| Rpl5 | 4,192E6 | 0,000E0 | 0,000E0 | 0,000E0 | 0,000E0 | 2,525E6 | 0,000E0 | 0,000E0 | 1,489E7 | 8,785E6 | 1,103E7 | 5,937E6 | 1,00441 |
| Ddx6 | 2,957E6 | 0,000E0 | 0,000E0 | 0,000E0 | 3,206E6 | 0,000E0 | 0,000E0 | 0,000E0 | 3,461E6 | 3,831E6 | 6,166E6 | 9,339E6 | 1,00270 |
| Slc25a4 | 1,698E6 | 7,351E6 | 0,000E0 | 1,059E6 | 3,352E6 | 1,211E7 | 2,933E6 | 0,000E0 | 2,164E7 | 1,745E7 | 2,958E7 | 4,144E7 | 0,99845 |
| Rps17 | 0,000E0 | 0,000E0 | 0,000E0 | 1,502E6 | 5,282E5 | 0,000E0 | 6,366E5 | 0,000E0 | 3,424E6 | 4,714E6 | 5,012E6 | 1,517E6 | 0,99095 |
| Ywhaq | 1,928E6 | 0,000E0 | 0,000E0 | 0,000E0 | 0,000E0 | 0,000E0 | 2,935E6 | 0,000E0 | 5,409E6 | 1,060E7 | 8,585E6 | 3,149E6 | 0,98461 |
| Psma1 | 0,000E0 | 0,000E0 | 0,000E0 | 0,000E0 | 1,168E6 | 0,000E0 | 1,726E6 | 0,000E0 | 0,000E0 | 2,465E6 | 2,512E6 | 4,362E6 | 0,97780 |
| Rab14 | 0,000E0 | 1,164E6 | 0,000E0 | 5,827E5 | 0,000E0 | 6,900E5 | 0,000E0 | 0,000E0 | 1,635E6 | 0,000E0 | 2,367E6 | 3,416E6 | 0,97735 |
| Rps2 | 4,037E6 | 4,756E6 | 2,043E7 | 1,349E7 | 1,156E7 | 1,293E7 | 1,255E7 | 0,000E0 | 5,062E7 | 3,107E7 | 9,003E7 | 9,675E7 | 0,97162 |
| Rpl34 | 2,459E6 | 0,000E0 | 5,106E6 | 3,934E6 | 1,672E6 | 4,212E6 | 3,776E6 | 0,000E0 | 6,278E6 | 1,986E6 | 6,959E6 | 4,244E7 | 0,97032 |
| Rpl17 | 0,000E0 | 0,000E0 | 0,000E0 | 1,475E6 | 1,217E6 | 1,960E6 | 4,335E6 | 0,000E0 | 3,582E6 | 2,663E6 | 5,460E6 | 1,465E7 | 0,95191 |
| Rpl21 | 0,000E0 | 0,000E0 | 0,000E0 | 0,000E0 | 3,820E6 | 5,534E6 | 1,080E6 | 0,000E0 | 4,707E6 | 3,138E6 | 7,443E6 | 1,543E7 | 0,94291 |
| Top2a | 0,000E0 | 0,000E0 | 0,000E0 | 0,000E0 | 2,732E5 | 1,504E6 | 8,756E5 | 0,000E0 | 1,146E6 | 7,606E5 | 2,503E6 | 3,208E6 | 0,93497 |
| Cfl1 | 8,331E6 | 8,908E6 | 6,183E6 | 4,811E6 | 1,275E6 | 6,316E6 | 2,090E6 | 0,000E0 | 2,102E7 | 1,768E7 | 3,511E7 | 4,298E7 | 0,91585 |
| Rpl12 | 1,655E6 | 0,000E0 | 5,570E6 | 2,483E6 | 6,977E5 | 1,189E6 | 0,000E0 | 0,000E0 | 1,069E7 | 1,309E7 | 1,455E7 | 8,446E6 | 0,89947 |
| Psmd3 | 0,000E0 | 0,000E0 | 3,080E6 | 0,000E0 | 1,784E6 | 4,358E6 | 0,000E0 | 0,000E0 | 7,291E6 | 3,723E6 | 5,636E6 | 1,264E7 | 0,89906 |
| Surf4 | 7,784E5 | 1,001E6 | 0,000E0 | 0,000E0 | 0,000E0 | 0,000E0 | 5,077E5 | 0,000E0 | 3,087E6 | 9,124E5 | 1,119E6 | 3,356E6 | 0,89351 |
| Cltc | 2,599E6 | 6,696E5 | 3,339E5 | 0,000E0 | 1,912E6 | 0,000E0 | 6,301E6 | 0,000E0 | 7,028E6 | 6,893E6 | 1,095E7 | 1,189E7 | 0,88837 |
| Ehd2 | 0,000E0 | 0,000E0 | 0,000E0 | 1,265E6 | 6,861E5 | 2,061E6 | 0,000E0 | 0,000E0 | 4,235E6 | 4,057E6 | 3,761E6 | 3,388E6 | 0,85284 |
| Smarca5 | 0,000E0 | 0,000E0 | 0,000E0 | 1,908E6 | 0,000E0 | 1,669E6 | 0,000E0 | 0,000E0 | 1,766E6 | 1,851E6 | 2,606E6 | 3,663E6 | 0,84583 |
| Rpl7 | 4,098E6 | 1,564E7 | 7,494E6 | 1,192E7 | 5,326E7 | 3,827E7 | 2,016E6 | 0,000E0 | 5,020E7 | 2,981E7 | 9,677E7 | 1,276E8 | 0,83016 |
| Rpl10a | 3,559E6 | 2,532E6 | 6,550E6 | 1,029E7 | 3,247E6 | 1,391E7 | 1,069E7 | 0,000E0 | 2,009E7 | 7,492E6 | 1,821E7 | 6,741E7 | 0,82893 |
| Plec | 4,988E6 | 3,298E5 | 1,104E6 | 2,140E6 | 1,024E6 | 1,638E6 | 1,222E6 | 0,000E0 | 6,372E6 | 4,756E6 | 1,205E7 | 8,915E6 | 0,82859 |
| Myh9 | 6,562E6 | 7,797E6 | 8,951E6 | 6,176E6 | 5,226E6 | 6,976E6 | 6,888E6 | 0,000E0 | 2,237E7 | 3,388E7 | 3,792E7 | 4,332E7 | 0,82541 |
| Slc25a3 | 1,718E6 | 1,202E6 | 1,668E6 | 8,395E5 | 0,000E0 | 0,000E0 | 2,094E6 | 0,000E0 | 5,960E6 | 2,973E6 | 3,063E6 | 9,473E6 | 0,82390 |
| Snu13 | 0,000E0 | 1,892E6 | 0,000E0 | 0,000E0 | 1,895E6 | 4,215E6 | 0,000E0 | 0,000E0 | 2,559E6 | 6,488E6 | 5,855E6 | 7,065E6 | 0,81015 |
| Psma6 | 0,000E0 | 0,000E0 | 0,000E0 | 0,000E0 | 3,395E5 | 1,784E6 | 1,743E6 | 0,000E0 | 0,000E0 | 0,000E0 | 2,405E6 | 3,816E6 | 0,80858 |
| Mybbp1a | 0,000E0 | 1,881E6 | 1,460E6 | 1,856E6 | 1,890E6 | 2,531E6 | 4,522E6 | 0,000E0 | 8,674E6 | 2,215E6 | 8,164E6 | 1,450E7 | 0,80699 |
| Ddx5 | 0,000E0 | 0,000E0 | 1,101E6 | 2,031E6 | 0,000E0 | 0,000E0 | 2,058E6 | 0,000E0 | 6,245E6 | 5,558E6 | 4,494E6 | 3,727E6 | 0,80178 |
| Hspa4 | 0,000E0 | 0,000E0 | 6,709E5 | 0,000E0 | 4,252E5 | 1,505E6 | 0,000E0 | 0,000E0 | 2,200E6 | 0,000E0 | 2,037E6 | 2,081E6 | 0,80168 |
| Syncrip | 0,000E0 | 0,000E0 | 0,000E0 | 1,726E6 | 1,308E6 | 0,000E0 | 9,925E5 | 0,000E0 | 4,590E6 | 3,174E6 | 2,933E6 | 3,219E6 | 0,78610 |
| Rpl19 | 1,875E6 | 1,490E6 | 3,039E6 | 0,000E0 | 4,382E6 | 0,000E0 | 4,284E6 | 0,000E0 | 2,795E6 | 0,000E0 | 1,401E7 | 8,915E6 | 0,78435 |
| Ppid | 4,976E5 | 7,812E5 | 0,000E0 | 0,000E0 | 0,000E0 | 0,000E0 | 1,202E6 | 0,000E0 | 2,642E6 | 0,000E0 | 1,620E6 | 2,069E6 | 0,77435 |
| Hist1h1c | 1,959E6 | 2,169E7 | 9,713E7 | 2,579E7 | 8,749E7 | 9,700E7 | 1,901E7 | 0,000E0 | 3,575E7 | 2,313E7 | 3,105E8 | 1,992E8 | 0,76522 |
| Hspa9 | 0,000E0 | 0,000E0 | 0,000E0 | 0,000E0 | 6,115E5 | 1,276E6 | 1,550E6 | 0,000E0 | 0,000E0 | 3,673E6 | 3,263E6 | 1,736E6 | 0,76465 |
| Npm1 | 0,000E0 | 0,000E0 | 1,345E6 | 1,427E6 | 8,242E5 | 0,000E0 | 0,000E0 | 0,000E0 | 1,956E6 | 4,731E6 | 2,006E6 | 3,205E6 | 0,76320 |
| Top1 | 0,000E0 | 0,000E0 | 8,430E5 | 6,894E5 | 2,714E5 | 1,902E6 | 2,440E6 | 0,000E0 | 6,321E6 | 1,992E6 | 4,922E6 | 3,947E6 | 0,76135 |
| Tagln2 | 0,000E0 | 5,301E6 | 8,567E5 | 3,501E6 | 2,500E6 | 0,000E0 | 2,132E6 | 0,000E0 | 0,000E0 | 3,668E6 | 1,170E7 | 8,542E6 | 0,75333 |
| Rps14 | 1,978E6 | 3,422E6 | 5,826E6 | 2,875E6 | 5,696E6 | 9,422E6 | 6,643E6 | 0,000E0 | 1,235E7 | 9,600E6 | 1,526E7 | 3,533E7 | 0,75153 |
| Rpl6 | 1,023E7 | 9,293E6 | 5,644E6 | 1,491E6 | 4,003E6 | 3,165E7 | 1,694E7 | 0,000E0 | 2,328E7 | 8,804E6 | 3,706E7 | 6,975E7 | 0,73161 |
| Rpl35 | 6,557E6 | 0,000E0 | 0,000E0 | 0,000E0 | 4,319E6 | 9,289E6 | 1,635E6 | 0,000E0 | 5,176E6 | 0,000E0 | 1,332E7 | 1,593E7 | 0,72969 |
| Rps11 | 6,196E6 | 8,758E6 | 1,349E7 | 3,804E6 | 1,022E7 | 2,167E7 | 6,551E6 | 0,000E0 | 1,942E7 | 1,516E7 | 5,527E7 | 3,781E7 | 0,72158 |
| Pcbp2 | 1,089E6 | 0,000E0 | 0,000E0 | 0,000E0 | 1,488E6 | 5,035E6 | 4,872E6 | 0,000E0 | 4,691E6 | 5,634E6 | 6,187E6 | 1,021E7 | 0,72047 |
| Hnrnpu | 1,702E6 | 2,774E6 | 6,167E6 | 4,489E6 | 9,921E6 | 1,761E7 | 5,165E6 | 0,000E0 | 4,496E7 | 3,052E7 | 2,274E7 | 3,899E7 | 0,71284 |
| Pgd | 0,000E0 | 0,000E0 | 1,109E6 | 0,000E0 | 0,000E0 | 1,028E6 | 1,281E6 | 0,000E0 | 4,117E6 | 0,000E0 | 2,346E6 | 1,593E6 | 0,66370 |
| Idh2 | 1,535E6 | 0,000E0 | 2,211E6 | 5,233E5 | 1,080E6 | 3,359E6 | 4,176E6 | 0,000E0 | 4,253E6 | 9,003E6 | 4,751E6 | 9,694E6 | 0,65170 |
| Mccc1 | 1,264E8 | 1,349E8 | 1,545E8 | 3,101E7 | 1,542E8 | 1,770E8 | 3,546E8 | 0,000E0 | 5,494E8 | 4,866E8 | 5,775E8 | 6,873E8 | 0,64997 |
| Vars | 0,000E0 | 0,000E0 | 1,930E6 | 4,014E6 | 1,761E6 | 2,149E6 | 3,742E6 | 0,000E0 | 1,699E7 | 8,116E6 | 5,965E6 | 8,627E6 | 0,63276 |
| Myh10 | 0,000E0 | 5,258E5 | 2,149E6 | 1,432E6 | 1,393E6 | 0,000E0 | 6,842E5 | 0,000E0 | 6,348E6 | 7,126E6 | 3,673E6 | 2,857E6 | 0,62573 |
| Vat1 | 0,000E0 | 7,845E6 | 5,438E6 | 4,114E6 | 0,000E0 | 3,413E6 | 4,410E6 | 0,000E0 | 4,659E6 | 1,602E7 | 1,078E7 | 1,571E7 | 0,62348 |
| Pc | 8,736E8 | 5,102E8 | 3,777E8 | 3,393E8 | 3,723E8 | 9,338E8 | 5,199E8 | 2,128E5 | 2,870E9 | 2,236E9 | 1,639E9 | 2,442E9 | 0,61868 |
| Eef2 | 6,115E6 | 5,515E6 | 5,703E6 | 7,519E6 | 7,618E6 | 1,310E7 | 9,971E6 | 0,000E0 | 1,981E7 | 8,755E6 | 2,880E7 | 2,684E7 | 0,60285 |
| Pcca | 2,006E8 | 1,198E8 | 4,871E7 | 1,378E8 | 7,757E7 | 2,033E8 | 3,560E8 | 0,000E0 | 6,097E8 | 5,039E8 | 4,315E8 | 6,899E8 | 0,59346 |
| Eif4a1 | 1,100E7 | 3,137E7 | 1,636E6 | 6,419E6 | 1,410E7 | 2,823E7 | 1,247E7 | 0,000E0 | 3,655E7 | 1,519E7 | 6,124E7 | 4,019E7 | 0,58608 |
| Taldo1 | 1,979E6 | 0,000E0 | 2,450E6 | 5,257E6 | 0,000E0 | 3,084E6 | 1,381E6 | 0,000E0 | 5,739E6 | 5,136E6 | 7,877E6 | 5,417E6 | 0,57497 |
| Serpinb6 | 1,641E6 | 0,000E0 | 0,000E0 | 0,000E0 | 5,332E6 | 4,262E6 | 1,533E6 | 0,000E0 | 1,848E7 | 6,719E6 | 6,075E6 | 5,906E6 | 0,57441 |
| Crip2 | 0,000E0 | 5,011E6 | 1,532E6 | 6,881E6 | 9,018E5 | 4,907E6 | 0,000E0 | 0,000E0 | 3,562E6 | 3,808E6 | 1,041E7 | 7,356E6 | 0,56765 |
| Cct8 | 3,317E6 | 2,436E6 | 7,735E5 | 3,434E5 | 2,249E6 | 3,819E6 | 4,811E6 | 0,000E0 | 9,072E6 | 1,179E7 | 7,018E6 | 8,294E6 | 0,53790 |
| Esd | 6,142E7 | 2,451E7 | 4,511E7 | 1,713E7 | 3,408E7 | 9,679E7 | 3,175E7 | 0,000E0 | 1,198E8 | 4,140E7 | 1,155E8 | 1,427E8 | 0,52151 |
| Fkbp1a | 5,914E6 | 6,765E6 | 5,150E6 | 2,507E6 | 6,994E6 | 2,373E6 | 1,108E6 | 0,000E0 | 8,598E6 | 6,499E6 | 7,282E6 | 1,647E7 | 0,48905 |
| Rpl10 | 2,930E6 | 4,042E6 | 6,781E5 | 0,000E0 | 2,163E6 | 4,130E6 | 5,536E6 | 0,000E0 | 5,903E6 | 1,559E7 | 7,711E6 | 6,289E6 | 0,45862 |
| Hspa8 | 1,428E7 | 2,550E7 | 2,224E7 | 1,470E7 | 8,789E6 | 1,438E7 | 3,591E7 | 0,000E0 | 6,525E7 | 3,986E7 | 4,905E7 | 4,410E7 | 0,43835 |
| Myo1c | 7,267E6 | 8,078E6 | 9,209E6 | 6,046E6 | 4,695E6 | 1,051E7 | 1,043E7 | 0,000E0 | 1,303E7 | 1,425E7 | 1,890E7 | 1,821E7 | 0,42158 |
| Pkm | 4,386E7 | 5,690E7 | 4,582E7 | 3,158E7 | 4,644E7 | 5,207E7 | 4,168E7 | 0,000E0 | 1,042E8 | 6,132E7 | 1,099E8 | 9,937E7 | 0,41980 |
| Hnrnpa3 | 0,000E0 | 6,370E6 | 4,076E6 | 3,090E6 | 2,454E6 | 5,300E6 | 6,635E6 | 0,000E0 | 3,698E6 | 9,019E6 | 8,620E6 | 8,657E6 | 0,39352 |
| Ehd4 | 2,676E6 | 1,130E6 | 1,913E6 | 2,136E6 | 5,433E5 | 2,672E6 | 2,786E6 | 0,000E0 | 0,000E0 | 2,845E6 | 3,037E6 | 5,306E6 | 0,38174 |

Selection criteria:
1. Both of the duplicates in Amot p130 + Biotin group has positive values.
2. Either of the duplicates in Amot p130 + Biotin group is less than any of the control groups (Ctrl -/+ Biotin, p80/p130 -Biotin).

| | | | | | | | | | | | | |
|---|---|---|---|---|---|---|---|---|---|---|---|---|
| Color code | 1,000E0 | 2,000E0 | 3,000E0 | 4,000E0 | 5,000E0 | 6,000E0 | 7,000E0 | 8,000E0 | 9,000E0 | 1,000E1 | 1,100E1 | 1,200E1 |

### Table S4

| Term | Overlap | P-value | Adjusted P-value | Log10 adjusted P-vaue | Genes |
|---|---|---|---|---|---|
| Integrin signalling pathway Homo sapiens P00034 | 9/156 | 1,16E-05 | 6,51E-04 | 3,186273143 | ACTN1;CAV1;FLNA;FLNB;ACTN4;TLN1;CRK;RHOA;CRKL |
| Parkinson disease Homo sapiens P00049 | 7/81 | 8,14E-06 | 9,12E-04 | 3,040062684 | YWHAE;HSPA9;PSMA6;HSPA8;YWHAQ;PSMA1;YWHAB |
| Pentose phosphate pathway Homo sapiens P02762 | 2/8 | 0,002152737 | 0,060276622 | 1,219851091 | TALDO1;PGD |
| Pyruvate metabolism Homo sapiens P02772 | 2/8 | 0,002152737 | 0,08036883 | 1,094912354 | PKM;PC |
| Huntington disease Homo sapiens P00029 | 5/124 | 0,005190295 | 0,116262598 | 0,934559975 | VAT1;ACTR2;DYNC1H1;CAPN2;DYNLL1 |
| DNA replication Homo sapiens P00017 | 2/19 | 0,01232486 | 0,230064059 | 0,638151223 | TOP2A;TOP1 |
| Nicotine pharmacodynamics pathway Homo sapiens P06587 | 2/28 | 0,025852072 | 0,361929006 | 0,44137661 | FLNA;KCNK3 |
| Cytoskeletal regulation by Rho GTPase Homo sapiens P00016 | 3/70 | 0,024916444 | 0,398663097 | 0,399393963 | CFL1;MYH9;MYH10 |
| Angiogenesis Homo sapiens P00005 | 4/142 | 0,03891336 | 0,484255152 | 0,31492575 | DVL2;CRK;RHOA;CRKL |
| Asparagine and aspartate biosynthesis Homo sapiens P02730 | 1/5 | 0,043960207 | 0,492354318 | 0,307722249 | ASNS |
| Serine glycine biosynthesis Homo sapiens P02776 | 1/6 | 0,052518882 | 0,534737705 | 0,271859193 | PSAT1 |
| FGF signaling pathway Homo sapiens P00021 | 3/99 | 0,059318151 | 0,553636072 | 0,256775621 | YWHAE;YWHAQ;YWHAB |
| Dopamine receptor mediated signaling pathway Homo sapiens P05912 | 2/52 | 0,078984415 | 0,631875319 | 0,199368608 | FLNA;KCNK3 |
| EGF receptor signaling pathway Homo sapiens P00018 | 3/109 | 0,074525034 | 0,642061832 | 0,192423147 | YWHAE;YWHAQ;YWHAB |
| Inflammation mediated by chemokine and cytokine signaling pathway Homo sapiens P00031 | 4/188 | 0,088804017 | 0,663069991 | 0,178440627 | CAMK2D;MYH9;MYH10;RHOA |
| Opioid proopiomelanocortin pathway Homo sapiens P05917 | 1/18 | 0,149468589 | 0,727847044 | 0,137959878 | KCNK3 |
| Glycolysis Homo sapiens P00024 | 1/17 | 0,141781006 | 0,756165367 | 0,121383218 | PKM |
| Opioid proenkephalin pathway Homo sapiens P05915 | 1/18 | 0,149468589 | 0,760931 | 0,118654722 | KCNK3 |
| 5HT4 type receptor mediated signaling pathway Homo sapiens P04376 | 1/16 | 0,13402433 | 0,790038158 | 0,102351932 | KCNK3 |
| Axon guidance mediated by semaphorins Homo sapiens P00007 | 1/17 | 0,141781006 | 0,793973635 | 0,100193919 | RHOA |

**178 proteins (fc>0) listed in Table S3**
**Top 20 enrtiched pathways**
Statistical information
http://amp.pharm.mssm.edu/Enrichr

Table S5

#### Information of human patient samples

**The human tissue samples were collected in agreement with approval from the Research Ethics Committee at Uppsala University (Ups 02-577, #2011/473).**

All images listed in Figure 1 were obtained from Human Atlas Protein database (https://www.proteinatlas.org/search/amot). Official consent was granted.

| | | | | |
|---|---|---|---|---|
| **Endometrium**<br>HPA067290<br><br>Female, age 39<br>Endometrium (T-84000)<br>Normal tissue, NOS (M-00100)<br>Patient id: 4569 | **Placenta**<br>HPA067290<br><br>Female, age 30<br>Placenta (T-88100)<br>Normal tissue, NOS (M-00100)<br>Patient id: 2515 | **Cerebral cortex (Brain)**<br>HPA067290<br><br>Female, age 52<br>Cerebral cortex (T-X2020)<br>Normal tissue, NOS (M-00100)<br>Patient id: 3740 | **Glioma**<br>HPA067290<br><br>Female, age 71<br>Brain (T-X2000)<br>Glioma, malignant, High grade (M-93803[illegible]<br>Patient id: 2811 | **Breast**<br>HPA067290<br><br>Female, age 49<br>Breast (T-04000)<br>Normal tissue, NOS (M-00100)<br>Patient id: 4930 |
| **Breast cancer**<br>HPA067290<br><br>Female, age 30<br>Breast (T-04000)<br>Duct carcinoma (M-85003)<br>Patient id: 4850 | **Pancreas**<br>HPA067290<br><br>Female, age 58<br>Pancreas (T-59000)<br>Normal tissue, NOS (M-00100)<br>Patient id: 5075 | **Pancreatic cancer**<br>HPA067290<br><br>Female, age 58<br>Pancreas (T-59000)<br>Adenocarcinoma, NOS (M-81403)<br>Patient id: 5075 | **Prostate**<br>HPA067290<br><br>Male, age 61<br>Prostate (T-77100)<br>Normal tissue, NOS (M-00100)<br>Patient id: 5474 | **Prostate cancer**<br>HPA067290<br><br>Male, age 84<br>Prostate (T-77100)<br>Adenocarcinoma, High grade (M-814033)<br>Patient id: 2817 |
| **Colon**<br>HPA067290<br><br>Female, age 84<br>Colon (T-67000)<br>Normal tissue, NOS (M-00100)<br>Patient id: 1958 | **Colorectal cancer**<br>HPA067290<br><br>Female, age 86<br>Colon (T-67000)<br>Adenocarcinoma, NOS (M-81403)<br>Patient id: 4094 | **Kidney**<br>HPA067290<br><br>Male, age 61<br>Kidney (T-71000)<br>Normal tissue, NOS (M-00100)<br>Patient id: 1859 | **Epididymis**<br>HPA067290<br><br>Male, age 29<br>Epididymis (T-79100)<br>Normal tissue, NOS (M-00100)<br>Patient id: 5469 | **Carcinoid**<br>HPA067290<br><br>Male, age 41<br>Pancreas (T-59000)<br>Carcinoid, malignant, NOS (M-82403)<br>Patient id: 2618 |
| **Melanoma**<br>HPA067290<br><br>Male, age 41<br>Smooth muscle (T-1X300)<br>Malignant melanoma, Metastatic site (M-[illegible]<br>Patient id: 2246 | **Head and neck cancer**<br>HPA067290<br><br>Female, age 85<br>Head-Neck (T-Y0000)<br>Oral tissue (T-51000)<br>Squamous cell carcinoma, metastatic, N[illegible]<br>Squamous cell carcinoma, NOS (M-80703)<br>Patient id: 4432 | **Cervical cancer**<br>HPA067290<br><br>Female, age 32<br>Cervix (T-83000)<br>Squamous cell carcinoma, NOS (M-80703)<br>Patient id: 2841 | | |

| Name | Company | Catalog number |
|---|---|---|
| *In vivo* | | |
| Tamoxifen | Sigma | T5648 |
| *In vitro* | | |
| WB | | |
| Protease Inhibitor | Roche | 4693159001 |
| SDS sample buffer | Novex | 1225644 |
| Sample reducing agent | Novex | 1176192 |
| Polyacrylamide Bis-Tris 4-12% gradient gel | Novex | NP0322BOX |
| Nitrocellulose membrane | Whatman | 10401396 |
| Western Lightning Plus-ECL | PerkinElmer | 203-170071 |
| IF | | |
| Paraformaldehyde solution 4% in PBS | ChemCruz | sc-281692 |
| Fluoroshield with DAPI | Sigma | F6057 |
| IP | | |
| IgG from rabbit serum | Sigma | I8140 |
| IgG from mouse serum | Sigma | I8765 |
| IgG from rat serum | Sigma | I8015 |
| Protein G Sepharose 4 fast flow | GE Healthcare | 17-0618-01 |
| Cell Culture | | |
| RPMI-1640 | GIBCO | 21875-034 |
| FBS | GIBCO | 10270-106 |
| Penicillin/streptomycin (P/S) | GIBCO | 15140-122 |
| BAE (Bovine Aortic Endothelial Cells) | Sigma | B304-05 |
| BAE medium | Sigma | B211-500 |
| DMEM medium | GIBCO | 41965-039 |
| Lipofectamine RNAiMAX Reagent | Thermo Fisher Scientific | 13778075 |
| Lipofectamine 3000 | Thermo Fisher Scientific | L3000008 |
| Blebbistatin | Sigma | B0560 |
| BioID | | |
| biotin | Sigma | B4501 |
| urea | Sigma | U5378 |

| Name | Company | Sequence if it's customized product |
|---|---|---|
| ON-TARGETplus control pool Non-targeting pool. D-001810-10 | GE Healthcare Dharmacon | - |
| ON-TARGETplus SMARTpool Mouse Amot. L-058986-01 | GE Healthcare Dharmacon | - |
| Bovine control siRNA (Customized product) | GE Healthcare Dharmacon | 5'-AUUGUAUGCGAUCGCAGACUU-3' |
| Bovine Amot siRNA (Customized product) | GE Healthcare Dharmacon | 5'-GGAGAAGGCUAUUCCGCUAUU-3' |
| siGENOME™ Control Pool 1# D-001206-13 | GE Healthcare Dharmacon | - |
| Human Amot siRNA (Customized product) | GE Healthcare Dharmacon | 5'-AUUGUAUUCUGAAACGUUGGU-3' |

**Mouse p130-Amot-GFP plasmid**

Vector ID: VB190414-1114pch

Vector Name: pcDNA3.1(+)-EGFP:{mAmot[NM_153319.3]*}

Date Created (PacificTime): 2019-04-14

Vector Size: 9439 bp

Vector Type: Mammalian Gene Expression Vector

Plasmid Copy Number: High

Antibiotic Resistance: Ampicillin

Cloning Host: Stbl3 (or alternative strain)

**empty vector plasmid**

Vector ID: VB190314-1055uyu

Vector Name: pLV[Exp]-Neo-CMV>Stuffer300

Date Created (PacificTime): 2019-03-13

Vector Size: 8402 bp

Viral Genome Size: 4927 bp

Vector Type: Mammalian Gene Expression Vector

Plasmid Copy Number: High

Antibiotic Resistance: Ampicillin

Cloning Host: Stbl3 (or alternative strain)

**Mouse p130-Amot-BirA plasmid**

Vector ID:VB190414-1109xdu

Vector Name: pLV[Exp]-Neo-CMV>{BirA(R118G)}:{mAmot[NM_153319.3]*}

Date Created (PacificDate Created (PacificTime)Time): 2019-04-14

Vector SizeVector Size: 12443 bp

Viral Genome Size: 8968 bp

Vector TypeVector Type: Mammalian Gene Expression Lentiviral Vector

Inserted Promoter: CMV

Inserted ORF: {BirA(R118G)}, {mAmot[NM_153319.3]*}

Inserted Marker: Neo

Plasmid Copy Number: High

Antibiotic Resistance: Ampicillin

Cloning Host: Stbl3 (or alternative strain)

**Mouse p80-Amot-BirA plasmid**

Vector ID:VB190411-1454njh

Vector Name: pLV[Exp]-Neo-CMV>{BirA(R118G)}:{hAMOT[NM_133265.2](G449D)}

Date Created (PacificDate Created (PacificTime)Time): 2019-04-11

Vector SizeVector Size: 11099 bp

Viral Genome Size: 7624 bp

Vector TypeVector Type: Mammalian Gene Expression Lentiviral Vector

Inserted Promoter: CMV

Inserted ORF: {BirA(R118G)}, {hAMOT[NM_133265.2](G449D)}

Inserted Marker: Neo

Plasmid Copy Number: High

Antibiotic Resistance: Ampicillin

Cloning Host: Stbl3 (or alternative strain)

### Table S6

| Primary Ab | Full name | Conpany | Catalog number | Dilution |
|---|---|---|---|---|
| Isolectin B4 | Biotinylated griffonia simplicifolia lectin 1 | VECTOR | B-1205 | IF 1:300 for retina |
| GFP | Chk pAb to GFP | Abcam | ab13970 | IF 1:200 |
| ERG | Rb mAb to ERG | Abcam | ab92513 | IF 1:200 |
| GFAP | Glial Fibrillary Acidic Protein | Dako | Z0334 | IF 1:300 |
| Fibronectin | Fibronectin Antibody FBN11 | Thermo Scientific | MA5-11981 | IF 1:300 |
| FAK | Rb mAb to FAK EP695Y | Abcam | ab40794 | WB-1:500 |
| Fibronectin | Rb x Ms Fibronectin | Millipore | AB2033 | IF 1:300 WB 1:2000 |
| Paxillin | Purified Mouse Anti-Paxillin mAB | BD | 610620 | IF 1:200 WB 1:500 |
| Kank2 | Anti-KANK2 antibody produced in rabbit | Sigma | HPA015643-100UL | WB 1:500 |
| Zyxin | Recombinant Anti-Zyxin antibody [EPR4302] | Abcam | ab109316 | WB 1:500 |
| Talin | Ms mAb to Talin 1 804 | Abcam | ab157808 | WB 1:500 |
| Myosin | Myosin light chain 2 Rabbit Ab | Cell signalling | 3672S | WB 1:500 IF 1:200 |
| Vinculin | Monoclonal Anti Vinculin | Sigma | V9131-2ml | WB 1:500 |
| β-actin | Ms mAb to Actin | Abcam | ab3284 | WB 1:2000 |
| GAPDH | Anti-GAPDH antibody [6C5] | Abcam | ab8245 | WB 1:2000 |
| Myc Antibody | Myc-Tag (9B11) Mouse mAb | Cell signalling | #2276 | IF 1:4000 WB 1:500 |
| β1 integrin | Integrin beta 1 / CD29 antibody | Genetex | GTX 128839 | WB 1:500 |
| β1 integrin | Anti-β1 integrin mAB | BD | 610467 | WB 1:500, IP:2ug/70 millions cells, IF 1:100 |
| β1 integrin (Active) | Purified Mouse Anti-CD29 | BD | 610468 | WB 1:500, IP:2ug/70 millions cells, IF 1:100 |
| β1 integrin (alpha 4) | Purified Rat anti-Mouse CD29 9EG7 | BD | 553715 | WB 1:500, IP:2ug/70 millions cells, IF 1:100 |
| β1 integrin (alpha 5) | Anti-Integrin α5β1 Antibody, clone BMB5 | Millipore | MAB2514 | IP:2ug/70 millions cells |
| β3 integrin (alpha v) | Anti-Integrin αVβ3 Antibody, clone LM609 | Millipore | MAB1976 | IP:2ug/70 millions cells |
| β5 integrin (alpha v) | Anti-Integrin αVβ5 Antibody, clone P1F6 | Millipore | MAB1961 | IP:2ug/70 millions cells |
| β3 integrin | Anti-Integrin beta 3 antibody | abcam | ab119992 | WB 1:500 |
| CD31 | Purified Rat Anti-Mouse CD31 | BD | 557355 | IF 1:200 |
| NG2 | Anti-NG2 Antibody, clone 132.38 | Sigma | 05-710 | IF 1:200 |
| BirA | Anti-BirA antibody [6C4c7] | abcam | ab232732 | WB 1:500 IF 1:200 |
| Amot | Angiomotin | Innovagen, Lund, Sweden | Purified from rabbit serum | WB-1:500, IP:2ug/70 millions cells, IF/IHC 1:100 |
| AmotL1 | Angiomotin Like 1 | Innovagen, Lund, Sweden | Purified from rabbit serum | WB-1:500, IP:2ug/70 millions cells, IF/IHC 1:100 |
| AmotL2 | Angiomotin Like 2 | Innovagen, Lund, Sweden | Purified from rabbit serum | WB-1:500, IP:2ug/70 millions cells, IF/IHC 1:100 |
| **Reagent** | | | | |
| Phalloidin | Texas Red-X-Phalloidin | Life Technologies | T7471 | IF 1:300 for cell; 1:200 for retina |
| Phalloidin | Phalloidin-Atto 647N | Sigma | 65906-10nmol | IF 1:300 for cell; 1:200 for retina |
| Phalloidin | Phalloidin, Fluorescein Isothiocyanate Labeled | Sigma | P5282 | IF 1:300 for cell |
| TO-PRO-3 | TO-PRO- iodide (642/661) | Life Technologies | T3605 | IF 1:1000 |

| | Secondary Antibody | Conjugation | Catalog number | Conpany | Dilution |
|---|---|---|---|---|---|
| For WB | ECL® Anti-rabbit IgG | HRP linked whole antibody from donkey | NA934V | GE Healthcare | 1:10000 |
| | ECL® Anti-mouse IgG | HRP linked whole antibody from donkey | NA931V | GE Healthcare | 1:10000 |
| | ECL® Anti-rat IgG | HRP linked whole antibody from donkey | NA935V | GE Healthcare | 1:10000 |
| For IF staining | goat anti-Mouse IgG (H+L) | Alexa Fluor® 405 conjugate | A31553 | LifeTechnologies | 1:500 |
| | goat anti-Rat IgG (H+L) | Alexa Fluor® 488 conjugate | A11006 | LifeTechnologies | 1:500 |
| | donkey anti-Sheep IgG (H+L) | Alexa Fluor® 488 conjugate | A11015 | LifeTechnologies | 1:500 |
| | goat anti-Chicken IgG (H+L) | Alexa Fluor® 488 conjugate | A11039 | LifeTechnologies | 1:500 |
| | donkey anti-Goat IgG (H+L) | Alexa Fluor® 488 conjugate | A11055 | LifeTechnologies | 1:500 |
| | chicken anti-Rabbit IgG (H+L) | Alexa Fluor® 488 conjugate | A21441 | LifeTechnologies | 1:500 |
| | donkey anti-Mouse IgG (H+L) | Alexa Fluor® 555 conjugate | A31570 | LifeTechnologies | 1:500 |
| | donkey anti-Rabbit IgG (H+L) | Alexa Fluor® 555 conjugate | A31572 | LifeTechnologies | 1:500 |
| | goat anti-Mouse IgG (H+L) | Alexa Fluor® 594 conjugate | A11005 | LifeTechnologies | 1:500 |
| | goat anti-Rat IgG (H+L) | Alexa Fluor® 594 conjugate | A11007 | LifeTechnologies | 1:500 |
| | goat anti-Rabbit IgG (H+L) | Alexa Fluor® 594 conjugate | A11037 | LifeTechnologies | 1:500 |
| | goat anti-Rat IgG (H+L) | Alexa Fluor® 633 conjugate | A21094 | LifeTechnologies | 1:500 |
| | donkey anti-Mouse IgG (H+L) | Alexa Fluor® 647 conjugate | A31571 | LifeTechnologies | 1:500 |
| | donkey anti-Rabbit IgG (H+L) | Alexa Fluor® 647 conjugate | A31573 | LifeTechnologies | 1:500 |
| | goat anti-Mouse IgG (H+L) | Cy3® | A10521 | LifeTechnologies | 1:500 |
| | goat anti-Rabbit IgG (H+L) | Cy3 | A10520 | LifeTechnologies | 1:500 |
| | Streptavidin | Alexa Fluor® 405 conjugate | S32351 | LifeTechnologies | 1:500 |
| | Streptavidin | Alexa Fluor® 488 conjugate | S32354 | LifeTechnologies | 1:500 |

| Kit | Description | Company | Catalog number |
|---|---|---|---|
| RNeasy Mini Kit | RNA Isolation | QIAGEN | 74104 |
| EndoFree® Plasmid Maxi Kit | Plamid purification | QIAGEN | 12362 |
| CYTOOCHIPS™ STARTER'S A X18 | CYTOO chip | CYTOO | 10-900-00-18 |
| Duolink® In Situ Orange Starter Kit Mouse/Rabbit | PLA kit | Sigma | DUO92102 |
| Lipofectamine 3000 Transfection Reagent | lentiviral production | Invitrogen | L3000015 |
